# A systematic survey of distal element-gene regulatory interactions with Direct-Capture Targeted Perturb-seq

**DOI:** 10.1101/2025.09.16.676677

**Authors:** Judhajeet Ray, Evelyn Jagoda, Maya U. Sheth, James Galante, Dulguun Amgalan, Eugenio Mattei, Andreas R. Gschwind, Chad J. Munger, Jacob Huang, Glen Munson, Madeleine Murphy, Timothy Barry, Vasundhara Singh, Aarthee Baskaran, Helen Kang, Robbyn Issner, Charles B. Epstein, Fadi J. Najm, Eugene Katsevich, Elizabeth Gaskell, Lars M. Steinmetz, Jesse M. Engreitz

## Abstract

Identifying the impact of distal regulatory elements on gene expression is a core challenge in human genetics. Large-scale CRISPR screens have not captured lower effect size element-gene interactions due to selection bias and limited statistical power. We developed a framework for highly powered CRISPR screens, consisting of Direct-Capture Targeted Perturb-seq (DC-TAP-seq), unbiased target selection, and a pipeline accounting for statistical power. Surveying 9,666 random distal element-gene pairs revealed that most elements that reduced gene expression had effect sizes <10%, which prior studies were underpowered to detect. Out of the 84 interactions with negative effects, 45 (53.6%) occurred within 100 Kb of the promoter of the target gene. Many elements were bound by CTCF without classical enhancer chromatin, and both housekeeping and non-housekeeping genes had similar frequencies of distal regulatory elements. We highlight limitations of predictive models and suggest that new models consider elements with smaller effect sizes. Our study provides an expanded view of distal regulatory elements and a framework for building more comprehensive maps of distal regulation.

## Introduction

Gene regulation is coordinated by noncoding elements that quantitatively tune gene expression^1–6^. While massive consortium efforts such as ENCODE, IGVF, IHEC and others have created expansive maps of distal noncoding elements^7–9^, major goals remain, including: connecting distal elements to their target genes, characterizing the properties of such element-gene regulatory interactions, building predictive models to infer their effects across cell types and states, and ultimately interpreting the functions of noncoding variants associated with disease^9–12^.

To this end, we and others have developed high-throughput tools to perturb candidate regulatory elements with CRISPR interference (CRISPRi) and measure the effects on the expression of nearby genes using various single-cell readouts such as whole-transcriptome single-cell RNA-seq (Perturb-seq)^13,14^, targeted Perturb-seq (TAP-seq)^15^, or flow sorting^16–18^. Such CRISPRi tiling approaches have now been applied to study thousands of candidate element-gene regulatory interactions^14–27^. These data, while representing only a small fraction of the genome, have nonetheless provided initial insights about the properties of distal regulatory elements and their target genes. In particular, distal regulatory elements identified in previous studies are frequently marked by H3K27ac; are most often located within 100 kb of their target promoters^16,27^; have effects that depend on physical 3D contacts with a target promoter^16,22,28^; and are found less frequently for housekeeping genes^16^.

These initial CRISPRi datasets have also enabled training and validating predictive models of element-gene regulatory interactions^16,29–32^. For example, we recently used CRISPR perturbation datasets to train a supervised classifier to predict which elements act as enhancers to regulate which nearby genes. This ENCODE-rE2G model performs well at identifying enhancer-gene regulatory interactions in the training CRISPR dataset, and can generalize across cell types to predict the effect of eQTL and GWAS variants^29^. Thus, large-scale CRISPRi tiling datasets can reveal rules of distal regulation that help to build predictive models that generalize across cell types and states.

Yet, existing datasets have biases that skew our view of the regulatory genome and/or limit the scope of what models can predict:

1. *Statistical power.* Previous Perturb-seq studies have found many elements with small effect sizes on gene expression (*e.g.*, 5–25%)^14,15,19,20,22,27^. However, these studies also appear to have low statistical power, and thus may have missed many other real element-gene regulatory interactions with similar effect sizes. Targeted Perturb-seq (TAP-seq)^15^, or other methods to sequence selected genes^33,34^, could help to achieve better-powered measurements, but have limitations in scalability and have not yet been applied to examine noncoding elements with high power.
2. *Selection bias.* Existing large-scale studies have been designed to examine a biased sample of the genome. Some experiments have selected genes of interest (*e.g.*, near transcription factors in K562 cells^16,27,28^, or the beta-globin locus^15,18^) and studied the effects of nearby elements. Other studies have selected candidate elements of interest (*e.g.*, with strong H3K27ac ChIP-seq signal^14^, overlapping noncoding variants associated with phenotypes^19^, or that affect cellular proliferation^20^), and studied the effects on nearby genes. Thus, estimates of properties of element-gene regulatory interactions from these datasets may not generalize to the rest of the genome.
3. *Cell type representation.* The largest CRISPRi perturbation datasets to date have been collected in a single workhorse cell line: K562 erythroleukemia cells^15,16,18–20,27,28^. This cancer cell line has been deeply profiled by the ENCODE Project, and CRISPR perturbations in this cell type have been highly valuable in understanding the relationships between different epigenomic datasets and constructing initial predictive models^16,29–32^. However, comparable datasets are still limited in other cell types, especially in non-cancer lines that may offer broader perspectives into the regulatory landscape of the human genome. Therefore, large datasets in other cell types and cell lines would be highly valuable for evaluating the ability of these predictive models to generalize across cell types and for understanding any differences between cell types.

To address these limitations, we developed and applied a new framework to survey regulatory element-gene interactions. This framework combines 3 components: (1) an improved and cost-effective targeted Perturb-seq method, called Direct-Capture Targeted Perturb-seq (DC-TAP-seq), which is built upon CRISPRi tiling^28,35,36^ , and TAP-seq^15^; (2) an approach for unbiased selection of elements and genes; and (3) a design strategy to account for statistical power and ability to detect small effects. We applied this framework to build a dataset of 9,666 randomly sampled element-gene pairs in two cell lines: K562 and human induced pluripotent stem cells (WTC11). This dataset, by virtue of its unbiased selection of elements and genes, and higher statistical power, provides an expanded view of the effect sizes, chromatin states, and gene targets of distal regulatory elements. These data validate the accuracy of recent models designed to identify enhancer-gene interactions, and also reveal gaps that have implications for building the next generation of predictive models. Overall, this work provides a foundation for expanding large-scale CRISPR studies to better understand gene regulation and to develop more accurate and generalizable predictive models.

## Results

### Direct-Capture Targeted Perturb-seq for highly powered CRISPRi screens

We developed a method for perturbing candidate regulatory elements in their endogenous locations in the genome and sensitively measuring effects on nearby genes, called Direct-Capture Targeted Perturb-seq (DC-TAP-seq) (**Fig. 1a**). This method builds on CRISPRi tiling^28,35,36^ , and TAP-seq^15^, which involves reading out the effects on gene expression using single-cell RNA-seq with a targeted PCR to enrich for a panel of genes of interest. In DC-TAP-seq, we deliver a library of gRNAs targeting a set of elements of interest; activate CRISPRi for 48 hrs to perturb expression; and use 10x Genomics 3’ single-cell RNA-seq to read out the gRNA(s) expressed in each cell as well as the expression of a selected panel of genes.

**Figure 1.**
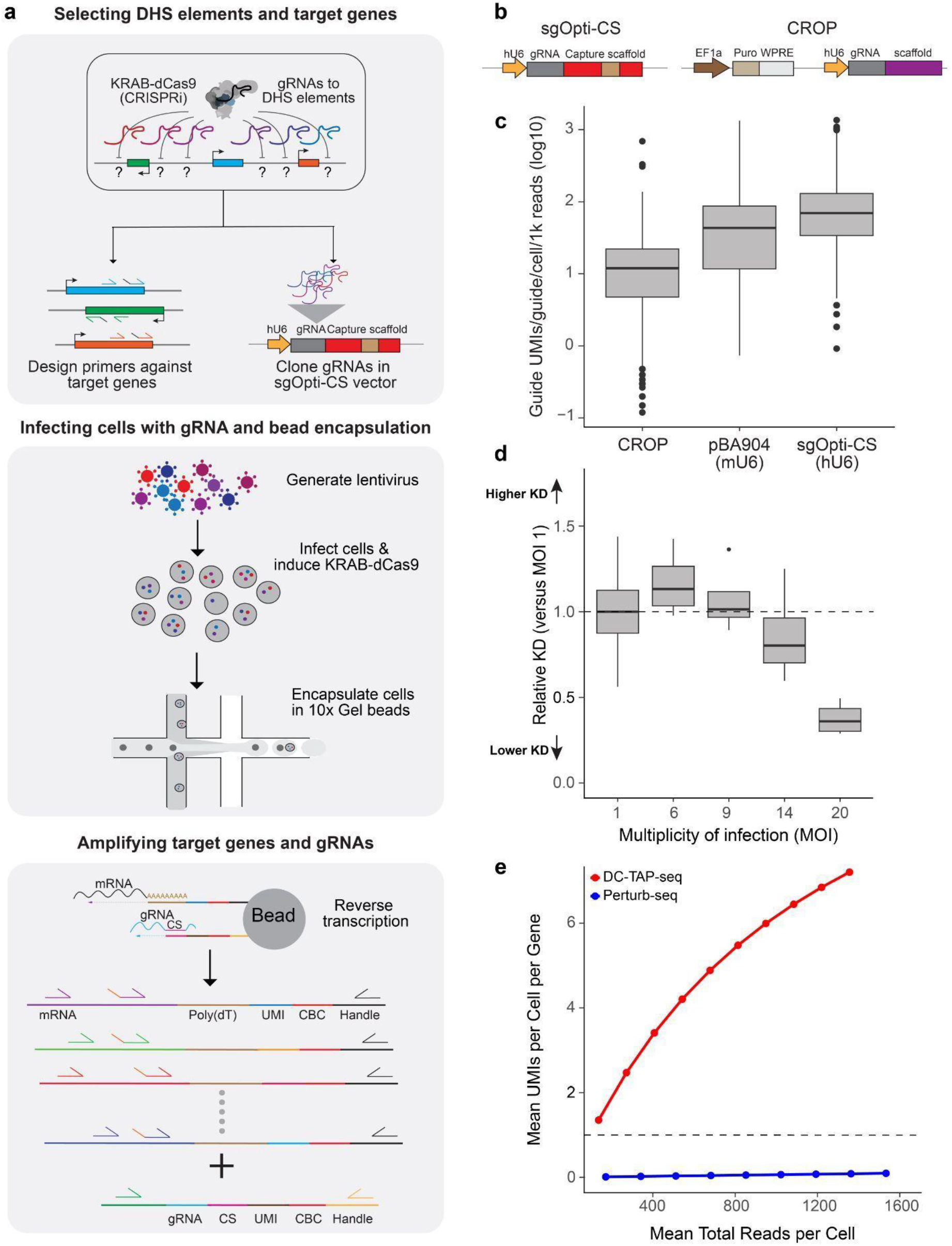
Overview of direct-capture targeted perturb-seq. **a.** DC-TAP-seq method to link regulatory elements to target genes. gRNAs are designed against DHS (DNase hypersensitive sites) elements in a particular locus and candidate target genes near those elements are selected for primer design. The pool of gRNAs is cloned in the sgOpti-CS vector and packaged in lentivirus for infecting cells expressing KRAB-dCas9, which are then processed with the 10x Genomics 3’ scRNA-seq workflow. Following bead capture and reverse transcription, mRNAs and gRNAs are amplified via nested multiplex PCRs and prepared for next-generation sequencing. **b.** Vector design for sgOpti-CS and CROP-seq showing the presence of the capture scaffold that enables direct capture of hU6 derived gRNAs from the sgOpti-CS vector. **c.** Number of guide UMIs captured per guide per 1,000 reads per cell for CROP-seq, mU6 direct-capture vector (pBA904) and hU6 direct-capture vector (sgOpti-CS). See Methods, ‘Data visualization’ section for definition of box plot elements. **d.** Relative efficiency of knockdown (KD) across different multiplicities of infection (MOI) in K562 cells, averaged across 4 gRNAs targeting GATA1 promoters and enhancers and normalized to effects at MOI = 1. **e.** UMI capture efficiency for DC-TAP-seq and Perturb-seq with the 10x Genomics 3’ scRNA-seq workflow for 93 genes in K562 across read depths. The y-axis is the average of the mean UMIs per cell per gene for the final 93 genes detected in the DC-TAP-seq experiment (out of the 97 amplified genes). The dashed horizontal line represents 1 UMI per cell per gene, an approximate threshold for obtaining high statistical power in designing DC-TAP-seq experiments.

DC-TAP-seq improves on the original TAP-seq^15^ method by increasing gRNA detection efficiency and scalability. First, we modified the multiplexed PCR strategy to simultaneously amplify selected mRNAs of interest (as in TAP-seq) as well as a short Pol-III-derived gRNA transcript through a direct capture strategy^34^ (**Fig. 1a**). We cloned an improved direct-capture gRNA expression vector with an hU6 promoter (**Fig. 1b**), which increased the number of gRNA UMIs detected by 7.3-fold versus the previous TAP-seq strategy using a CROP-seq vector (**Fig. 1c**) and by 1.6-fold versus the original direct-capture vector with an mU6 promoter (**Fig. 1c**). Second, to increase the cost-effectiveness of CRISPRi screens, we deliver gRNAs at a high multiplicity of infection (MOI), as in prior enhancer screens^14^, and analyze data with SCEPTRE^37^, testing multiple element perturbations per cell. Through dose-titration of lentivirus, we found that we could deliver MOI of up to approximately 10 without compromising the knockdown efficiency compared to that of cells infected at MOI1 (i.e., 1 guide plasmid per cell) using DC-TAP-seq (**Fig. 1d, Extended Data Fig. 1a,c,d, Supplementary Table 1**). To confirm that we maintain high mRNA UMI capture, we conducted a pilot DC-TAP-seq experiment at low MOI in K562 targeting 97 genes (**Supplementary Table 2**) and measured the mRNA UMI capture efficiency as a function of total reads per cell. Our improved DC-TAP-seq protocol indeed captured 90-fold more UMIs than regular 10x scRNA-seq (Perturb-seq) at a fixed sequencing depth (∼1400 reads) (**Fig. 1e**). As expected, our protocol achieves an average of 1 UMI per cell per gene at substantially lower read depth than Perturb-seq, enabling ≥80% power to detect 25% effect sizes with only 250-500 cells per target (**Fig. 1e, Extended Data Fig. 1b**). Altogether, DC-TAP-seq improves gRNA detection, reduces the number of cells, and preserves knockdown efficiency, while retaining the advantages in sensitivity, power, and cost of the original TAP-seq compared to untargeted Perturb-seq approaches.

### Testing thousands of random element-gene connections in K562 and hiPSC cells

We applied DC-TAP-seq to experimentally map thousands of candidate element-gene regulatory interactions to understand the global architecture of distal gene regulation. We designed these DC-TAP-seq experiments to measure a total of ∼10,000 random distal element-gene (DE-G) regulatory interactions across two cell types, K562 and WTC11 hiPSCs, with ≥80% statistical power to detect 25% effects on gene expression (5,059 DE-G interactions in K562, 4,607 in WTC11) (**Supplementary Table 3**). We first selected 25 loci in K562 and 24 loci in WTC11. As introduced above, prior studies employed a biased selection strategy for both target genes and elements (*e.g.*, high H3K27ac, genes encoding transcription factors, etc.)^14^. Here, we aimed to create a dataset that would better generalize to the entire genome. As such, our only selection criteria for the 49 loci targeted across K562 and WTC11 were that they contain at least one gene (the “index gene”) expressed at a sufficient level for statistical power (≥50 transcripts per million (TPM) in the tested cell type). We then selected ∼40 DHS peaks at random within a 0.5-4.5 Mb window around the index gene (**Fig. 2a**). We then designed our panel to measure effects on all genes in the locus that were expressed above 30 TPM. We did not target regions without DHS signal, because targeting regions without DHS or ATAC signal does not typically impact gene expression outside of the gene body^28,38^. Different loci were selected in each cell type, so as not to bias the selection of elements or genes to those that are shared across cell types. We included targeted PCR primers to read out effects on total 303 and 203 genes in these loci in K562 and WTC11 respectively (**Supplementary Table 4**). In total, the CRISPRi libraries included 14,968 and 16,201 gRNAs total, including ∼15 gRNAs per element, 800 non-targeting and safe-targeting negative control gRNAs, and 1,178 and 175 gRNAs targeting previously studied positive control enhancers and promoters for K562 and WTC11, respectively.

**Figure 2.**
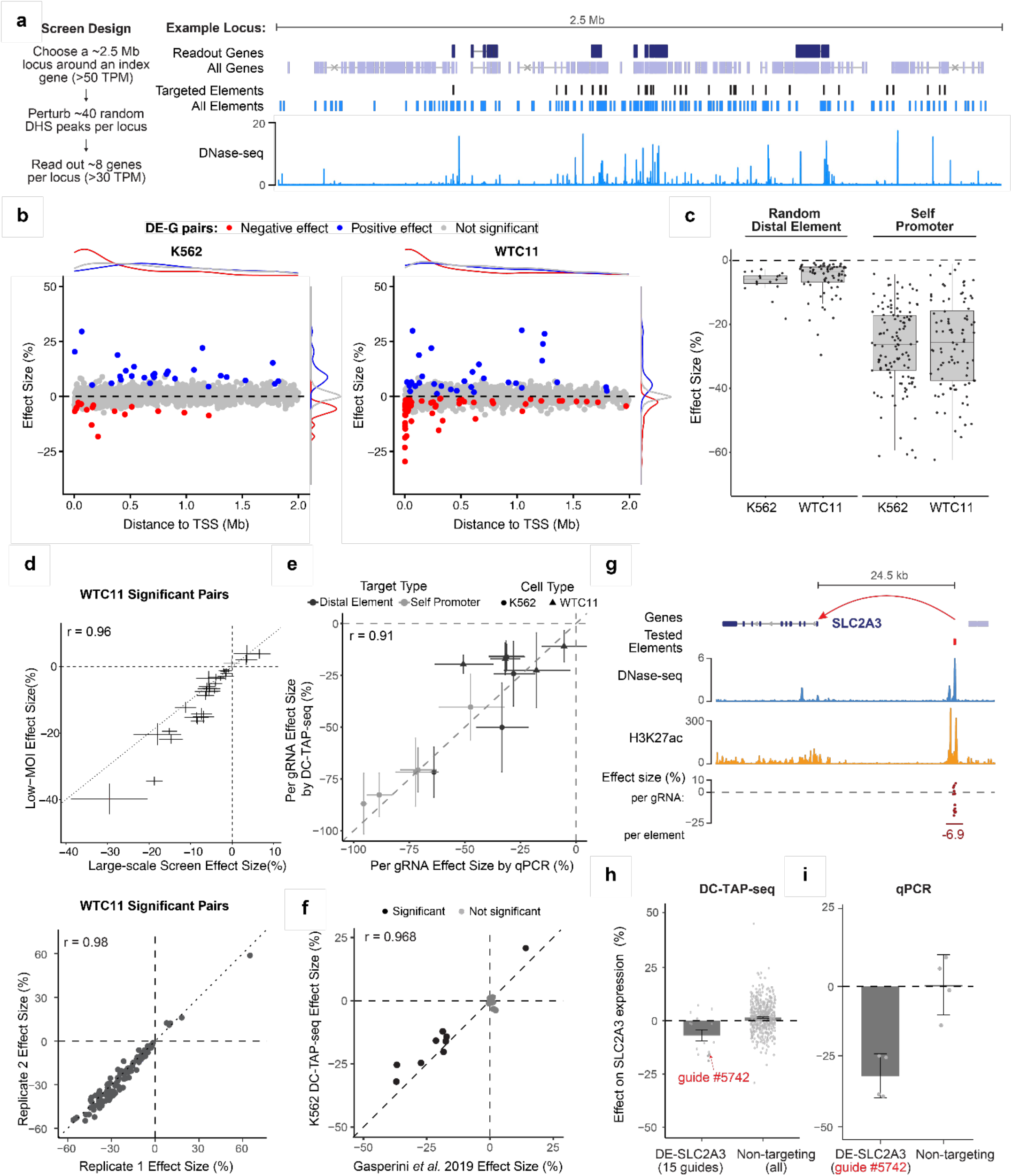
DC-TAP-seq enables highly sensitive mapping of thousands of randomly selected distal regulatory elements. **a.** Left: Design procedure for random element DC-TAP-seq experiments. Right: Example locus showing random distal elements (DNase-seq peaks, dark blue) and genes (dark blue) that were read out with DC-TAP-seq. We selected 24 or 25 loci per cell type, and targeted ∼40 random elements per locus with ∼15 gRNAs each. We measured effects on the expression of ∼8 nearby genes per locus (range: 1–34). **b.** CRISPRi effect sizes (% change in gene expression) as a function of distance between each perturbed DE and TSS of the readout gene. Each dot represents a single DE-G pair tested in K562 (left) or WTC11 (right). Red, blue, and gray lines represent density along each axis. One outlier (K562, effect size 72%) is not shown in the plot. **c.** Box plot showing the distribution of negative effect sizes for all significantly regulated randomly selected DE-G pairs in K562 and WTC11 (left), compared to effects of targeting gene promoters on the expression of that same gene (“Self Promoter”, right). **d.** Reproducibility of effect sizes across experiments in WTC11: (top) biological replicates - large scale screen and low MOI validation screen, and (bottom) technical replicates - random grouping of 10x lanes for significant element-gene pairs. **e.** Comparison of effect sizes between DC-TAP-seq and qPCR measurements for selected elements (Distal Element) and TSS-targeting (Self Promoter) guides from K562 and WTC11. Effect size is represented as percent change in gene expression relative to control guides, with –100% indicating maximal knockdown. Error bars: 95% confidence intervals (c.i.) (n=2-4 biological replicates) **f.** Comparison of effect sizes for shared element-gene pairs between DC-TAP-seq and previously published Perturb-seq data from Gasperini et al^14^. Dark gray: Significant in both studies. Light gray: Not significant in both studies. **g.** Example locus for a significant DE-G pair identified in WTC11. Tracks show a link between SLC2A3 and a distal element (chr12: 7960676-7960977; high H3K27ac and 24.5 kb away), DNase-seq signal, H3K27ac ChIP-seq signal (read pileup coverage), and the effect sizes of gRNAs on *SLC2A3* expression. **h.** Comparison of effect sizes on *SLC2A3* expression for gRNAs targeting the element DE-SLC2A3 shown in panel (g) (chr12:7960676-7960977), and all non-targeting guides. Guide #5742 was validated by qPCR in panel (i). Error bars: 95% c.i. (SCEPTRE). **i.** Comparison of effect sizes on *SLC2A3* gene expression for guide #5742 and a non-targeting guide measured by qPCR. Error bars represent 95% c.i. (n=4 biological replicates).

We obtained highly powered and reproducible DC-TAP-seq measurements of DE-G regulatory interactions. DC-TAP-seq experiments in K562 and WTC11 yielded 95,997 and 180,116 cells at an MOI of 7.3 and 5.1, respectively (**Supplementary Table 5, Extended Data Fig. 2a,b**). This depth provided an average coverage of 47 and 56 cells per gRNA (**Extended Data Fig. 2b**), and ≥80% power to detect 25% effects for most tested random DE-G pairs (84% and 85%, **Supplementary Table 3**). A significant fraction of random DE-G pairs had sufficient power to detect 15% effects (36% and 65%, **Supplementary Table 3**, **Supplementary Table 6**). We analyzed these data with an updated differential expression analysis pipeline based on SCEPTRE (see **Methods)**, which produced well calibrated *p*-value distributions (**Extended Data Fig. 2c**) and yielded highly reproducible estimates of element-gene effect sizes across technical replicates (Pearson’s R = 0.95 for K562 and 0.98 for WTC11, **Fig. 2d, Extended Data Fig. 1e**). Most positive control element-gene pairs showed significant effects (promoters: 32 of 39 in K562, and 4 of 7 in WTC11; distal elements: 20 of 34 in K562, and 0 of 2 in WTC11). (**Supplementary Table 6**). Of the 10 promoter and 16 distal element positive controls that did not show significant knockdown, the majority did not have sufficient power for detection in this experiment. 2 positive control promoters (both K562) and 6 distal elements (5 in K562, and 1 in WTC11) did have good power for 15% effects, but did not show a significant effect (**Supplementary Table 6**). Taken together, DC-TAP-seq achieved the expected results for both promoter and distal element positive control gRNAs in the majority of cases across both cell lines when DE-G pairs were well powered. Effect sizes for significant DE-G pairs from K562 DC-TAP-seq were also highly correlated with those from a previous K562 Perturb-seq experiment^14^ for 9 regulated shared DE-G pairs (Pearson’s R = 0.97, **Fig. 2f**). Together, this demonstrates the high technical quality of these large scale DC-TAP-seq experiments.

Of the 9,666 DE-G pairs tested, and after filtering for promoters, positive controls, and elements in gene bodies (see **Methods**), we identified 18 (K562) and 66 (WTC11) (84 total) DE-G pairs that led to significant decreases in gene expression (“negative effects”) and 28 (K562) and 33 (WTC11) (61 total) that led to significant increases (“positive effects”) (FDR = 20%). We found 1,796 (K562) and 2,915 (WTC11) additional random DE-G pairs that were tested with good power for 15% effects but were not significant (**Supplementary Table 6**). Significant DE–G pairs in both cell models were enriched for being located within 100 kb of the target gene TSS (**Fig. 2b**): of 145 significant pairs, 55 were located within 100 kb, compared to 986 of 9,666 tested pairs overall (fold enrichment = 3.7; one-sided hypergeometric test, *P* = 3.8 × 10^−19^). This result is consistent with previous perturbation studies^16^. Among the 84 significant DE–G pairs with negative effects, 45 (53.6%) were located within 100 kb of the target-gene TSS, and 70 (83.3%) had effect sizes below 10% (**Supplementary Table 3**). The median negative effect size of these DE-G pairs was 4.3%, compared to 26% for targeting promoters on the same genes (**Fig. 2c**). The majority of regulatory elements with a significant effect in either direction connect to a single gene, with only 2.2% of elements in K562 and 9.1% in WTC11 connected to more than one gene (**Extended Data Fig. 3a**). Conversely, the majority of genes are connected to a single element (**Extended Data Fig. 3b**). Considering that we did not measure the effect of every element or every gene in each locus, these connectivity patterns are broadly consistent with prior CRISPR screens.

To validate a set of significant DE-G pairs, we repeated the DC-TAP-seq experiment for a subset of 23 significant and 5 not significant DE-G pairs (28 DE-G pairs total) in WTC11 at a range of MOIs: low MOI (1.1, 1.5) and high MOI (12.6, 14.4), compared to the medium MOI (5.1) of our initial experiment (**Extended Data Fig. 4, Extended Data Fig. 5**). 23 of the 23 significant DE-G pairs from the initial experiment were also significant in these validation experiments, with very high correlation in effect sizes (Pearson’s R = 0.96 and 0.91 for low and high MOI, respectively) (**Fig. 2d, Extended Data Fig. 1d**). In addition, both validation screens were better powered than the initial screen for the 28 DE-G pairs tested. Four of the five pairs that were not significant in the original screen had small effect sizes that the original screen was underpowered to detect, and each validation screen detected four of them (**Extended Data Fig. 6**). At high MOI (>10), the observed effect sizes were highly correlated with but notably weaker in magnitude than in the medium- or low-MOI experiments (**Fig. 1d, Extended Data Fig. 1a,c and d**), possibly due to competition for limited Cas9 from multiple gRNAs.

To further validate the effect sizes of a subset of elements, we selected one of the top-performing guides for each of 9 distal elements and 5 self promoters (i.e., guides targeting the TSS of the same gene). We infected these guides in an arrayed fashion and found that all 14 guides showed a qPCR effect size in the same direction as the effect size measured by DC-TAP-seq, and of these, 13 showed significant decreases in gene expression measured by qPCR (**Fig. 2e)**. The qPCR effect sizes on the target genes correlated with that measured by DC-TAP-seq (Pearson’s R = 0.91, **Fig. 2e**). As a visual example of the data, perturbation of a H3K27ac-marked element located 24.5 kb upstream of *SLC2A3* led to a 6.9% decrease in gene expression by element-level analysis (across all guides targeting the element) in the large scale screen (**Fig. 2g,h**). We selected one of these guides (guide #5742) for orthogonal validation by qPCR, and confirmed reduced *SLC2A3* expression relative to a non-targeting guide (**Fig. 2i)**.

Together, this experiment provides the first high-sensitivity CRISPRi dataset utilizing a randomly sampled design to explore global properties of accessible elements and their effects on distal gene regulation. We next further examined the 9,666 randomly selected DE-G pairs, and compared their properties to DE-G pairs discovered in previous studies.

### Most regulatory elements have small effects on gene expression

The majority of the significant DE-G regulatory interactions with a negative effect had very small effect sizes (0.874% - 29.6%, median: 4.3%). Out of 84 significant DE-G pairs, 70 have an effect size <10% **(Fig 2b,c)**. The magnitude of effect sizes was therefore much smaller, on average, than the effect size distributions reported in previous studies, in part due to differences in statistical power.

To compare effect sizes across studies, we uniformly reprocessed 4 previous enhancer Perturb-seq experiments^19,20,27^ and developed a simulation-based framework for estimating statistical power. In this framework, we (i) for each gene, estimate mean and variance from the real Perturb-seq dataset; (ii) for a given element-gene pair, randomly sample cells from the experiment and replace their expression with simulated values representing a specific effect size; (iii) test the simulated cells for a significant effect on expression, using the same testing procedure, covariates, and significance threshold as for the real data; and (iv) repeat this procedure to empirically estimate power (see **Methods**).

Indeed, the effect sizes of significant DE-G pairs with negative effects in this DC-TAP-seq dataset were smaller than in these previous studies (median: 4.3% versus 23.1%, **Fig. 3a**). This difference was not due to experimental differences in perturbation efficiency, because shared elements across studies had similar effect sizes (**Fig. 2f**). Rather, this DC-TAP-seq dataset was much better powered to identify small effect sizes (**Fig. 3b**). For example, at an effect size of 15%, our DC-TAP-seq experiments had good power for nearly half of the tested pairs (4,817 out of 9,666 pairs with ≥80% power). In comparison, previous experiments achieved such power for only 0-11% of tested pairs (**Fig. 3b**). This large improvement is achieved by the high efficiency of capturing mRNA and gRNA UMIs with DC-TAP-seq and by profiling a large number of single cells per element (**Fig. 1e, Supplementary Table 7**). We note that previous studies also identified DE-G pairs with larger effect sizes (*e.g.*, >30%) that were not represented in our dataset, which appears to be due to selection biases in the elements tested (see below). Some elements with such large effect sizes were included as positive controls in our DC-TAP-seq experiment, and had effect sizes that matched previous experiments (**Fig. 2f**).

**Figure 3.**
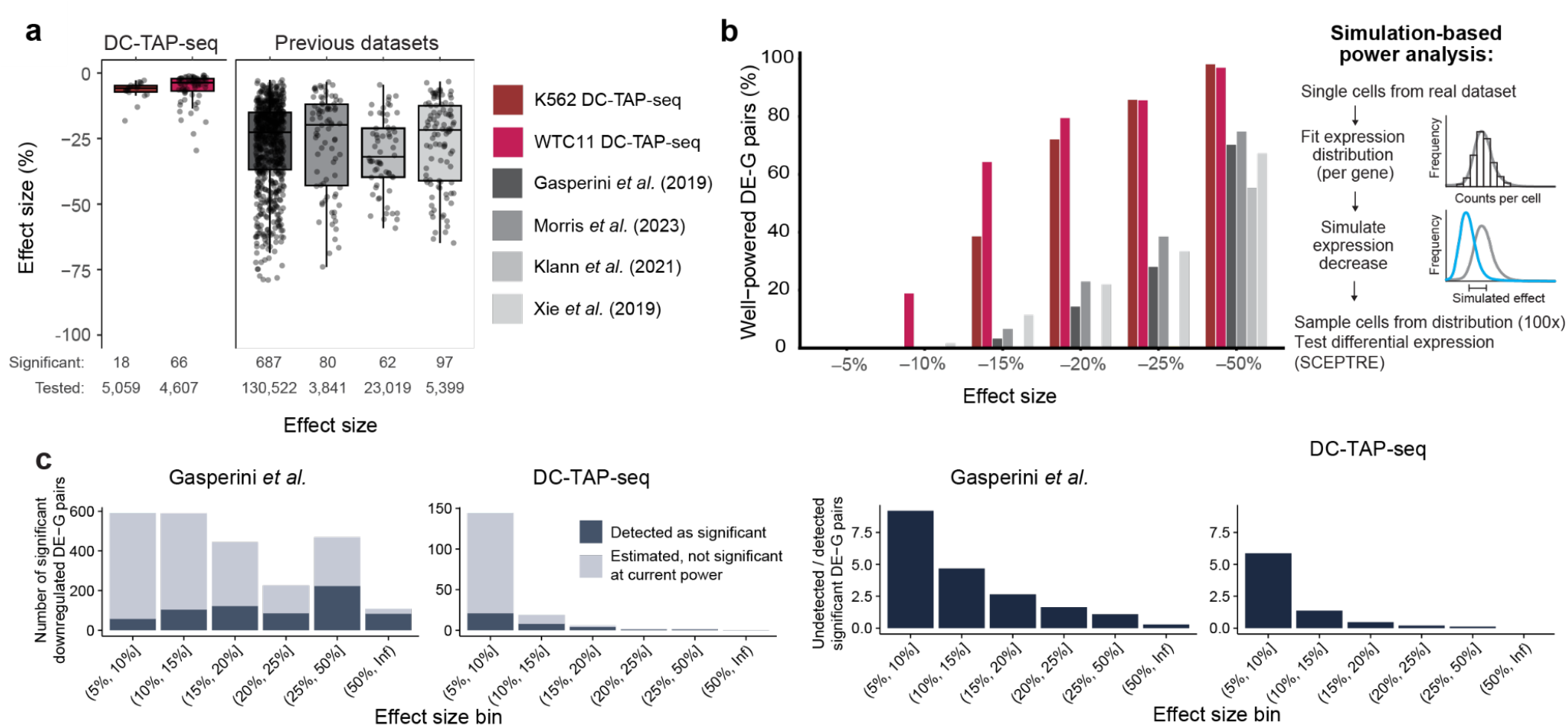
Most randomly selected regulatory elements exert only weak repression of gene expression. **a.** Boxplot of effect sizes for significant down-regulated random DE-G pairs in (left) K562 and WTC11 DC-TAP-seq and (right) four previous Perturb-seq datasets. Box plots: median ± IQR. Points: DE-G pairs. Bottom: Numbers of tested and significant elements. **b.** Left: Proportion of well-powered promoter and DE-G pairs within 1Mb at various simulated effect sizes across the DC-TAP-seq (red) and other (gray) experiments. Right: Overview of simulation-based framework to estimate statistical power to detect certain effect sizes in a given Perturb-seq experiment. Starting with the single-cell count matrix and guide assignments, we fit a negative-binomial model to single-cell counts for each gene; impose a particular decrease in mean expression; adjust the expression of the target gene in cells assigned to that guide; and re-run differential expression testing with SCEPTRE, using all of the same covariates and p-value thresholds as in the real analysis. We repeat this sampling and differential testing 100 times and calculate power as the fraction of simulated tests that are significant. **c.** Left: Number of significant DE-G pairs with negative effect sizes that were detected as significant (dark grey) or undetected as significant (light grey) for different effect size bins for Gasperini *et al.* dataset^14^ (left) and the random DC-TAP-seq dataset (right). The total number of significant DE-G pairs for an effect size bin was computed as: (# detected significant positive DE-G pairs in bin) / (sum of power to detect maximum effect size in bin across all tested pairs) x (total number of tested pairs) (see **Methods**). Right: Ratio of estimated undetected to detected significant DE-G pairs with a negative effect size for each effect size bin.

Given these observations about statistical power, there are likely to be a large number of regulatory elements with smaller effect sizes on gene expression (*e.g.*, 5–10%) that were tested but not detected as significant in this work and previous Perturb-seq studies. When we re-examined the largest of these datasets (Gasperini *et al*)^14^, we found that there are likely to be 9-fold more DE-G pairs with 5–10% effect sizes than the study was powered to detect individually (**Fig. 3c**). Even with the increases in power gained by this experimental design there are likely to be 5-fold more DE-G pairs with effect size 5-10% than our study was powered to detect **(Fig. 3c)**.

Thus, in a random survey of DE-G pairs across the genome with negative effects on gene expression, a majority of DE-G regulatory interactions have small effect sizes on gene expression (<10%), which previous studies have not been powered to detect. These data also highlight the importance of future study designs that are powered to detect small effect size regulatory interactions.

### Distinguishing direct *cis*-regulatory interactions from indirect effects

Before further investigating the properties of DE-G pairs, we considered that this CRISPR experiment, like other Perturb-seq studies, has the potential to detect not only direct, *cis*-regulatory interactions (*e.g.*, an enhancer regulates a nearby target promoter via 3D contact)^28^ but also indirect effects on nearby genes, in which an element directly regulates nearby gene A which produces a protein that happens to regulate nearby gene B^16^ (**Fig. 4a**).

**Figure 4.**
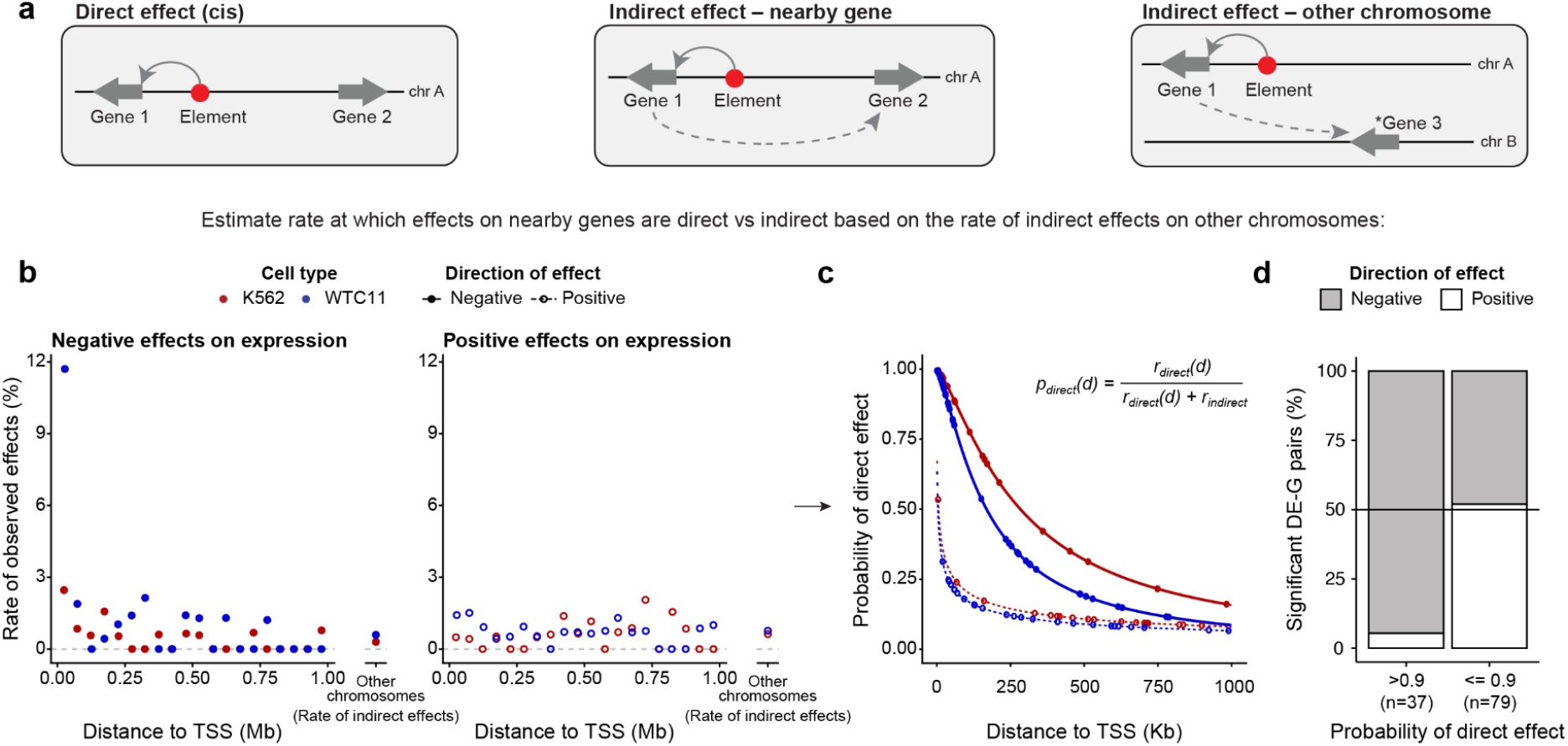
Distinguishing *cis*-regulatory interactions from indirect effects. **a.** Direct effects (left) represent elements that act in *cis* on nearby genes on the same chromosome, *e.g.* such as classical enhancers that physically interact with their target promoter. Indirect effects (center, right) represent cases where elements have a direct effect on one gene (Gene 1) which then in turn affects the expression of another gene either nearby on the same chromosome (Gene 2, center) or on another chromosome (Gene 3, right). Thus, effects of CRISPR perturbations to a given gene may result from either direct or indirect effects. We use the rate of effects on genes on other chromosomes as an estimate of the frequency of such indirect effects. **b.** The rate of significant DE-G pairs (# significant / # tested) as a function of genomic distance from the distal element to the gene TSS (for pairs within 1 Mb each other on the same chromosome) and for DE-G pairs in which we pair a DE with genes on other chromosomes. Each dot represents one distance bin. The rate of significant DE-G pairs is higher for negative effect sizes at short genomic distances, but otherwise is on the same order of magnitude as the rate for pairs on other chromosomes **c.** Probability that a significant DE-G pair represents a direct effect, as a function of genomic distance (*x*-axis; up to 1Mb), direction of the effect (filled: negative effects; open: positive effects), and dataset (red: K562; blue: WTC11). Line: fitted estimate. Dots: significant DE-G pairs. **d.** Among the significant DE-G pairs that are most likely to be direct effects (>0.9 probability), most have negative effects on gene expression. Among the pairs with <0.9 probability of being a direct effect, there is a similar number of positive and negative effects. Only significant DE-G pairs within 1 Mb of the gene TSS are shown.

We designed a procedure to estimate the probability that a given DE-G pair in a CRISPR experiment represents a direct, *cis* effect, based on the assumption that the rate of direct effects will depend on genomic distance whereas indirect effects will not. We first computed the rate of effects of DEs on genes on other chromosomes (∼0.5% of tested DE-G pairs), which we assume to be indirect effects or false discoveries (**Fig. 4b**). We confirmed that the rate of such indirect effects observed for the elements and genes tested in our DC-TAP-seq experiment was similar to those of larger sets of elements and genes tested in a separate Perturb-seq experiment^14^ (**Extended Data Fig. 16b,c**). Next, we calculated the rate of effects of DEs on nearby genes on the same chromosome, which are a mix of direct effects, indirect effects, and false discoveries (**Fig. 4b**). We then calculated the difference between these two rates to derive a probability that each significant DE-G pair represents a direct *cis*-regulatory interaction, as a function of genomic distance, direction of effect, and dataset (**Fig. 4c**, **Methods**).

Overall, we estimate that 59% and 36% of the DE-G pairs with significant negative effects in K562 and WTC11, respectively, represent indirect effects, rather than *cis*-regulatory interactions. For negative effect sizes, DE-G pairs located <54 kb (K562) or <32 kb (WTC11) apart are >90% likely to represent direct *cis* effects, whereas DE-G pairs located >648 kb (K562) or >386 kb apart are <25% likely (**Fig. 4c**). For positive effect sizes, most DE-G pairs at any distance are more likely to represent indirect effects (**Fig. 4c**). As expected, the DE-G pairs likely to represent direct *cis* effects had predominantly negative effect sizes (83% in K562 and 97% in WTC11), whereas DE-G pairs likely to represent indirect effects did not (41% in K562 and 52% in WTC11) (**Fig. 4d**, **Extended Data Fig. 7**).

Thus, in a random survey of distal elements and genes, indirect effects appear to be quite frequent, likely capturing effects of distal elements on dosage-sensitive genes that have global regulatory effects. Accordingly, in subsequent analysis to assess the properties of *cis*-regulatory interactions, we considered the probability at which each DE-G pair is a direct versus indirect effect.

### An expanded repertoire of distal regulatory elements

Our screen design and our high power to detect even low effect sizes provides an improved view of the landscape of element-gene regulatory interactions, contrasting with previous large-scale CRISPR studies. We therefore next utilized the DC-TAP-seq data to interrogate the chromatin states and category of target gene of the significant DE-G pairs.

To characterize the chromatin states of each tested distal element, we used ChIP-seq data for H3K27ac (a mark of classical enhancers), H3K27me3 (a mark of Polycomb-bound elements), and CTCF (a transcription factor that shapes 3D genome architecture). We defined five categories of elements: (1) High H3K27ac elements, (2) H3K27ac elements, (3) CTCF elements (elements with CTCF signal and no H3K27ac peak), (4) H3K27me3 elements (no CTCF or H3K27ac), and (5) no H3K27ac (no CTCF and no H3K27me3) (**Fig. 5a**). We then compared the proportion of elements in each category between the sets of DE-G pairs that were tested, well-powered, or significant.

**Figure 5.**
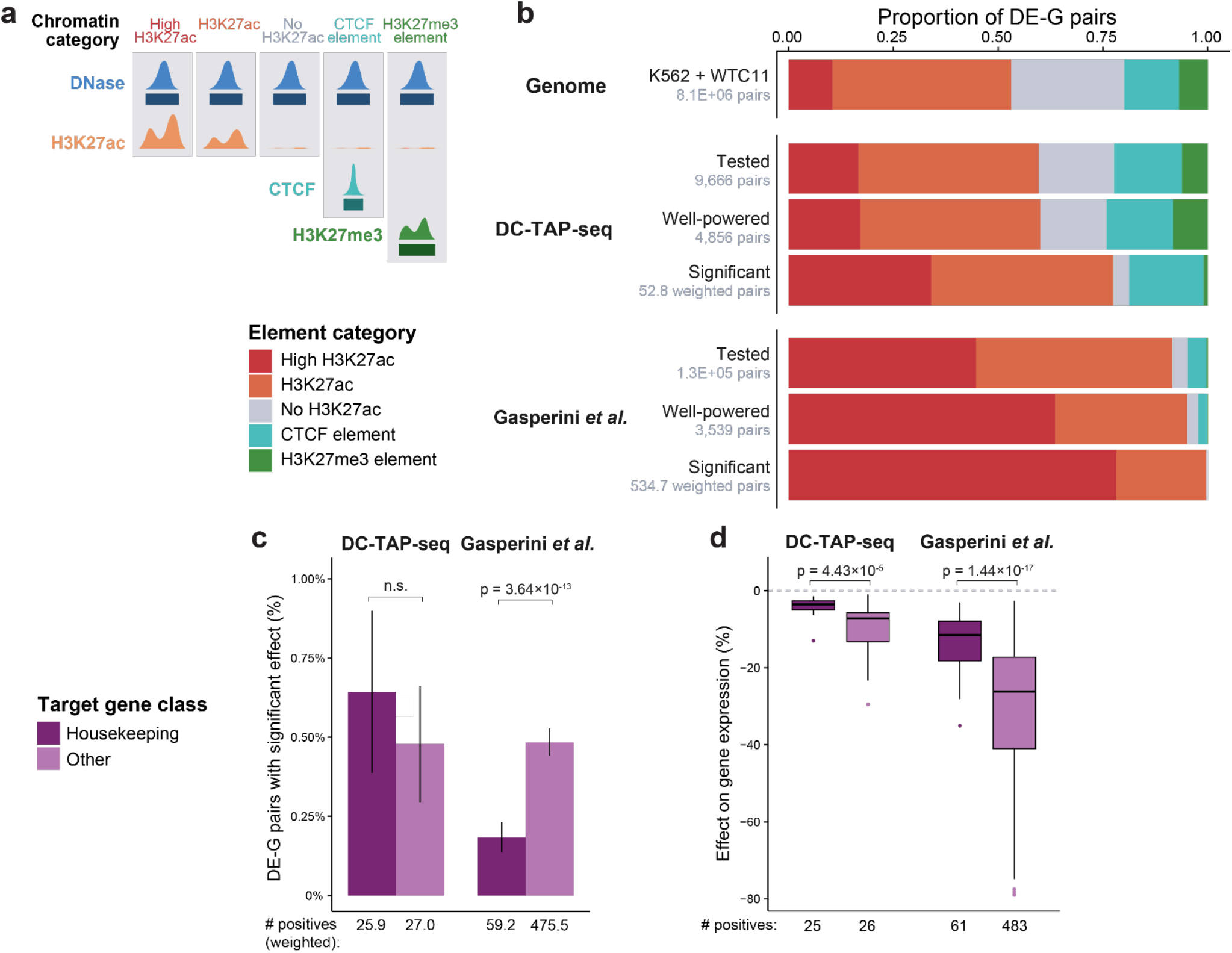
An expanded view of distal gene regulation. **a.** Schematic showing theoretical ChIP-seq signals and peaks for the five chromatin categories (see Methods for precise definitions). **b.** Proportion of DE-G pairs for which the element is classified into different chromatin categories for all genome-wide DE-G pairs, tested DE-G pairs, well-powered DE-G pairs (tested with power to detect a 15% effect size in ≥80% of pairs, plus significant pairs with lower power), and DE-G pairs with negative effect sizes found as significant (weighted by probability of direct effect) for DC-TAP-seq and Gasperini *et al.*^14^ **c.** Fraction of tested DE-G pairs that were found to have a significant negative effect (weighted by probability of direct effect), stratified by dataset and target gene class. Error bars indicate 95% confidence intervals around the observed proportion, with the standard error estimated from the Agresti– Coull adjusted proportion and lower bounds are truncated at 0. Numbers below indicate the number of significant pairs with the target gene in the indicated class, weighted by probability of direct effect. P-values report significance from two-sample chi-squared tests with Yates’ continuity correction, adjusted with Benjamini-Hochberg correction. **d.** Effect sizes for significant DE-G pairs with negative effect sizes (probability of direct effect > 50%) stratified by target gene class, for DC-TAP-seq and Gasperini *et al.*^14^ P-values report significance from two-sided Wilcoxon rank-sum tests comparing housekeeping and other target genes within each dataset.

Due to the intentional focus of our study on random DHS peaks across each locus, the proportion of distal elements in these categories differed markedly between DC-TAP-seq and other datasets (**Fig. 5b**). DC-TAP-seq tested 59.6% DE-G pairs with elements marked by H3K27ac, while Gasperini *et al*^14^. tested 91.9%. This difference is also present in the significant DE-G pairs: adjusting for estimated indirect effects, 77.5% of the regulatory DE-G pairs with a negative effect size we identified corresponded to classical enhancers, compared to 99.6% of regulatory DE-G pairs from Gasperini *et al.*^14^ (**Fig. 5b**). Across a separate collection of previous CRISPR datasets^29^, 76.0% of tested DE-G pairs and 96.7% of regulatory pairs had H3K27ac-marked elements (see **Methods**).

Among the DE-G pairs not marked by H3K27ac in DC-TAP-seq (26% of both positive and negative effect significant pairs, 38 of 145) 58% corresponded to elements bound by CTCF (22 of 38) (**Fig. 5b, Supplementary Table 3**). As CTCF is a well characterized loop-anchor protein, such elements may regulate gene expression by altering 3D contacts between nearby enhancers and promoters^39,40^. Their elevated frequency in DC-TAP-seq (15% of significant pairs) versus previous experiments (<1% of significant pairs)^29^ suggests that regulatory effects of CTCF-bound sites may be more prevalent than previously estimated. Notably, while K562 and WTC11 differed in their frequencies of certain chromatin states genome-wide (**Extended Data Fig. 9a**), the element-gene pairs with significant regulatory effects were similar with respect to their chromatin states, and CTCF-bound sites were discovered in both cell types (**Extended Data Fig. 9a**). Based on observations from previous studies^41,42^, we expected that the observed negative effects on target gene expression are due to disruption of CTCF binding at the targeted elements. To test this, we conducted CTCF ChIP-seq to evaluate the impact of CRISPRi perturbation on CTCF binding at two of the significant CTCF-bound elements (**Extended Data Fig. 10**). At one of these elements (*FAM111A* locus), we did indeed observe a decrease in CTCF ChIP-seq signal upon CRISPRi perturbation (*P* = 0.047 and P = 0.023, Wilcoxon two-sided test for the two guides tested), while at the other (*CDC37*) we did not. We cannot therefore conclude a shared mechanism across the significant CTCF-bound elements. Indeed, these data suggest that the effect on gene expression of targeting elements bound by CTCF can be mediated by both CTCF-dependent and CTCF-independent mechanisms.

We analyzed a larger compendium of ENCODE ChIP-seq data for each cell type, and found that elements with significant negative effects were broadly enriched for transcription factor occupancy and active histone modifications, corroborating their assignment as regulatory elements (**Extended Data Fig. 14)**.

We also re-examined the regulatory landscape of housekeeping genes (here, genes that are ubiquitously and uniformly expressed across cell types, see **Methods**). Previous studies have reported that the promoters of housekeeping genes are less responsive to enhancers in reporter assays^43–45^. In fact housekeeping (P2) promoters were found to have on average 10-fold fewer enhancers than non-housekeeping (P1) promoters^43^. Genome-wide CRISPR experiments have similarly found that housekeeping genes have far fewer enhancers than other genes^16^. For example, in the Gasperini *et al.*^14^ dataset, DEs paired with housekeeping genes were 2.6-fold less likely to have a significant effect than DEs paired with other genes (**Fig. 5c**). However, this conclusion is potentially affected by limitations in statistical power.

The fraction of DE-G pairs with significant negative effect sizes did not differ significantly between housekeeping genes and other genes (p = 0.33, chi-squared test with Benjamini-Hochberg correction) (**Fig. 5c**). However, the effect sizes of regulatory elements targeting housekeeping genes with a negative effect (51 total DE-G pairs) were significantly smaller (mean −4.0% vs −9.8% for other genes; p*_Wilcox_* = 4.43×10^−5^; **Fig. 5d**), in a range that previous studies were underpowered to detect (**Fig. 3b**). The same trends held when analyzing K562 and WTC11 DC-TAP-seq data separately (**Extended Data Fig. 9b,c**). For example, DC-TAP-seq identified a regulatory element ∼30 kb away from the housekeeping gene *TP53*, which showed the chromatin state of a classical enhancer including DNase-seq and H3K27ac signals (**Extended Data Fig. 11**). Thus, housekeeping genes have more distal regulatory elements than previously estimated, which appear to be similar to enhancers for non-housekeeping genes but with smaller effects on expression.

To confirm these observations through independent genetic perturbations, we examined expression quantitative trait loci (eQTLs) in lymphoblastoid cell lines, where we categorized elements and genes in a similar way. Indeed, both housekeeping and other genes showed enrichment for eQTL variants in distal elements (**Extended Data Fig. 12**), although further functional testing will be needed.

Thus, the unbiased selection strategy and high statistical power of the DC-TAP-seq screens provides an updated view of the landscape of distal regulatory elements, revealing a prevalence of CTCF-bound regulatory elements and many weak interactions with housekeeping genes. Further, our data hint at the complexities of CTCF-mediated control of gene expression. Further work will be needed to test thousands more DE-G connections in order to expand the number of DE-G pairs in each element category, particularly for CTCF-bound elements, and fully understand their role in control of gene expression. The data here can be used to guide element selection in future studies.

### Predictive models identify strong enhancers but not other types of element-gene pairs

Large CRISPR perturbation datasets have previously been used to train and benchmark predictive models of enhancer-gene regulatory interactions^29,30^. Given that the DC-TAP-seq CRISPR dataset contains element-gene pairs with distinct properties from previous experiments, we examined predictions of enhancer-gene regulatory interactions from ENCODE-rE2G^29^, a supervised model trained on a combination of previous CRISPR datasets^15,16^. Indeed, the performance of ENCODE-rE2G varied substantially between certain subsets of this DC-TAP-seq dataset:

#### Direct versus indirect effects

As expected, ENCODE-rE2G had a higher recall for DE-G pairs that were more likely to be direct versus indirect effects (46% vs 0%, **Fig. 6a**). Notably, the relationship between indirect effects and genomic distance (**Fig. 4c**) can explain in part why ENCODE-rE2G and other models appear to perform worse at long genomic distances^29^ — because the “gold-standard” CRISPR data is likely to be a mix of direct and indirect effects.

**Figure 6.**
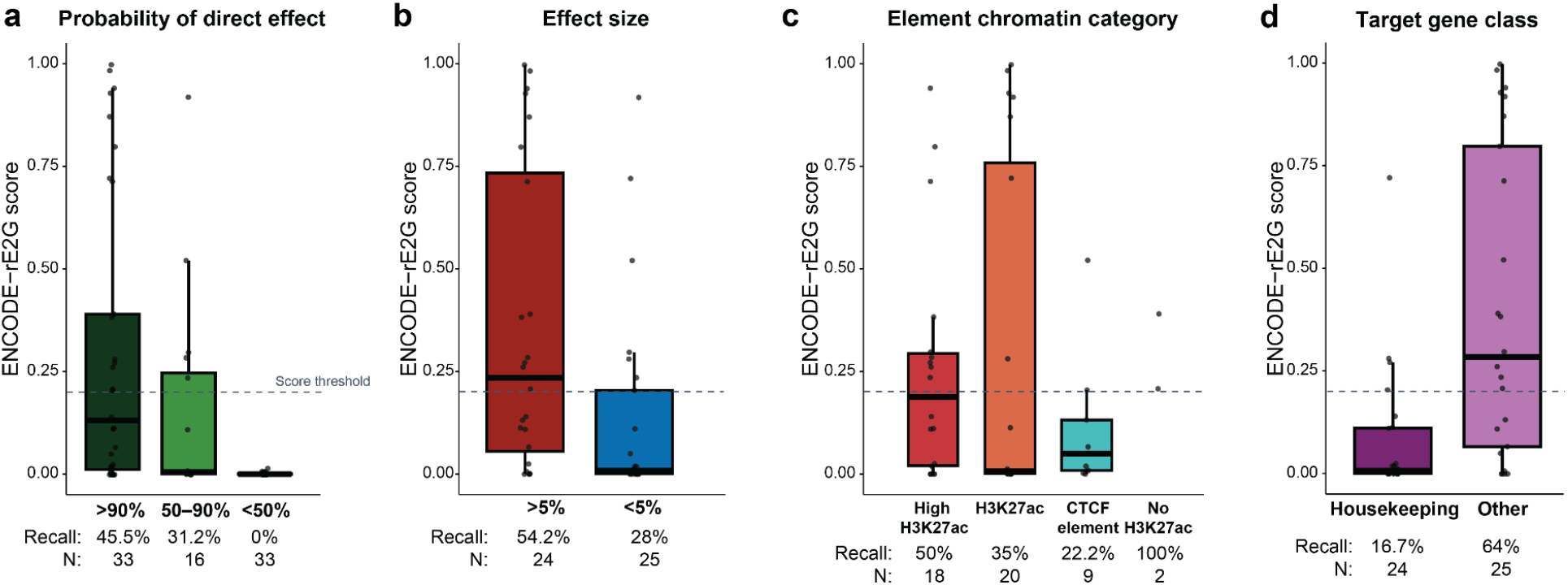
Sensitivity of predictive model for different categories of DC-TAP-seq DE-G pairs. **a.** ENCODE-rE2G scores for significant downregulated DE-G pairs from DC-TAP-seq, stratified by probability of direct effect. Recall is defined as the percent of pairs with scores above the model threshold. To annotate pairs with ENCODE-rE2G scores, elements were expanded to 500 bp and merged if overlapping. **b.** ENCODE-rE2G scores for significant downregulated DE-G pairs from DC-TAP-seq with a probability of direct effect greater than 50%, stratified by effect size. **c.** ENCODE-rE2G scores for significant downregulated DE-G pairs from DC-TAP-seq with a probability of direct effect greater than 50%, stratified by chromatin category of the element. **d.** ENCODE-rE2G scores for significant downregulated DE-G pairs from DC-TAP-seq with a probability of direct effect greater than 50%, stratified by class of the target gene.

#### Effect sizes

ENCODE-rE2G achieved higher recall for DE-G pairs with >5% effect sizes than those with <5% effect sizes (54% vs 28%, **Fig. 6b**). This may be in part due to the fact that the CRISPR dataset previously used to train the ENCODE-rE2G model^29^ has relatively few examples of DE-G pairs with <5% effect sizes and such effects would presumably be more difficult to distinguish.

#### Chromatin states

DE-G pairs corresponding to classical enhancers (marked by H3K27ac) were predicted with higher recall by ENCODE-rE2G than DE-G pairs corresponding to CTCF elements without H3K27ac (35–50% vs 22%, **Fig. 6c**). This likely reflects that ENCODE-rE2G and other models are designed to identify distal elements that function as enhancers, as opposed to CTCF elements, which are thought to act through distinct mechanisms.

#### Promoter class

ENCODE-rE2G recall was substantially worse for housekeeping genes than for other genes (17% vs 64%, **Fig. 6d**), despite the model including a binary feature indicating which genes are housekeeping genes. This may reflect that this DC-TAP-seq dataset is powered to detect many more weak distal elements for housekeeping genes than the datasets used to train the ENCODE-rE2G model (**Fig. 5d**), and/or that promoter responsiveness appears to be a quantitative, not qualitative, feature of human gene promoters^43^.

Because ENCODE-rE2G was trained on prior CRISPR datasets, we repeated similar analyses using two other models that were not (ABC^16^ and AlphaGenome^46^), and observed similar patterns (**Extended Data Fig. 13)**.

Thus, this CRISPR dataset utilizing random selection design reveals the need for considering indirect effects, quantitative effect sizes, chromatin state, and promoter class in evaluating models. We also suggest that new models trained on well-powered CRISPR perturbation datasets will be needed to predict the effects of the full repertoire of DE-G regulatory interactions, particularly regarding CTCF-bound elements and distal elements with smaller effect sizes.

## Discussion

The functional dissection of distal regulatory elements across the genome remains a central challenge in human genetics. While chromatin annotations and biochemical maps have cataloged millions of candidate regulatory elements across many cell types^3,4,47^, efforts to experimentally connect these elements to their target genes have been more limited. This study introduces a framework for highly powered CRISPRi screens to address this challenge, by combining DC-TAP-seq, unbiased screen design, and tools to consider statistical power. Applying this framework, we conducted a representative survey of 9,666 DE-G pairs in two human cell types.

Our results highlight the importance of statistical power and unbiased element selection for understanding the regulatory genome. Regulatory interactions in DC-TAP-seq had a median effect size of 4.3%—much smaller compared to the effects reported in previous studies (median = 23%, **Fig. 3a**). Indeed, we estimate that previous Perturb-seq experiments failed to detect more than two thirds of the true regulatory interactions tested (**Fig. 3c**). Here, this increased power to detect small effect sizes allowed us to find effects of CTCF sites and distal regulatory elements for housekeeping genes. In parallel studies, we have applied DC-TAP-seq to validate subtle changes in the effects of enhancers upon tuning their 3D contact frequencies^22^ and sensitively detect target genes of enhancers containing risk variants for coronary artery disease^48^. Such small effects appear to be a prevalent feature of human gene regulatory elements, and adequate power will be critical for future studies to understand rules of gene regulation and linking variants to target genes.

The prevalence of regulatory element-gene interactions with small effect sizes raises many questions about the biological significance of such small effects. In our view, this was not an unexpected result. Enhancers appear to be intrinsically compatible with many promoters^43,44,49^, and therefore might have varying effects on different genes in the genome. Many genes are dose-sensitive, and small effects on their expression can impact cellular functions or organismal fitness^50,51^. Robustness is a common feature of regulatory-element gene networks^52,53^, and it is possible that while any individual distal element has a small effect on a gene, their combined effects could be significant. That said, while some of these effects could be functional in the evolutionary sense, others may simply reflect randomness in biochemical interactions that does not impact fitness^54^. Further documenting the full range of element-gene regulatory interactions will be an important step toward building quantitative models to predict gene expression, and, ultimately, understanding the impact on function.

This study provides one of the largest datasets of CRISPRi perturbations of distal elements in a non-K562 cell type (see also ^55^). Reassuringly, DC-TAP-seq data appears very similar between K562 and hiPSCs. While genome-wide accessible elements in the two cell types differ somewhat in their frequencies of certain chromatin signatures (**Extended Data Fig. 9a**), the element-gene pairs with significant regulatory effects identified in the two DC-TAP-seq screens are very similar with respect to their effect sizes (**Fig. 2c**), distance distributions (**Fig. 2b**), and chromatin states (**Extended Data Fig. 9a**). Thus, based on studies of these 2 cell types, the properties of element-gene regulatory interactions we identify here may also apply to others.

Highly powered DC-TAP-seq datasets will be key for evaluating, and perhaps in the future training, large-scale predictive models of distal regulatory interactions. Here, our DC-TAP-seq data provided an early opportunity to evaluate ENCODE-rE2G^29^, a recent predictive model of enhancer-gene regulatory interactions (**Fig. 6**, see also ^30^). We find that ENCODE-rE2G can indeed identify regulatory interactions corresponding to strong enhancers (*i.e.*, >5% effect on expression, marked by H3K27ac). At the same time, our data reveal that 15% of regulatory interactions correspond to CTCF-bound elements that likely act through distinct mechanisms that are not modeled by ENCODE-rE2G (**Fig. 5b, Extended Data Fig. 10)**, for example by altering 3D contacts, and nearly half of random regulatory interactions have small effect sizes (<5%) that were not well represented in the ENCODE-rE2G training dataset. DC-TAP-seq will enable creating additional, larger datasets to enable building new models to detect such effects.

We note several considerations for future studies to apply DC-TAP-seq:

- We expect that the high sensitivity of DC-TAP-seq is useful not only for random surveys of DE-G interactions, as we pursue here, but also for studies to detect the targets of elements containing GWAS variants. This is because causal genes can be expressed at varying levels, and small regulatory effects could be missed unless tested with sufficient statistical power^48^.
- While we designed these DC-TAP-seq experiments to achieve better statistical power than previous studies, we still appear to be underpowered to detect many real regulatory interactions (**Fig. 3c**). Technical factors differing between element-gene pairs, such as cell number, guide efficiency, and gene expression levels, also affect discovery and could be considered in future approaches.
- Our primary analysis used a 20% FDR threshold to define significant regulatory interactions, implying that approximately one in five hits may be a false discovery, which should be taken into account when interpreting regulatory interactions in this study. Future larger studies may be able to apply more stringent thresholds as statistical power increases.
- Our strategy of randomly selecting regulatory elements with DHS signal allowed us to estimate global properties and model performance without biasing element selection to high DHS signal or other features. However, this selection strategy also yielded a total of only 145 regulatory pairs, only ∼52% of which likely represent direct effects. For training improved models, we expect that optimal study design will require substantially larger random datasets, and/or datasets that up-sample likely regulatory interactions, including categories not predicted well by current models. For example, while in our experience targeting regions without DHS or ATAC signal has not impacted gene expression^28^, more comprehensive studies are required to similarly test regulatory elements not marked by DHS peaks.

In summary, we present a CRISPRi-based framework to perturb thousands of candidate regulatory elements and measure their effects on gene expression in a cost-effective and highly sensitive fashion. DC-TAP-seq should now enable generating large-scale datasets to profile millions of candidate regulatory interactions across cell types, and thereby advance our understanding of distal gene regulation and train predictive models to interpret genetic variation.

## Methods

### Construction of sgOpti-CS vector

We constructed the sgOpti-CS vector by replacing the gRNA scaffold of the sgOpti plasmid (Addgene #85681) with the capture sequence containing scaffold from the pBA904 plasmid (Addgene #122238). We digested the sgOpti plasmid with NsiI and EcoRI (New England Biolabs) and used the same enzymes to digest a gBlock (IDT) containing the scaffold with the capture sequence. Following purification, we ligated the cut vector and the digested gBlock with T4 DNA ligase (New England Biolabs), transformed into stable competent cells (New England Biolabs), picked colonies, purified plasmids and sequence verified the final construct.

### Constructs for pilot scale studies

We cloned individual guides targeting the (i) TSS of *GATA1* (2 guides); (ii) an enhancer of *GATA1* (2 guides); (iii) an enhancer of *HDAC6* (2 guides); and (iv) 4 negative control guides in the CROP-HyPR vector backbone. We then combined each plasmid in an equimolar ratio to make the P10 pool. Next, we cloned only the targeting guides used in the P10 pool, added enhancer and TSS guides for *MYC* (2 guides each), TSS guides for *HDAC6* (2 guides) and included 87 negative control guides into the pBA904 vector. We then combined each plasmid in an equimolar ratio to make a 99-guide pool. Next, we cloned the same set of targeting guides used in the 99-guide pool, along with 85 negative control guides, into the sgOpti-CS vector and combined them to make a 97-guide pool.

For pilot studies in WTC11, we cloned the TSS targeting guides for 8 iPSC specific genes into the sgOpti-CS vector. We then combined them with additional sgOpti-CS plasmids with guides targeting the TSS and enhancer of *HDAC6* and 83 negative control guides to make a 98-guide pool. See **Supplementary Table 1** for details.

### Genome build

Coordinates are reported in human genome build hg38 unless otherwise specified.

### Gene reference

Gene definitions are reported in GENCODE version 32 unless otherwise specified.

### Unbiased target selection for CRISPRi DC-TAP-seq screens

Our goal for this design was to select putative *cis-*regulatory elements with as little prior bias as possible. To that end, we decided that the only data that we would use to choose elements would be (1) whether the element is a DNase I hypersensitivity sequencing (DHS) peak, (2) whether it is within a reasonable distance from a gene (<2 Mb), and (3) the gene is expressed at a reasonable level to achieve good power with DC-TAP-seq.

To identify a set of DHS peaks for each cell type, we considered peaks from DHS bed files (K562: Bed - ENCFF185XRG, WTC11 - ENCFF854DSG) using bedtools.

Preliminary analysis of previous datasets^15^ showed that TAP-seq generally has good power to detect a 25% effect on genes with >30 TPM and even higher power to detect a 25% effect on genes with >50 TPM in the perturbed cell line. Therefore, we designed the screen to target elements near at least one gene with 50 TPM. To do this, we filtered genome-wide TPM data for each of the cell types (K562 TPM data from Gasperini et al.^14^; WTC11 data from an in-house generated multiome experiment and identified all genes with at least 50 TPM (**Supplementary Table 9)**. Of those genes, we randomly selected 24 (WTC11) or 25 (K562) of these genes to be “index genes” and created 0.5–4.5 Mb loci

(median 2.4 Mb) around the index gene TSS (**Supplementary Table 3, Supplementary Table 4**). Within these loci, we randomly selected ∼40 DHS elements of any DHS signal strength to be included in the target set. In total, we selected and analyzed 664 and 761 random DHS elements for K562 and WTC11 cells, respectively (**Supplementary Table 3**). We also targeted the promoters of the index gene and some other locus genes with at least 30 TPM expression, for which we also expected to have good power. Promoters were defined as previously described^28^ (https://github.com/broadinstitute/ABC-Enhancer-Gene-Prediction/blob/main/reference/hg38/CollapsedGeneBounds.hg38.TSS500bp.bed).

As positive controls for the K562 screen, we targeted established enhancer-gene pairs from previous CRISPR-based experiments in K562 cells^15,28^, targeting both the promoters and the known distal elements for the gene (**Supplementary Table 3**). For the WTC11 screen, we targeted 7 positive control genes as well as enhancers for 2 genes selected from the literature^56,57^. Finally, we recovered gRNAs targeting 948 and 997 elements in the genome for K562 and WTC11 cells, respectively (**Supplementary Table 3**), although additional non-scoring gRNAs were also included in the final guide pool.

### gRNA library design and cloning

Final gRNA design files for the K562 and WTC11 DC-TAP-seq screens are available on the IGVF Data Portal with accessions **IGVFFI9925XMSV** for **K562** and **IGVFFI6016AIPN** for **WTC11**.

To design gRNAs targeting candidate elements and gene promoters, we first created standard target elements. For candidate elements, we took the summit of DHS peaks (from DHS bed files ENCFF185XRGl for K562 and ENCFF854DSG for WTC11) as the central point and extended 150 bp in either direction, creating a 300 bp window in which to design gRNAs. To target promoters, we used 500 bp elements previously defined as -250 to +250 relative to the transcription start site (TSS) (https://github.com/broadinstitute/ABC-Enhancer-Gene-Prediction/blob/main/reference/hg38/CollapsedGeneBounds.hg38.TSS500bp.bed). We used `liftOver` to map elements from hg38 to hg19 for CRISPR gRNA design and off-target scoring. We then designed 15 independent gRNAs per element according to our established pipeline (https://github.com/EngreitzLab/CRISPRDesigner)^16,21^. As negative controls, we included 400 non-targeting gRNAs that lack matching sequence in the genome and 400 safe-targeting gRNAs that target non-genic sequences with no known open/active chromatin marks. For positive control gene TSS and enhancers in K562 cells, we included the same gRNAs that we used in our previous study^16^ that fall within the 500 bp TSS and the 300 bp enhancer region. In cases with fewer than 15 gRNAs per region, we designed additional gRNAs to supplement. For WTC11, we added 10 additional TSS targeting positive control guides. Overall, we designed 14,968 gRNAs for the K562 and 16,201 gRNAs for the WTC11 CRISPRi screens.

Pools of gRNA oligos containing the guide protospacer sequence flanked by partial human U6 promoter (TATCTTGTGGAAAGGACGAAACACCG) and partial scaffold (GTTTAAGAGCTAAGCTGGAAACAGCATAG) sequences were synthesized by Agilent Technologies. The guide pool was amplified with a forward primer (GGCTTTATATATCTTGTGGAAAGGACGAAACACCG) and a reverse primer (CTAGCCTTATTTAAACTTGCTATGCTGTTTCCAGCTTAGCTCTTAAAC). This amplicon was cloned into the sgOpti-CS vector via Gibson assembly and bacterial electroporation. The plasmid library was sequenced on Illumina MiSeq standard v2 50-cycle kit and shown to include all designed gRNAs with relatively equal coverage of each (there was a 2 fold difference in count frequency between the top and bottom 10th percentiles of gRNAs).

### Gene selection for DC-TAP-seq and primer design

Here, we aimed to create a dataset that would better generalize to the entire genome. As such our only selection criteria for the 49 loci targeted across K562 and WTC11 were that they contain at least one gene (which we term the ‘index gene’) with ≥50 transcripts per million (TPM) in the tested cell type.

For DC-TAP-seq in K562 cells, we designed primers to read out the expression of the following genes: 1) 25 index genes of our random loci, 2) 195 locus genes in our random loci - these include genes that were not targeted with gRNAs, 3) 33 positive control genes, 4) 11 genes for which we have validated primers from our pilot scale study or from our previous study^15^, and 5) 39 housekeeping genes with 30– 150 TPM that were not targeted for perturbation and whose expressions are not expected to change. In total, we designed primers for 303 genes (**Supplementary Table 4)**.

For WTC11 cells, we designed primers to read out the expression of the following genes: 1) 24 index genes of our random loci, 2) 138 locus genes in our random loci, including genes that were not targeted with gRNAs, 3) 8 positive control genes, and 4) 33 housekeeping genes with 30–150 TPM that were not targeted for perturbation and whose expressions are not expected to change. In total, we designed primers for 203 genes (**Supplementary Table 4)**.

Primers were designed using a pipeline adapted from the original TAP-seq paper^15^, with minor application-specific modifications. The pipeline positions primers 300–500 bp upstream of the polyadenylation site and checks for off-target binding and primer dimer formation. We used Gencode transcript annotations and scRNAseq data from the corresponding cell type. Following the initial run, we manually inspected primer landing sites to ensure that the primers mapped to annotated exons and proximal to the 3’ scRNA-seq signal. In cases where the tool misannotated polyA (pA) sites, we edited the pA site call based on the scRNA-seq signal and reran the pipeline. We additionally assessed multiplex PCR compatibility across the final set using Primer 3 complementarity metrics (QC integrated in the design pipeline). Following the in-silico design we tested the primer panel for each cell type in pilot DC-TAP-seq experiments to make sure we see all the genes being amplified. We acknowledge that targeted amplification can introduce gene-specific differences in capture or amplification efficiency. Nevertheless, the absolute gene expression correlation between TAP-seq and whole-transcriptome scRNA-seq ranges from 0.56 to 0.86^15^. Because comparisons were made between perturbed and control cells using the same primer panel within each cell type, any primer-specific effects should be consistent across experimental conditions and are therefore unlikely to systematically bias differential expression estimates.

### Cell culture

We grew K562 cells harboring doxycycline-inducible KRAB-dCas9-IRES-BFP (lentivirally integrated, AddGene Plasmid #85449)^16,28^ in RPMI-1640 (Thermo Fisher Scientific, Waltham, MA) with 10% heat-inactivated FBS (HIFBS, Thermo Fisher Scientific), 2 mM L-glutamine, and 100 units/ml streptomycin and 100 mg/ml penicillin. We maintained K562 cells at a density between 100,000 and 1 million per mL.

We seeded human iPS cells (WTC11) harboring doxycycline-inducible KRAB-dCas9-P2A-mCherry^58^ (gift from Ioannis Karakikes, Stanford University) in 6-well plates coated with 1:100 LDEV-Free Reduced Growth Factor Basement Membrane Matrix (Thermo Fisher Scientific). The culture medium used was StemFlex Medium supplemented with 10% StemFlex supplement (SF complete media, Thermo Fisher Scientific), 2 µl of Normacin per ml of media (1:500 final, Invivogen), and 0.4 µl of 10 mM Rock Inhibitor (Y-27632 Dihydrochloride, STEMCELL Technologies). After 24 hours of seeding, we continued to maintain the cells with SF complete media without Rock Inhibitor and passaged them with 1 ml of accutase (STEMCELL Technologies) as necessary, aiming to maintain an 80-90% confluence to prevent stacked cell growth.

We maintained HEK293Ts between 20 and 80% confluency in DMEM with 1 mM Sodium Pyruvate, 25 mM Glucose (Thermo Fisher Scientific) and 10% HIFBS.

For all CRISPRi experiments, we read out effects on expression after activating CRISPRi for 48 hours with doxycycline. We selected this timepoint due to extensive previous experiments characterizing and applying CRISPRi in large-scale screens to study regulatory elements^16,21,28,33,38^. These studies have shown that, at least at the 48-hour timepoint, (i) the strongest-scoring gRNAs are located at the center of the accessible site; (ii) there is little evidence of H3K9me3 spreading beyond the targeted accessible chromatin site by ChIP-seq^16,36,59^; (iii) the quantitative effects of CRISPRi perturbations match those of genetic deletions of the same elements^36,60^; and (iv) computational models trained on these CRISPRi data perform well at predicting effects of genetic perturbations in the form of eQTLs and GWAS variants^16,21,29,30^.

### Lentivirus production

We plated 550,000 HEK293T cells on 6-well plates (Corning, Corning, NY) and 24 hours later transfected with 360 µg pMD2 g, 900 ng psPAX2, and 1.2 µg transfer plasmid using FuGENE (FuGENE Transfection, Middleton WI). For pooled screens, the cell number and plasmid mass were scaled proportionally to 14 million cells on a 15 cm plate (Corning). 16 hours post-transfection we changed the media to DMEM with 20% HIFBS. At 48 hours post-transfection, we harvested viral supernatants and filtered them through a 0.45 µM syringe filter. We aliquoted the lentivirus in cryovials and stored at -80°C.

For WTC11 infections, we concentrated the virus using Lenti-X Concentrator (Takara Bio, 631231). The harvested virus was mixed 3:1 (virus to Lenti-X Concentrator) and incubated for 1 hour at 4°C before spinning for 45 min at 1,500 x g. The supernatant was discarded and the virus pellet was resuspended in 1/10th volume with PBS and immediately used for infection.

### Lentivirus transduction and cell harvest

For lentivirus transduction of K562 cells, we plated enough cells for at least 1000x library coverage in a 12-well plate and added varying volumes of virus in the presence of 8 mg/mL of polybrene. We performed spinfection by centrifuging the plates at 1200g for 40 minutes at 37°C and transferred the plates to the incubator. After 24 hours, we pelleted and resuspended the cells in media containing 1 µg/mL puromycin. We selected cells for 48 hours with puromycin and performed cell count and viability check to confirm that the non-infected control cells are about 50% dead. We incubated the cells in fresh selection media for additional 48 hours and confirmed the non-infected control cells are all dead. From this point, we cultured the infected cells in media containing 0.3 µg/mL puromycin, maintaining 1000x library coverage when passing. In parallel, we collected ∼1M cells for ddPCR analysis. Once the cells were stabilized and exhibited normal doubling time, we treated the cells with 1 µg/ml Doxycycline for 48 hours. We then harvested cells to use directly for DC-TAP-seq or frozen cells as stock. To freeze cells, we centrifuged the cells and resuspended the resulting cell pellet in Bambanker cryopreservation media (Wako Chemicals) at a density of 1M cells per ml. We stored the cryovials in a Mr Frosty Freezing Container (Thermo Fisher Scientific) at -80°C for 24 hours before transferring for long-term storage in liquid nitrogen.

For lentivirus transduction of WTC11 cells, we plated 250,000 cells per well in 6-well plates using SF complete media containing Rock Inhibitor (RI). After 24 hours, we replaced the media with prewarmed SF complete media without RI and added 1 ug/mL of polybrene (Sigma-Aldrich) to transduce the cells. Following a 15-minute incubation at 37°C, we removed the plates and infected the cells with varying amounts of virus to achieve different MOIs. We then spun the plates at 1200 x g for 40 minutes at 23°C before placing them back in the incubator to allow the cells to recover overnight. After 24 hours of infection, we replaced the virus-containing media with prewarmed SF complete media without Rock Inhibitor and supplemented with 0.3 µg/mL of puromycin. For the pilot-scale experiment, we transduced 400,000 cells, and for the large-scale experiment, we transduced approximately 2 million cells (to maintain a 750-1000x coverage at high MOIs). We removed the media 24 hours later and added fresh SF complete with 0.3 µg/ml puromycin. We maintained the cells for an additional 72 hours in the selection media, after which we collected the cells from each viral infection and re-seeded 1 million cells per well in 3 wells of a 6-well plate for the pilot or 3.2M cells for the large-scale experiment in SF complete with RI (no puromycin) and 1 µg/ml Doxycycline. At the same time, we collected ∼500,000 cells from each infection for ddPCR analysis. Upon determining the MOI we continued with the cells that were infected at the desired MOI(s). After 24 hours of Doxycycline induction, we removed the media and added fresh SF complete with 1 µg/ml Doxycycline without RI. After an additional 24 hours (48 hours total of Doxycycline induction), we collected the cells and centrifuged them for 5 minutes at 23°C at 350 x g. We removed the media and resuspended the resulting pellet in Cryostor cryopreservation media (STEMCELL Technologies) at a density of 2 million cells per ml and aliquoted into cryovials. We then transferred the vials to a Mr Frosty Freezing Container (Thermo Fisher Scientific) and stored at -80°C for 24 hours before transferring for long-term storage in liquid nitrogen.

### Digital droplet PCR

We used digital droplet PCR (ddPCR) using primers targeting the gRNA lentivirus to estimate the multiplicity of infection for each experiment.

We purified genomic DNA (gDNA) with the QIAmp DNA blood Mini column extraction kit (Qiagen, 51104) following manufacturer instructions. We eluted the gDNA in 100 ul of nuclease free water. We digested >400 ng of the purified gDNA with HindIIIHF (NEB) enzyme with a concentration of 0.4 U/ml (1ul) in a total reaction mixture of 50 ul at 37°C for 1 hr and then another 15 minutes at 80°C for inactivating the enzyme. We diluted gDNA samples with water to obtain a gDNA concentration of 2.5 ng/ul.

To prep the samples for ddPCR we made 24 µl of reaction mix with 10 µl of the purified gDNA (25 ng), 12 ul of the ddPCR Supermix for probes, no dUTP (Bio-Rad, 1863024), 1 µl of 22.5 uM primer mix containing 2 primer pairs (target and reference) and 1 µl of 0.6 uM probe mix containing 2 Taqman probes (target and reference). For droplet generation we then loaded Bio-Rad’s ddPCR chip following manufacturer’s instructions by securing chip in the holder and then loading 20 ul of sample into middle channels and 70 ul of Droplet Generation Oil for Probes in bottom channels. We used 50% glycerol for empty wells so the chip will not run dry. We covered the chip with a provided gasket and started the droplet generation run. Once the run was completed, we transferred 40 ul of the generated emulsion into a 96 well plate, we covered the plate with one sheet of pierceable foil, and sealed the plate at 180°C for 5 s. We then transferred the plate to a thermocycler (within 30 minutes of droplet generation) and cycled at 95°C for 10 min for 1 cycle; 40 cycles at (94°C for 30 s, 60°C for 60 s), 1 cycle at 98°C for 10 min, and cool to 4 °C. After cycling, we removed the plate and stored it at 4°C overnight, before transferring to the QX200 droplet reader (Bio-Rad) to read the droplet fluorescence using Quantasoft software.

### Cell preparation for DC-TAP-seq

For K562 screens, we used both fresh and frozen cells to perform DC-TAP-seq. For fresh cells, we harvested ∼1M cells into a tube. For frozen cells, we thawed a vial in a 37°C water bath until a small frozen cell pellet remained and gently added the cells into 5 ml media. We centrifuged both the fresh and frozen cells at 23°C at 200 x g for 3 minutes and washed the cells by resuspending the cell pellet in 1 ml PBS + 0.04% BSA and re-centrifuging. We resuspended the cell pellet in 1 ml PBS + 0.04% BSA and filtered the cells with a 70 µm Flowmi Tip Strainer (Millipore) to achieve a cell concentration of 700-1200 cells/µl. We then obtained an accurate cell concentration by measuring with a hemocytometer and kept the cells on ice until loading on a 10x chip.

For WTC11, we used fresh or thawed cells from frozen stocks to perform DC-TAP-seq. We added Accutase to the fresh cells and quenched with 2 ml of SF complete for every ml of accutase to harvest and spun at 200 x g for 5 minutes at 23°C. We removed the supernatant and washed the cells with 1x PBS + 0.04% BSA. Finally, we resuspended the cells in 1x PBS + 0.04% BSA to achieve a density of 1000 cells/µl, which was used for loading on the 10x Genomics Chip wells. For frozen cells, we thawed a vial in a 37°C water bath for approximately 1 minute and quenched the thawed cells with 9 ml of SF complete media by slowly adding the first 1 ml into the vial. We then centrifuged the cells at 350 x g for 4 minutes, removed the supernatant, and resuspended the cell pellet in 1 ml of SF complete media. We centrifuged the cells for 4 minutes at 23°C at 350 x g, removed the supernatant and washed the cells in 2 ml of 1x PBS + 0.2% BSA. We finally resuspended the pellet in 1x PBS + 0.2% BSA to achieve a cell concentration of 1000 cells/µl and kept the cells on ice until loading on the 10x Genomics chip.

### DC-TAP-seq library preparation

A detailed protocol for DC-TAP-seq can be found in protocols.io^61^. We acquired cells encapsulated into droplets using the Chromium Next GEM Single Cell 3’ Kit V3.1 (10x Genomics) following manufacturer instructions. For loading the cells into the 10x channels, we used the recommended cell suspension numbers at an approximate loading of 16,500 - 20,000 cells per lane. We followed the 10x 3’ reagent kit v3.1 protocol through the GEM-RT incubation step until the cDNA clean-up and elution steps.

For PCR 1 (all-in-one PCR) we used 35 µl of the purified cDNA from each lane, 2.5 µl 100 uM gene-specific outer primer mix, 4 µl 10 uM Partial Read1, 1 µl 40 uM Partial Read 1N, 1.2 µl 100 uM Partial TSO and 50 µl KAPA HiFi-Hotsart Readymix in 100 µl total volume. We set up the PCR at 95 °C for 3 min for one cycle; 11 cycles at [98 °C for 20 s, 67 °C for 60 s, 72 °C for 60 s], one cycle at 72 °C for 5 min, and cooling to 4 °C. We cleaned up the samples using double sided SPRI where we first added 65 µl SPRIselect beads (Beckman Coulter) to 100 µl of the PCR reaction and transferred the supernatant (containing smaller/gRNA amplicons) into a fresh tube. We then washed the pellet (containing larger/mRNA amplicons) twice with 80% ethanol and eluted in 40 µl of Buffer EB (Qiagen). For the supernatant (guide amplicon), we added 55 µl of additional SPRI beads followed by two 80% ethanol washes and eluted in 40 µl of Buffer EB. We further cleaned the guide amplicons with 40 µl of SPRIselect beads followed by two 80% ethanol washes and finally eluted in 50 µl of Buffer EB. We obtained a yield range of 280-4000 ng for both mRNA and guide after PCR1.

For mRNA PCR 2, we first diluted all PCR1 products to 2ng/ul. We used 10ng of PCR1 cleanup (5ul), 2.5 µl 100 uM gene-specific inner primer mix, 4 µl 10 uM Partial Read1, and 50 µl KAPA HiFi-Hotsart Readymix in 100 µl total volume. We set up the PCR at 95 °C for 3 min for one cycle; 8 cycles at [98 °C for 20 s, 67 °C for 60 s, 72 °C for 60 s], one cycle at 72 °C for 5 min, and cooling to 4 °C. We cleaned up the samples using 150 ul ofSPRIselect where we first added 80 µl SPRIselect beads to 100 µl of the PCR reaction followed by two 80% ethanol washes and eluted in 30 µl of Elution buffer. For the guide PCR 2, we followed the Chromium Next GEM Single Cell 3’ Kit V3.1 CRISPR Screening Library Construction instructions by adding 50ul of 10x amp mix (PN 2000047) and 45 ul of 10x Feature SI Primers 3 (PN 2000263) to 5ul of the guide PCR1 cleanup for a total reaction volume of 100ul. We set up the PCR at 98 °C for 45 s for one cycle; 11 cycles at [98 °C for 20 s, 58 °C for 5 s, 72 °C for 5 s], one cycle at 72 °C for 1 min, and cooling to 4 °C. We cleaned up the samples by adding 100µl of SPRIselect beads to 100 µl of the PCR reaction. We then washed the pellet twice with 300ul of 80% ethanol and eluted in 30 µl of Buffer EB. We obtained a yield range of 240-1080ng for both mRNA and guide after PCR2.

For mRNA PCR 3, the indexing PCR, we used 2.5 moi ul of a 10 uM stock custom p7 primer unique to each sample and 4ul of a common 10uM stock of a p5 primer, along with 50ul of the KAPA HiFi Hotstart Readymix. We set up the PCR at 95 °C for 3 min for one cycle; 8 cycles at [98 °C for 20 s, 60 °C for 15 s, 72 °C for 45 s], one cycle at 72 °C for 5 min, and cooling to 4 °C. We cleaned up the samples using 150 ul of SPRIselect where we first added 80 µl SPRIselect beads to 100 µl of the PCR reaction followed by two 80% ethanol washes and eluted in 30 µl of Elution buffer. For the guide index PCR, we continued to follow the 10x CRISPR Screening Library Construction manual by adding 50ul of 10x amp mix (PN 2000047) and 25ul of Buffer EB to 5ul of cleaned up PCR 2 products with 20 μl of an appropriate sample index from the 10x DualIndex Plate NT Set A (PN 3000483) for a total reaction volume of 100ul. We cleaned up the samples using double sided SPRI where we first added 80 µl SPRIselect beads to 100 µl of the PCR reaction and transferred 170ul of the supernatant into a fresh tube. We then added 20 µl of additional SPRI beads to the saved supernatant and followed by two 80% ethanol washes and eluted in 30 µl of Buffer EB.We obtained a yield range of 510-2010ng for both mRNA and guide after PCR3.

To obtain size distribution, concentration and to assess the quality of our final libraries we obtained Bioanalyzer traces using the dsDNA high sensitivity kit (Agilent). We pooled and sequenced the final guide and mRNA libraries using NextSeq 500/550 or NovaSeq 6000 or NovaSeq X Plus (Illumina) with paired-end reads of 28 cycles for read 1, 90 cycles for read 2, and 8 cycles for both index 1 and 2.

### Single guide RT-qPCR validation

We selected a single guide for each target to compare the knockdown efficiency of the same guide as calculated using DC-TAP-seq. List of targeting gRNAs and RT-qPCR primers for amplified genes can be found in **Supplementary Table 10**. We used 2 non-targeting guides; NC1 (GATCGCGAGGACCCGTTCCGCC) and NC2 (GACTCGTCACATGGGGTTGCGA) as negative controls. Individual guides were cloned in sgOpti-CS and were infected in K562 or WTC11 lines by lentiviral transduction at low MOI. Following puromycin selection, we grew the cells in replicates. Cells were induced with 1 µg/ml Doxycycline for 48 h and harvested for RT-qPCR in buffer RLT (Qiagen). We extracted RNA using Dynabeads MyOne Silane beads (Thermo), treated samples with TURBO DNase (Thermo), and cleaned again with Dynabeads MyOne Silane beads. We used Superscript IV reverse transcriptase and random hexamers with oligo dT to convert the RNA to cDNA. We performed qPCR with SYBR Green I Master mix (Roche) with primers for genes listed in **Supplementary Table 10** along with *GAPDH* (Forward = AGCACATCGCTCAGACAC, Reverse = GCCCAATACGACCAAATCC), and calculated the differences using the ΔΔ Ct method.

### Single guide CTCF ChIP-seq

We selected 4 guides corresponding to two CTCF-bound elements with no H3K27ac signal, and >10% negative effect sizes in WTC11. List of guides can be found in the sheet CTCF_guides in **Supplementary Table 10.** We used 2 non-targeting guides; NC1 (GATCGCGAGGACCCGTTCCGCC) and Crop_15807 (GGCTCACGTAGTCCGTTATTA) as negative controls. Individual guides were cloned in sgOpti-CS and were infected in WTC11 lines by lentiviral transduction at low MOI. Following puromycin selection, we grew the cells in three independent cultures (i.e. biological triplicate) . Cells were induced with 1 µg/ml Doxycycline for 48 h and harvested for ChIP-seq following ENCODE protocols (antibody to CTCF Diagenode: part number is C15410210, lot number is A2359-00234P. All CTCF ChIP-seq protocols are available at the ENCODE portal: https://www.encodeproject.org/chip-seq/transcription_factor/. FASTQ files were processed using the ENCODE ChIP-seq pipeline (v2; https://github.com/ENCODE-DCC/chip-seq-pipeline2). Filtered BAM files generated by the pipeline were converted to bedGraph format using bedtools genomecov^62^, with the signal scaled by sequencing depth (bedtools genomecov -ibam ∼{bam}-bg -scale ∼{scale_factor} -pc | sort -k1,1 -k2,2n > ∼{sample_name}.bedGraph). The resulting bedGraph files were subsequently converted to bigWig format using the UCSC bedGraphToBigWig utility^63^. CTCF signal intensity at the regions of interest was quantified using the pyBigWig Python library (source code available on GitHub). CTCF motifs were annotated using the JASPAR track from the UCSC Genome Browser (hg38; https://hgdownload.soe.ucsc.edu/gbdb/hg38/jaspar/JASPAR2024.bb), specifically filtering for the CTCF motif ID MA0139 (source code available on GitHub).

Statistical analysis of CTCF ChIP-seq data: Each sgRNA was therefore compared using n = 3 libraries against n = 6 control libraries. The control group combines the three NC2 (negative control 2) and three NT1 (non-targeting) libraries, one value per library, without averaging or weighting. All comparisons used a two-sided Wilcoxon rank-sum test with the exact null distribution (no ties); asymptotic p-values lead to the same conclusions. We did not correct for multiple testing as each sample has an individual gRNA, thus each test is an independent event.

### Single-cell RNA-seq data processing with Cell Ranger

We aligned reads to transcripts and identified cells using the 10x Cloud Analysis web browser version of Cell Ranger Count v7.0.0 (pilot K562 DC-TAP-seq) or v7.0.1 (large-scale K562 DC-TAP-seq) or v7.1.0 (pilot K562 TAP-seq and large-scale WTC11 DC-TAP-seq) or v7.2.0 (pilot K562 scRNA-seq) or v9 (validation screen). For all the analysis except for the validation screen, we input our transcriptome FASTQ files as Targeted Next GEM 3’ Gene Expression libraries with a reference file containing the bait sequences of the genes for which we designed primers for. We generated this reference file by subsetting the bait sequences found in the 10x Genomics `human_pulldown_probes.csv` file (built on GRCh38-2020-A). The transcriptome reference used for Cell Ranger analysis was GRCh38-2020-A (hg38, GENCODEv32). For the guides, we input the guide FASTQ files as CRISPR Guide Capture libraries and the feature_reference file containing the guide sequences. The result of this analysis is a filtered featured barcode matrix containing cell barcodes corresponding to real cells (as opposed to empty droplets) with UMI counts corresponding to every gene in the targeted transcriptome and every guide in the guide library.

### Comparing guide capture between vectors

To compare overall guide capture efficiency, we performed the following pilot experiments in K562: (i) TAP-seq^15^ (amplifying 97 genes) with the P10 guide pool that was cloned in the CROP-HyPR vector (CROP-seq derived guides); (ii) DC-TAP-seq (amplifying 97 genes) with the 99-guide pool cloned in the pBA904 vector (mouse U6 promoter); (iii) DC-TAP-seq (amplifying 97 genes) with the 97-guide pool cloned in the sgOpti-CS vector (human U6 promoter). For each experiment, we took the average of the total guide UMIs captured in every single cell relative to the number of guides detected in that cell. To compare across experiments, we then divided this number by the average reads per cell and multiplied by 1000 to get the average UMI captured per guide per cell per 1000 reads.

### Comparing effect sizes in different MOI experiments

To understand whether knockdown effect size is impacted by MOI, we compared DC-TAP-seq experiments amplifying 97 genes in K562 performed at different MOIs using the 97-guide pool cloned in the sgOpti-CS vector. We focused on four guides targeting either the *GATA1* TSS (2 guides) or known *GATA1* enhancers (2 guides), and measured their effects on *GATA1* expression. We first averaged the effect sizes for each guide on *GATA1* expression across two biologically independent experiments done at low MOI (MOI 1). We then used these averages to normalize the effect sizes of each guide at higher MOIs (MOI 6, 9, 14 and 20). A normalized value of 1.0 indicates the same magnitude of GATA1 knockdown as the average effect observed at MOI 1. For WTC11, we followed the same general approach, though the low MOI (MOI 1) data came from a single experiment. In this dataset, we assessed the impact of promoter perturbations (with TSS targeting guides) on the expression of their corresponding genes—specifically: *BAG3*, *NANOG*, *ROCK1*, *SOX2*, and *YBX1*. Effect sizes at higher MOIs were normalized to the effect size observed for each promoter-gene pair at MOI 1. In these experiments, an MOI>10 reduces knockdown effect size. We also performed two validation studies in WTC11 which similarly support this conclusion. We compared DC-TAP experiments for 30 elements (353 DE-G pairs) at various MOI (12.6, 14.4, 1.1 & 1.5) and compared these to the large scale WTC11 screen. While all MOI produce remarkably concordant results (R=0.91-0.97), there is marked decrease in knockdown effect size with an MOI >10.

### UMI capture complexity analysis

To calculate the mean UMIs per cell per gene captured at various sequencing depths between DC-TAP-seq and Perturb-seq, we performed two experiments in K562 cells. In the DC-TAP-seq experiment, we amplified 97 genes upon infecting the cells with the 97-guide pool. In the Perturb-seq experiment, we performed 10x Genomics 3’ scRNA-seq for the whole transcriptome after infecting cells with a 16-guide pool cloned in pBA904 (12 targeting + 4 negative control guides from the 99-guide pool). We then performed a downsampling analysis on the `molecule_info.h5` files generated by Cell Ranger from these two experiments. The total reads across all genes and cells in each study were downsampled at various fractions (DC-TAP-seq: 0–1; Perturb-seq: 0–0.1, due to the much higher total amount of reads in the Perturb-seq dataset). This allowed us to compare, at matched read depth per cell, the number of UMIs captured for the 93 detected genes (out of the 97 amplified genes) in both the assays. We calculated the mean UMIs counts per cell per gene for these genes at each fraction of reads.

### Data analysis for DC-TAP-seq experiments

We analyzed DC-TAP-seq data for the large random experiments using the following overall procedure:

1. We defined a set of candidate element-gene pairs for differential expression testing, connecting all elements with all tested genes within 2 Mb (in hg19)
2. We conducted differential expression analysis for each element-gene pair, using SCEPTRE (effective FDR < 20%)
3. We computed statistical power for each element-gene pair, using a simulation-based procedure (see next section)
4. We re-annotated all element-gene pairs to distinguish “promoters” (*i.e.*, promoters and promoter-proximal elements that overlap or are within 1 kb of annotated promoters) and “distal elements” (*i.e.*, tested elements that do not overlap an annotated promoter), and exclude from consideration elements in gene bodies of the tested gene.

These analyses are implemented in https://github.com/EngreitzLab/DC_TAP_Paper/tree/v2.0.3/workflow. The final output of this pipeline is a comprehensive table of all tested element-gene pairs for both K562 and WTC11, annotated with: genomic coordinates (hg38 and hg19) of the element, target gene information, SCEPTRE p-value and effect size with confidence intervals, power simulation results for a range of effect sizes between 2% and 50%, overlap flags (promoter/gene-body), and category labels for each pair. This table **(Supplementary Table 3)** constitutes the foundation for all reported figures and conclusions regarding element-gene connectivity in our DC-TAP-seq screens.

Each step is explained in more detail below:

#### 1. Creating discovery pairs for SCEPTRE input: testing all element-gene pairs within 2 Mb

We created a set of distance-matched element-gene pairs to test for differential expression. Regardless of element type (promoter, candidate cis-regulatory element, etc.), the distance from the center of each targeted element to the transcription start site of all genes sequenced via TAP-seq were computed using `gencode.v32lift37.annotation.gtf` (see Data Availability section). Pairs were selected for testing if the distance was <2Mb. In our primary analysis, we tested 5,059 and 4,607 randomly selected distal elements (excluding positive controls, marked in **Supplementary Table 3** in columns selfPromoter, Positive_Control_DistalElement_Gene & Positive_Control_selfPromoter) for differential expression in K562 and WTC11, respectively, with a false discovery rate of 20%. In a secondary analysis for backward compatibility with other benchmarking analyses described elsewhere^29,30^, we tested 7,475 and 6,574 element-gene pairs (inclusive of randomly selected distal elements, promoters, and positive controls) for differential expression in K562 and WTC11, respectively, with a false discovery rate of 10%.

#### 2. Differential expression analysis with SCEPTRE: gRNA assignment, quality control, calibration

We then analyzed our set of element-gene pairs for differential expression using SCEPTRE v0.10.1^37^. The SCEPTRE object was initialized with high MOI settings, and was provided with the gene counts matrix, guide capture matrix, and “10x lane” as a covariate. All other parameters were SCEPTRE defaults (see https://katsevich-lab.github.io/sceptre/). We then performed gRNA assignment with SCEPTRE, thresholding assignment at 5 UMIs. We filtered both single cells and element-gene pairs based on SCEPTRE’s default settings, and verified calibration on negative controls using SCEPTRE’s `run_calibration_check()`.

Each gRNA was assigned to a mean of 47.07 cells in K562 and 56.38 cells in WTC11 with an MOI of 7.34 and 5.07 in K562 and WTC11 respectively. After filtering element-gene pairs, we were left with 7,493 pairs in K562 and 6,574 pairs in WTC11. Cell-wise filtering retained 93,291 out of 95,997 cells for K562 and 175,402 out of 180,116 cells for WTC11. Calibration check of the negative controls resulted in 3 out of 7,493 and 2 out of 6,574 false discoveries in K562 and WTC11 respectively. In our primary analysis, we found 113 and 172 significant element-gene pairs in K562 and WTC11, respectively (excluding positive controls FDR = 20%, **Supplementary Table 3**). In our secondary analysis, we found 258 significant hits in K562 and 237 significant hits in WTC11 (FDR = 10%, **Supplementary Table 3**).

#### 3. Statistical power analysis

We used a simulation-based method to compute statistical power for each element-gene pair (see following section). **Supplementary Table 3** includes power to detect different effect sizes (2%, 5%, 10%, 15%, 20%, 25%, 50%).

#### 4. Annotation and filtering of element-gene pairs

We re-annotated which elements in the screen overlap gene bodies and promoters. We started with the gRNA design files (in hg19). All genomic features were defined on GENCODE v32lift37 (https://www.gencodegenes.org/human/release_32lift37.html), consistent with the hg19 build used for the guide design.

We annotated “promoters” as elements located within 1 kb of the TSS of any protein-coding transcript, or that overlap the set of promoters previously defined by ABC^16^. All other elements were annotated as “distal”.

We annotated elements as overlapping a gene body by overlapping each element with the span of the first to last exon of each gene. Element-gene pairs involving an element that overlaps the gene body of the tested gene were excluded from further analysis, as in previous studies, due to the confounding effects of CRISPRi in gene bodies^16,28,38^.

After labelling with genic features, all element regions were lifted to hg38 using UCSC’s `liftOver`. Elements whose coordinates failed liftOver were discarded. In the K562 experiment, one element, chr8:145537346-145537700, was removed due to a partial deletion from hg19 to hg38.

Finally, based on these annotations, we defined the following categories of element-gene pairs for further analysis:

- **Distal element-gene pairs:** All pairs in which the element does not overlap a promoter, and where the element is not located in the gene body of the target gene
- **Positive control distal element-gene pairs:** A subset of distal element-gene pairs that were deliberately included as positive controls in our screen design
- **Random distal element-gene pairs:** A subset of distal element-gene pairs that correspond to elements selected through the random design process described above.
- **Promoter-gene pairs:** This category includes any pair where the perturbed element overlaps a promoter region defined above.
- **Self promoters.** A subset of promoter-gene pairs where the perturbed promoter is paired with its own gene. In other words, these pairs test whether perturbing a gene’s promoter affects the expression of that same gene
- **Distal promoter-gene pairs:** A subset of promoter-gene pairs where the perturbed promoter corresponds to a different, distal gene. In other words, these pairs test whether the promoter of one gene affects the expression of another nearby gene.
- **Positive control promoter-gene pair:** A subset of self promoters where the gRNAs were included as positive controls. These elements are expected to lead to strong decreases in gene expression.

After automatically annotating promoters and distal elements as above, we manually reclassified two specific element-gene pairs based on *post hoc* examination: (1) We reclassified *RTN4_TSS_8* reclassified as a distal element. This was originally designed as a promoter-targeting control, because it overlaps the TSS of a transcript of the *RTN4* gene). However, upon inspecting genome tracks indicative of active transcription in K562, we found that the particular transcript whose promoter *RTN4_TSS_8* overlaps is not expressed. In essence, *RTN4_TSS_8* is overlapping a “promoter” that is inactive in this cellular context. We therefore reasoned that perturbing this region might behave more like a distal element perturbation (since the active RTN4 promoter in K562 is elsewhere or the gene is lowly expressed). We reclassified *RTN4_TSS_8* as a distal element in our categorization. Consistent with its inactivity, perturbing *RTN4_TSS_8* did not produce a significant expression change in any nearby gene in K562. (2) We reclassified chr8:130594299–130594600 paired with *CCDC26* as a positive control distal element-gene pair. One targeted element on chromosome 8 was initially removed by our automated filters because it overlaps the gene body of *CCDC26*, the gene it was intended to regulate. Overlap with the target gene’s body is usually a disqualifying confounder due to potential interference with transcription. However, upon closer investigation we discovered that *CCDC26* has two transcript isoforms, and the isoform that overlaps this element is not expressed in K562. Moreover, this element was included in our design as a positive control distal enhancer, having shown regulatory activity in prior CRISPR screens. Therefore, marked this element as a positive control distal element-gene pair.

All element-gene pairs were subjected to the same power filtering for non-significant results: if a pair was not significant and underpowered (<80% power at 15% effect), it was noted as such. In our final summaries, we report the number of significant pairs in each category (further broken down by direction of effect, up or down) and the number of non-significant pairs that are well-powered vs. underpowered in each category. These summaries (**Supplementary Table 6**) give an overview of the screen’s outcomes stratified by regulatory category (distal element vs promoter, etc.) for both K562 and WTC11.

### Uniformly processing external enhancer perturbation datasets

To compare results from the K562 and WTC11 DC-TAP-seq screens to other published enhancer perturbation datasets in Figures 1-3, we re-analyzed and integrated 4 published CRISPR enhancer Perturb-seq perturbation screens in K562^19,20,27^. Below, we describe the processing steps applied to each dataset.

### Processing of Perturb-seq datasets

We reprocessed data for all datasets using the methodologies described in the original publications. This included standard preprocessing steps: cell quality filtering based on UMI counts, mitochondrial transcript fraction, and gene detection thresholds specific to each dataset. For each dataset, we generated three core data matrices: (1) an mRNA gene expression counts matrix (genes × cells), (2) a guide RNA counts matrix (guides × cells), and (3) an annotation file describing each guide and its target element. To ensure consistent evaluation across all datasets, we defined a unified set of candidate elements using the ABC pipeline^16^ for K562 DNase-seq data (ENCODE experiment accession ENCSR000EOT). Only guides that overlapped with these candidate elements were retained for downstream analysis. The preprocessing details of the raw data for each Perturb-seq dataset is described below.

### Processing of CRISPRi-Perturb-seq data from Gasperini *et al.*, 2019^14^

Processed data containing UMI counts per cell and gene, and detected guides per cell were downloaded from GEO (accession: GSE120861). In concordance with the analysis in Gschwind et al., 2023, we used the guide assignment matrix reported there and adopted GENCODE v26lift37 as the reference annotation. Briefly, we recomputed guide-to-element assignments for our analysis as follows: Genomic coordinates of guideRNA binding sites were inferred using BLAT to align guide sequences to the hg19 genome sequence. Guides were assigned to targeted elements by overlapping guide binding sites with the K562 candidate elements as described above and required input files for SCEPTRE were generated^37^. Coordinates were then lifted to hg38 using the same procedure as for our DC-TAP-seq datasets.

### Processing of CRISPRi-Perturb-seq data from Morris *et al.*, 2023

Data for the STING-seq v1 and v2 screens from Morris *et al.*, 2023^19^ were analyzed separately and differential expression results were combined into one dataset. Data from the individual screens were processed as follows:

#### STING-seq v1 screen

We retrieved the data for the STING-seq v1 screen (the smaller pilot screen from the study) from GSE171452, including count matrices for cDNA, cell-hashing (HTO), and guide-barcode (GDO). A Seurat object was initialised with the cDNA matrix; HTO and GDO matrices were added as separate assays. Cells were retained if their unique-gene count, total cDNA UMI count and mitochondrial transcript fraction lay between the 15th–99th, 20th–99th and 5th–90th percentiles, respectively. HTO counts were centre-log-ratio normalized and demultiplexed with HTODemux (positive.quantile = 0.99); only singlets were retained. The guide sequences were validated using UCSC’s BLAT API, as the modest scale of the guide library permitted API-based validation without server limitations, with requirements for 100% sequence similarity to unique genomic locations. The guides were then overlapped with the K562 candidate elements described above.

#### STING-seq v2 screen

We retrieved the raw data for the STING-seq v2 screen (the larger, full-scale screen from the study) from GSE171452. We performed quality control separately for each lane: cells were retained if their unique-gene count lay between the 1st-99th percentiles, total cDNA UMI count between the 10th–99th percentiles, and mitochondrial transcript fraction between the 1st–90th percentiles. HTO counts were centre-log-ratio transformed and demultiplexed with HTODemux (positive.quantile = 0.99), retaining singlets. Because this experiment included 188 antibody-tagged oligonucleotides (ADT), the ADT profiles were further filtered to keep cells whose total ADT UMIs were within the 1st–99th percentile of the lane distribution. We found that batch B in the dataset had a dramatically lower distribution of UMIs than any of the other batches, so we removed those cells from further processing, then merged the three lane-specific Seurat objects. To validate guide sequences, we established a local BLAT server using the hg38 reference genome (hg38.2bit) with parameters -stepSize=5 for the server and -minScore=20 -minIdentity=0 for client queries, requiring 100% sequence similarity to unique genomic locations. The guides were then overlapped with the K562 candidate elements.

### Processing of CRISPRi-Perturb-seq data from Xie *et al*., 2019

We obtained count files from GEO accession GSE129837. For each batch we constructed a Seurat object and removed low-quality cells (unique-gene and total-UMI counts below the 10th percentile or above the 99th, and mitochondrial-read fraction >10%). Guides were filtered by alignment to hg38 using BLAT and overlap with the DNase-seq elements in K562 defined above. We validated guide sequences using the approach described above for STING-seq v2 from Morris *et al.*, 2023^19^, then overlapped with the K562 candidate elements. The batch-level objects were merged to yield an mRNA expression matrix, a gRNA count matrix, and a metadata file.

### Processing of CRISPRi-Perturb-seq data from Klann *et al.*, 2021

We obtained the mRNA count matrix and the corresponding gRNA UMI matrix from ENCODE experiment ENCFF904ZDX. In line with the original analysis, the gene matrix was loaded into Seurat. Cells with >20 % mitochondrial UMIs or <10,000 total transcript UMIs were discarded, after which a single Seurat object was created containing the mRNA counts assay and the gRNA counts assay. Genomic coordinates in hg19 for the guides provided by the authors were lifted to GRCh38 with easylift, retaining only guides with unambiguous positions. The lifted-over guides were then overlapped with the K562 candidate elements.

### Analyzing differential expression and computing statistical power

We performed differential expression analysis using the same SCEPTRE framework as described above. For each dataset, we tested every gene against all targeted elements within 1 Mb of the gene’s transcription start site (TSS), with distances calculated using GENCODE v29 gene annotations. Any positive control perturbations (self promoters) were excluded from the tested pairs. SCEPTRE was configured using the high MOI setting to account for cells containing multiple guides, and a union integration strategy to aggregate effects across multiple guides targeting the same element. We employed two-sided testing to detect both activating and repressing effects, and included dataset-specific batch information as covariates when available. The analysis workflow consisted of guide assignment using SCEPTRE’s mixture model approach, followed by quality control filtering using SCEPTRE’s default quality control parameters and discovery analysis using SCEPTRE’s statistical framework. Consistent with the analyses of the K562 and WTC11 DC-TAP-seq screens, we applied a 20% FDR cutoff to define significant distal element-gene pairs **(Extended Data Fig. 8)**. We used a simulation-based method to compute statistical power for each element-gene pair (see section below).

### Annotation and filtering of element-gene pairs for external datasets

We annotated element-gene pairs overlapping gene promoters and target gene bodies using the same approach described above for the K562 and WTC11 DC-TAP-seq datasets. However, for the uniformly processed external datasets, GENCODE v29 genome annotations were used to define promoters and gene bodies and no manual adjustments were made. Any element-gene pairs where the targeted element overlapped an annotated promoter or any portion of the target gene’s body were filtered out.

### Computing statistical power in DC-TAP-seq and Perturb-seq datasets using SCEPTRE

To ensure confidence in the nonsignificant hits, we developed an empirical simulation-based approach to estimate statistical power for each element-gene pair in a given Perturb-seq dataset. For DC-TAP-seq and other datasets, we ran this analysis on all element-gene pairs that passed SCEPTRE quality controls by simulating different effect sizes in gene expression (*e.g.*, 5%, 10%, 15%, 20%, 25%, and 50% decreases), then testing the simulated counts for differential expression. To simulate UMI counts, we used a negative binomial distribution for each gene with SCEPTRE-derived dispersion estimates and DESeq2 size factor normalized mean^64^. This approach was necessary because: (1) SCEPTRE does not directly calculate size factors, which are essential for modeling realistic cell-to-cell technical variation in simulated counts, (2) mean expression values must be mathematically consistent with the size factors used to generate technical variability, requiring coordinated calculation from DESeq2, and (3) using SCEPTRE’s dispersion estimates ensures our power analysis reflects the actual statistical framework used for differential expression testing.

Using the number of perturbed cells determined by SCEPTRE for each element-gene pair, a decrease in expression was simulated for the same number of randomly selected cells. UMI counts for perturbed and control cells were drawn from the negative binomial gene expression distributions. Unchanged distributions with mean expression levels across all cells were used to simulate counts from control cells and effects in perturbed cells were simulated by injecting a specified effect size (decrease in mean expression). To simulate variability in effect sizes for guides targeting the same enhancer, in each iteration a guide-guide variability value was randomly drawn from a normal distribution with mean 1 and standard deviation of 0.13 for each guide and added to the specified effect size. The standard deviation for this normal distribution was learned by analyzing guide-guide variability from FlowFISH experiments^16^ across the range of observed effect sizes of significant enhancer-gene interactions. Differential expression testing was performed on simulated counts from each iteration with SCEPTRE using the same parameters and covariates as in the full screen. Simulation and differential expression analysis of each effect size was performed 100 times for each element-gene pair.

Our SCEPTRE analysis procedure uses the Benjamini-Hochberg procedure to control the false discovery rate. This procedure depends on the distribution of *p-*values in the experiment. To account for multiple testing of simulated perturbations in the context of the p-value distribution of the given CRISPR screen, a nominal p-value cutoff corresponding to the FDR threshold used in the differential expression analysis of the observed perturbation data was applied. Power was then calculated as the percentage of pairs out of the 100 reps that were significant with the correct direction of effect. The comparison of power across different datasets in Figure 3 uses FDR of 20% across all datasets.

Implementation code is provided in the sceptre_power_analysis snakemake ruleset within the analysis pipeline detailed in **Code Availability**.

### Estimating number of detected and undetected positives

We estimated the number of undetected significant downregulated DE-G pairs with effect sizes in a given range utilizing the power SCEPTRE calculations described in the previous section. For the analysis shown in **Fig. 3c**, we used the set of random distal element-gene pairs for DC-TAP-seq and distal element-gene pairs for Gasperini *et al*.^14^ For both datasets, we used power calculations performed without positive controls and defined “detected positives” as DE-G pairs called as significant at the 20% FDR threshold. To calculate the “true” number of positives in an effect size bin, we first summed the power to detect effect sizes at the upper limit of the bin across all tested pairs (*e.g.*, for the 5–10% bin, we summed power to detect 10% effect sizes). We then calculated a “detected positive rate” as the number of detected positives divided by the sum of power across tested pairs, and estimated the total number of positives (detected and undetected) as the positive rate multiplied by the total number of tested pairs.

This approach yields an approximate decomposition of positives into “detected” and “not detected at current power” categories that can be compared across datasets. We note that this estimation rests on several simplifying assumptions and has important limitations: it ignores heterogeneity of true effect sizes within each bin, is affected by selection bias due to binning on noisy effect size estimates, and assumes all effect sizes in a bin equal the bin maximum. In addition, it treats all significant positives as true positives, which overlooks false positives and potential indirect effects. While more realistic modeling would require considerably more complex formulas, we view this approach as a useful first-order summary and note that it should be interpreted with these assumptions in mind.

### Computing the probability that DE-G pair is a direct versus indirect effect

The observed DE-G pairs in CRISPR perturbation experiments contain direct, *cis*-acting effects of element perturbations, indirect, *trans*-acting effects, and false discoveries. We assume that effects of element perturbations on genes on other chromosomes are either indirect effects or false discoveries (hereafter, “indirect effects” refers to both of these categories). To calculate the direct effects rate in the used CRISPR datasets, we first empirically estimated the indirect effects rates. We calculated the distance-independent rates of indirect effects for the K562 and WTC11 DC-TAP-seq experiments and other CRISPR datasets processed as described^29^ by systematically testing distal element perturbations for effects on genes on other chromosomes. For each dataset we applied the same differential expression testing approach used to identify DE-G pairs, but tested each distal element perturbation against 100 randomly selected genes on other chromosomes. Only genes in the *cis* discovery pairs (see Methods section: Creating discovery pairs for SCEPTRE input) were considered to ensure similar expression distributions of tested genes between *cis*- and *trans*-acting analyses. The dataset specific indirect effects rate was calculated as the proportion of significant *trans*-acting differential expression tests (**Extended Data Fig. 16a**) using the uncorrected p-value cutoff corresponding to the chosen FDR cutoff in each dataset (see Methods section: Differential expression analysis). This approach was used to calculate indirect effect rates for both positive and negative effects on expression.

To calculate the direct effects rate for each dataset, first we calculated the observed positive hit rate in the *cis* discovery analysis as a function of distance to TSS. The *cis* discovery pairs within 1Mb of the TSS were grouped into 50 kb bins, and for each bin the positive hit rate was calculated as the proportion of positive tests for both positive and negative effects on expression. To calculate the rate of direct effects, for each distance bin the distance independent indirect effects rate was subtracted from the observed positive hit rate.

We calculated the average direct rate per distance bin across all used datasets and fitted a power law function to model the direct effects rate as function of distance to TSS using the `lm` function in R (4.2.0). Only distance bins with a direct rate greater than zero were included in this analysis, which removed 1 bin (800-850kb from TSS) out of 20 for negative and none for positive effects on expression (**Extended Data Fig. 15a**). This model was then applied to predict the expected direct effects rate for all *cis* discovery pairs in used CRISPR datasets based on their distance to TSS (**Extended Data Fig. 15**).

The probability of each *cis* discovery pair being the result of a direct versus an indirect effect was calculated from the expected direct effects rate based on its distance to TSS and the datasets indirect effects rate:

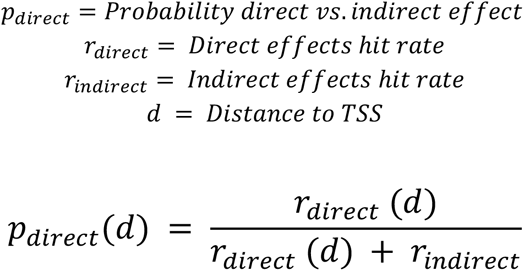

Under this model, when the two rates are equal, a significant hit is equally likely to arise from a direct or an indirect effect, and *p_direct_* is therefore 0.5. This approach was used to calculate probabilities of direct effects for both positive and negative effects on gene expression (**Extended Data Fig. 16d**). Calculated *trans*- and *cis*-rates per dataset, estimated direct effect rates per dataset and averaged across dataset, and fitted models for direct effect rates as function of distance to TSS are available on github: https://github.com/EngreitzLab/DC_TAP_Paper/tree/v2.0.3/direct_vs_indirect_effects/compute_direct_vs_indirect_effects/results.

To test the robustness of this analysis to the distance bin size used to compute the direct effects rate, we recalculated *p_direct_* using 25 kb and 100 kb bins and found that main conclusions remained unchanged.

To quantify the uncertainty of the probability of direct effects for each *cis* discovery pair, we used a non-parametric cluster bootstrap based on resampling perturbations separately in each dataset used to calculate direct effect probabilities. In each bootstrap replicate, we sampled an equal amount of distal element perturbations as in the original dataset with replacement. All *cis* and *trans*-acting tests of a sampled perturbation were included together, as many times as that perturbation was drawn. This was done to account for correlation between tests involving the same perturbations and to capture how sensitive the estimates are to which elements were targeted. Instead of repeating differential expression testing for each bootstrap replicate, we sampled differential expression results from the original *cis* and *trans*-acting analyses (see above).

For each bootstrap replicate, we recalculated the following, as described above:

1. the distance-independent indirect effects rate of each dataset.
2. the observed positive hit rate in each distance bin within 1 Mb of the TSS.
3. the direct effects rate per bin, *i.e.* the observed hit rate minus the indirect effects rate, set to zero where negative.

For each bootstrap replicate, we averaged the direct effect rates per distance bin across datasets and refitted the power law function to model the direct effects rate as function to distance to TSS using non-zero bins as described above. Each replicate’s power law model and dataset-specific indirect effects rate were then used to compute *p_direct_* for every *cis* discovery pair based on its distance to TSS using the formula defined above. We performed 1,000 bootstrap replicates calculating 1,000 estimates of *p_direct_* per DE-G pair. We report the median as the central estimate and define the 95% confidence interval as the 2.5th and 97.5th percentiles (percentile method). We calculated this separately for positive and negative effects on gene expression (**Extended Data Fig. 16d**).

### Defining element chromatin categories

To analyze the chromatin/epigenetic features of tested elements, we resized the elements targeted in the DC-TAP-seq experiments (generally 300 bp) to 500 bp (+/-250 bp from center) to make element sizes consistent with the set of genome-wide candidate elements defined by the ABC/ENCODE-rE2G pipeline, excluding promoters. In cases where resizing introduced overlaps between elements, overlapping tested elements were merged into a single tested element. We then determined whether resized elements overlapped H3K27ac, H3K27me3, or CTCF ChIP-seq peaks in their respective cell types. H3K27ac and H3K27me3 peaks were extended by 175 bp on each side due to the broad distribution of histone ChIP-seq signals. We also computed the H3K27ac RPM in each element expanded by 150 bp on each side. The epigenomic datasets used for this analysis are listed in **Supplementary Table 8.**

Next, we categorized elements according to the following ordered rules: If element overlaps H3K27ac peak:

- H3K27ac RPM in element >90% for cell type → High H3K27ac element
- Else → H3K27ac element

Else if element does not overlap H3K27ac peak:

- H3K27ac RPM in element >90% for cell type → H3K27ac high element
- H3K27ac RPM in element >50% for cell type → H3K27ac element
- Element overlaps CTCF peak → CTCF element
- Element overlaps H3K27me3 peak → H3K27me3 element
- Else → No H3K27ac element

To calculate the proportion of tested and significant element-gene pairs across a larger set of CRISPR perturbation datasets, we used datasets that we had previously curated, processed, and combined (available at https://github.com/EngreitzLab/CRISPR_comparison/blob/main/resources/crispr_data)^29^. We annotated tested and significant element-gene pairs as described above.

### Overlap with the broader ENCODE ChIP-seq catalog

To more fully characterize the chromatin context of tested elements, we intersected the same resized and merged elements from above with all additional histone modification, transcription factor, and other nuclear protein ChIP-seq experiments available for each cell type in ENCODE (hg38), beyond the H3K27ac, H3K27me3, and CTCF marks used for chromatin categorization (536 targets in K562, comprising 12 histone marks and 524 transcription factors and other nuclear proteins; 168 targets in WTC11, 28 histone marks and 140 transcription factors and other nuclear proteins). K562 elements were intersected with K562 datasets; WTC11 elements with matched iPSC datasets where available, otherwise the closest H1/H9 ESC experiments. Where multiple experiments targeted the same factor, we kept one representative, preferring conventional over Mint-ChIP-seq and better ENCODE audit status. Histone-modification peaks were extended by 175 bp on each side, as above; transcription-factor and CTCF peaks were used as called.

To test for enrichment, we compared the fraction of significant (positive) element–gene pairs overlapping each target against a background of well-powered tested elements using a two-sided Fisher’s exact test, reporting odds ratios with 95% confidence intervals and Benjamini–Hochberg FDR, computed separately for negative- and positive-effect positives. The ENCODE datasets are listed in https://github.com/EngreitzLab/DC_TAP_Paper/blob/v2.0.3/results/encode_catalog_overlap/metadata/encode_selected_files.csv; overlap and enrichment results are in, https://github.com/EngreitzLab/DC_TAP_Paper/blob/v2.0.3/results/encode_catalog_overlap/tables/enrichment_by_assay.csv, and https://github.com/EngreitzLab/DC_TAP_Paper/blob/v2.0.3/results/encode_catalog_overlap/tables/element_manifest.csv.

### Enrichment and recall of fine-mapped eQTLs

To test for eQTL enrichment in the labeled chromatin categories, we overlapped genome-wide promoter elements and the categorized distal elements with fine-mapped eQTLs in the same cell types (eQTLs from LCLs compared with elements from GM12878), using eQTL Catalogue v7 release (https://www.ebi.ac.uk/eqtl/). We defined promoters as previously described^28^ (https://github.com/broadinstitute/ABC-Enhancer-Gene-Prediction/blob/main/reference/hg38/CollapsedGeneBounds.hg38.TSS500bp.bed). The code and table describing the collation of eQTL variants is available here: https://github.com/anderssonlab/scE2G_analysis/blob/v1.0/3.Benchmarking/eQTL/eQTL_Catalogue_v7. To control for differences in statistical power due to expression level of eGenes, we selected a set of 1,041 non-housekeeping genes with expression in TPM matching the distribution of the 1,041 housekeeping genes by deciles. This resulted in a set of 100,814 variant-eGene pairs in the non- housekeeping set and 100,463 variant-eGene pairs in the housekeeping set used for calculating eQTL enrichment and recall.

We calculated enrichment and recall of fine-mapped variants with posterior inclusion probability (PIP) values in four bins: (0.01, 0.1], (0.1, 0.5], (0.5, 0.9], (0.9, 1]. For each element category and PIP bin, we calculated enrichment and recall as follows:

Enrichment(PIP bin, element category) = (fraction of variants in PIP bin overlapping elements in category) / (fraction of variants with PIP < 0.01 overlapping elements in category) Recall(PIP bin, element category) = fraction of variants in PIP bin overlapping elements in category

### Annotating DE-G pairs with enhancer-gene model scores

To annotate DE-G pairs from the K562 and WTC11 datasets with scores from the ENCODE-rE2G, ABC, and AlphaGenome predictive models, we resized the elements targeted in the DC-TAP-seq experiments (generally 300 bp) to 500 bp and merged overlapping regions as described above. If at least one of the merged elements had a significant effect on a given gene, the merged element was annotated as having a significant effect on the same gene. The resized element-gene pairs were annotated with ENCODE- rE2G scores using the CRISPR benchmarking pipeline as previously described^29^: https://github.com/EngreitzLab/CRISPR_comparison. The intermediate output file `expt_pred_merged_annot.txt.gz` was used in subsequent analyses to obtain predictor scores for all element gene-pairs.

### Data visualization

Box plots: Box plots are defined as follows: the middle line or point corresponds to the median; lower and upper hinges correspond to first and third quartiles; the upper whisker extends from the hinge to the largest value no further than 1.5 × IQR from the hinge (where IQR is the interquartile range, or distance between the first and third quartiles); and the lower whisker extends from the hinge to the smallest value, at most 1.5 × IQR of the hinge. Data beyond the end of the whiskers are outlying points and are plotted individually, unless box plots are shown on top of all data points or data distribution.

## Data Availability

- Random DC-TAP-seq screens are available on the IGVF Data Portal (https://data.igvf.org/): IGVFDS7288SJVF (K562) and IGVFDS3911MOCN (WTC11).
- Pilot DC-TAP-seq, validation DC-TAP-seq and CTCF ChIP-seq experiments are available at the NCBI Gene Expression Omnibus (GSE303901)
- Epigenomic datasets used to annotate WTC11 and K562 elements are available on the ENCODE Portal (www.encodeproject.org). File accessions are listed in **Supplementary Table 8.**
- Gene annotations (gencode.v32lift37.annotation.gtf) were obtained from GENCODE (https://ftp.ebi.ac.uk/pub/databases/gencode/Gencode_human/release_32/GRCh37_mapping/).
- The list of housekeeping genes was obtained from the “is_ubiquitous_uniform” column in the following gene annotations: https://github.com/EngreitzLab/ENCODE_rE2G/blob/main/resources/external_features/gene_pr omoter_class_RefSeqCurated.170308.bed.CollapsedGeneBounds.hg38.TSS500bp.tsv

## Code Availability

- DC-TAP-seq analysis pipeline: https://github.com/EngreitzLab/DC_TAP_Paper/tree/v2.0.3 (DOI: 10.5281/zenodo.24060844)
- Design and selection of gRNAs: https://github.com/broadinstitute/CRISPRiTilingDesign
- Design of DC-TAP-seq primers: https://www.bioconductor.org/packages/release/bioc/html/TAPseq.html
- ABC pipeline: https://github.com/broadinstitute/ABC-Enhancer-Gene-Prediction/
- ENCODE-rE2G pipeline: https://github.com/EngreitzLab/ENCODE_rE2G/tree/v1.0.0
- Comparison of CRISPR data to predictive models: https://github.com/EngreitzLab/CRISPR_comparison/tree/main
- Computing probabilities of direct and indirect effects: https://github.com/EngreitzLab/DC_TAP_Paper/tree/v2.0.3/direct_vs_indirect_effects

## Resource availability

- The sgOpti-CS vector used for gRNA expression is available on Addgene (https://www.addgene.org/256351/)

## Acknowledgements

J.M.E. acknowledges support from the Novo Nordisk Foundation Center for Genomic Mechanisms of Disease (NNF21SA0072102); the NHGRI Impact of Genomic Variation on Consortium (UM1HG011972); NHLBI R01HL159176; the NHGRI Genomic Innovator Award (R35HG011324); the Applebaum Foundation; Gordon and Betty Moore; and the BASE Research Initiative at the Lucile Packard Children’s Hospital at Stanford University. M.U.S. acknowledges the support of an NSF Graduate Research Fellowship (DGE-1656518) and a graduate fellowship award from Knight-Hennessy Scholars at Stanford University. A.R.G. and L.M.S. acknowledge the support of R01HG011664 and the NHGRI Impact of Genomic Variation on Function Consortium (UM1HG011972). D.A. acknowledges support from an AHA Postdoctoral Fellowship (821920 and 23POSTCHF1019753) and the Stanford Maternal and Child Health Research Institute (MCHRI) Postdoctoral fellowship. E.K. acknowledges support from NSF DMS- 2113072 and NSF DMS-2310654. We thank Stanford University and the Stanford Research Computing Center for providing computational resources and support as part of the Sherlock High-Performance Compute Cluster, and the Stanford Genomics Facility for DNA sequencing services.

We thank members of the Engreitz Lab and the Novo Nordisk Foundation Center for Genomic Mechanisms of Disease for feedback on the DC-TAP-seq method and results. We thank Robin Andersson, Wei-Lin Qiu, Gavin Schnitzler, Elisa Donnard, Elizabeth Roberts,and X. Rosa Ma for technical advice and feedback on the manuscript.

## Author Contributions

J.R., E.J., and J.M.E. developed the DC-TAP-seq method.

J.R., E.J., D.A., C.J.M., A.R.G., G.M., V.S., L.M.S., and J.M.E. designed and conducted pilot DC-TAP-seq experiments.

J.R., E.J., D.A., C.J.M., A.B., M.M., and H.K. analyzed pilot DC-TAP-seq experiments.

J.R., E.J., D.A., C.J.M., A.R.G., and J.M.E. designed the random DC-TAP-seq experiments.

D.A. and C.J.M. conducted the random DC-TAP-seq experiment in K562 cells.

J.R., G.M., and J.H. conducted the random DC-TAP-seq experiment in WTC11 cells.

A.R.G, J.G., and J.M.E. developed the statistical power analysis approach and pipeline.

E.J., J.G., A.B., and M.M. conducted quality control, power analysis, and differential expression analysis for random DC-TAP-seq experiments.

J.G. and A.R.G. uniformly processed and analyzed data from previous CRISPR Perturb-seq datasets.

J.R., M.U.S., and J.G. analyzed DC-TAP-seq with respect to chromatin state.

M.M. and M.U.S. analyzed eQTL data.

M.U.S. analyzed DC-TAP-seq data with respect to housekeeping genes.

M.U.S., J.G., and A.R.G. assessed predictions of the ENCODE-rE2G model.

A.R.G. and J.M.E. designed and conducted analysis to distinguish direct and indirect effects.

T.B.,E.K., and E.M. contributed to SCEPTRE analysis.

E.M. contributed to data organization and reporting.

E.M., E.G., G.M., F.J.N., J.M.E, and J.H. designed the validation experiments

G.M., C.B.E., and R.I. conducted the validation experiments

E.M. analyzed the validation experiments

J.R., M.U.S., E.J., D.A., J.G., A.R.G., J.M.E., E.G. and E.M. wrote the manuscript with input from all authors.

J.M.E. provided overall supervision and obtained funding for the project.

## Supplementary Information

### Extended Data Figures

**Extended Data Fig. 1.**
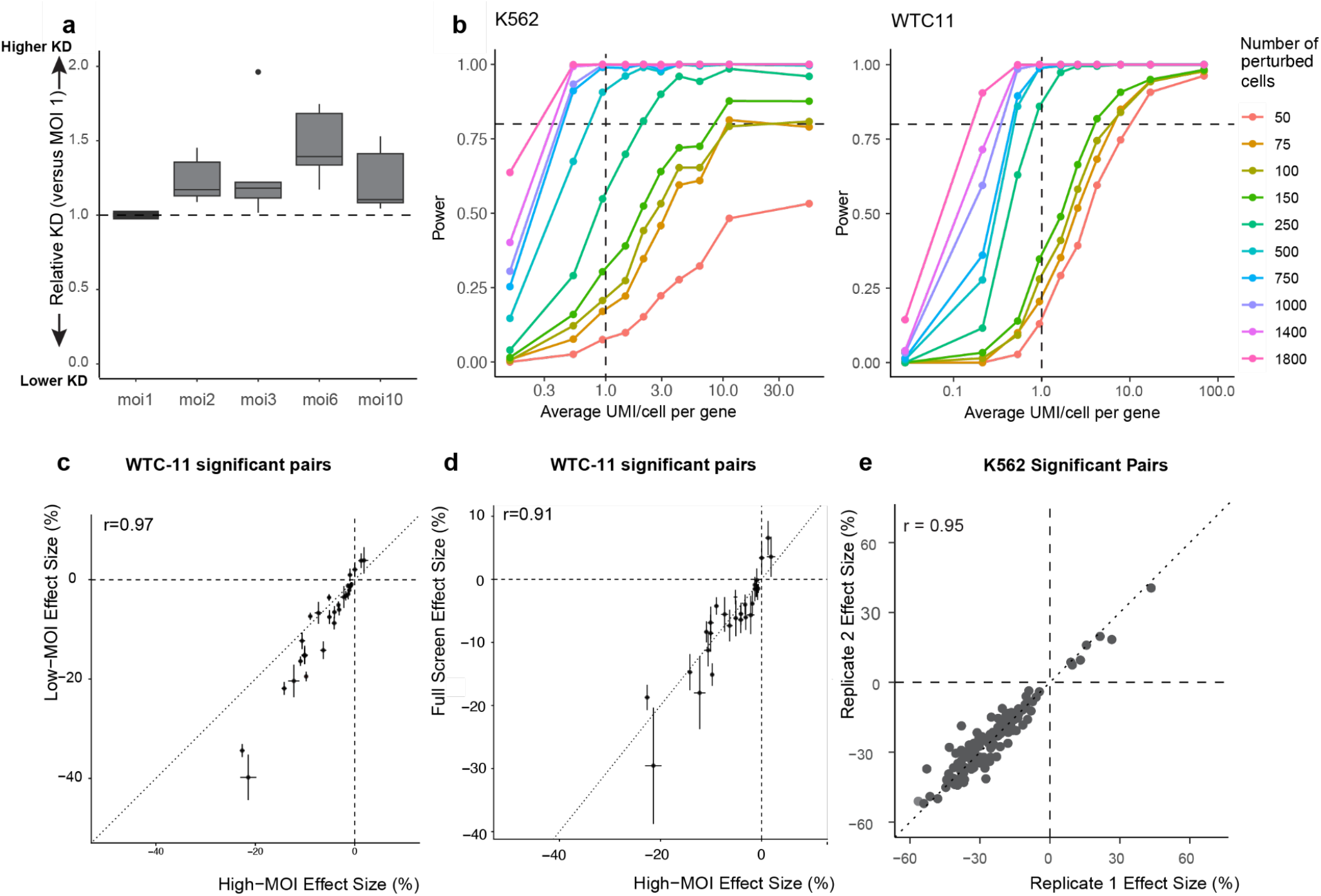
Effect of MOI on knockdown efficiency, and power analysis. **a.** Relative knockdown (KD) efficiency of guides for the promoters of *BAG3*, *NANOG*, *ROCK1*, *SOX2*, *YBX1* across different multiplicities of infection (MOI) in WTC11, normalized to effects of the respective guides at MOI 1. **b.** Power simulation curves for K562 (left) and WTC11 (right) estimating the number of cells per target required to achieve ≥80% power (horizontal dashed line) to detect a 25% effect size at an average mRNA capture efficiency of 1 UMI/cell/gene (vertical dashed line). **c.** Comparison of effect sizes between low MOI (multiplicity of infection)(MOI = 1.1 and 1.5 and high MOI (MOI = 12.6 and 14.4) DC-TAP-seq validation screens in WTC11. Effect size is represented as percent change in gene expression relative to control guides, with –100% indicating maximal knockdown. Error bars: 95% confidence intervals (c.i.) (n=2 biological replicates). **d.** Comparison of effect sizes between the large scale DC-TAP screen (MOI 5.07) and high MOI DC-TAP- seq validation screen (MOI = 12.6 and 14.4) in WTC11. Effect size is represented as percent change in gene expression relative to control guides, with –100% indicating maximal knockdown. Error bars: 95% confidence intervals (c.i.) (n=2 biological replicates) **e.** Reproducibility of effect sizes across technical replicates (random grouping of 10x lanes) in K562 for significant element-gene pairs (points).

**Extended Data Fig. 2.**
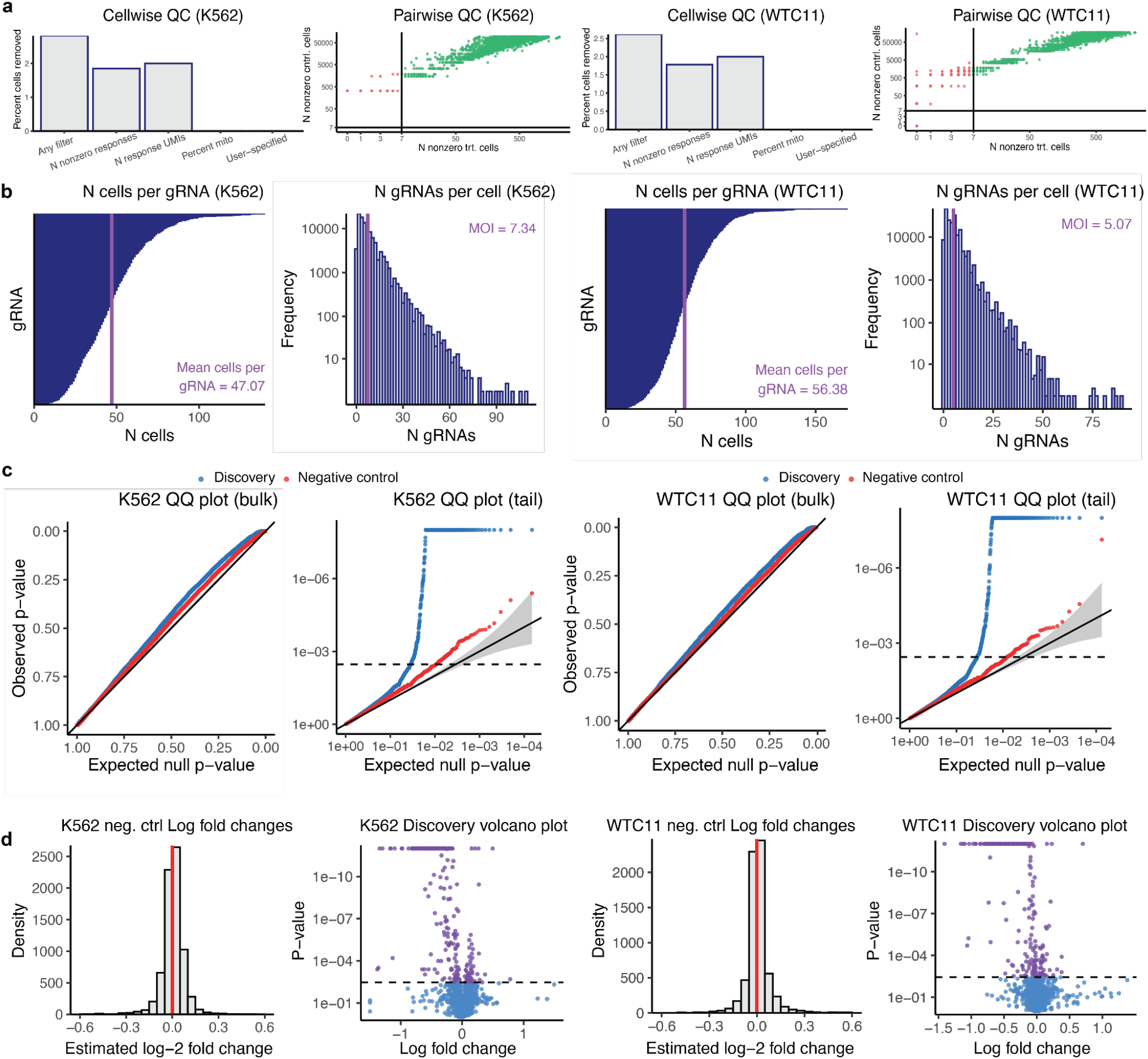
SCEPTRE analysis from K562 and WTC11 DC-TAP-seq experiments. **a.** SCEPTRE results of cellwise and pairwise QC plots for K562 (left) and WTC11 (right). Cellwise QC plot, horizontal axis denotes QC filters. Pairwise QC plot, each point corresponds to a target-response pair. **b.** Barplot of number of cells per gRNA and histogram of number of gRNAs per cell for K562 (left panels) and WTC11 (right panels). **c.** QQ plot of the p-values plotted for negative controls (red) and discovery pairs (blue) on an untransformed scale (bulk) and on a negative log10 transformed scale (tail) for K562 (left panels) and WTC11 (right panels). **d.** Histogram of estimated log_2_ fold changes for negative controls and volcano plot of the p-values and log_2_ fold changes of discovery pairs for K562 (left) and WTC11 (right). For K562, the number of false discoveries (at alpha 0.1) = 3, with a mean log_2_ fold change of 0.0026; number of significant discovery pairs called (at alpha 0.1) = 259 of 7493. For WTC11, the number of false discoveries (at alpha 0.1) = 2, with a mean log_2_ fold change of 0.0096; number of significant discovery pairs called (at alpha 0.1) = 237 of 6574. Alpha 0.1 refers to the parameter utilized by SCEPTRE to fit the null distribution. The number of SCEPTRE reported significant pairs include all discoveries from all tests performed (including promoters, positive controls and elements in gene bodies).

**Extended Data Fig. 3.**
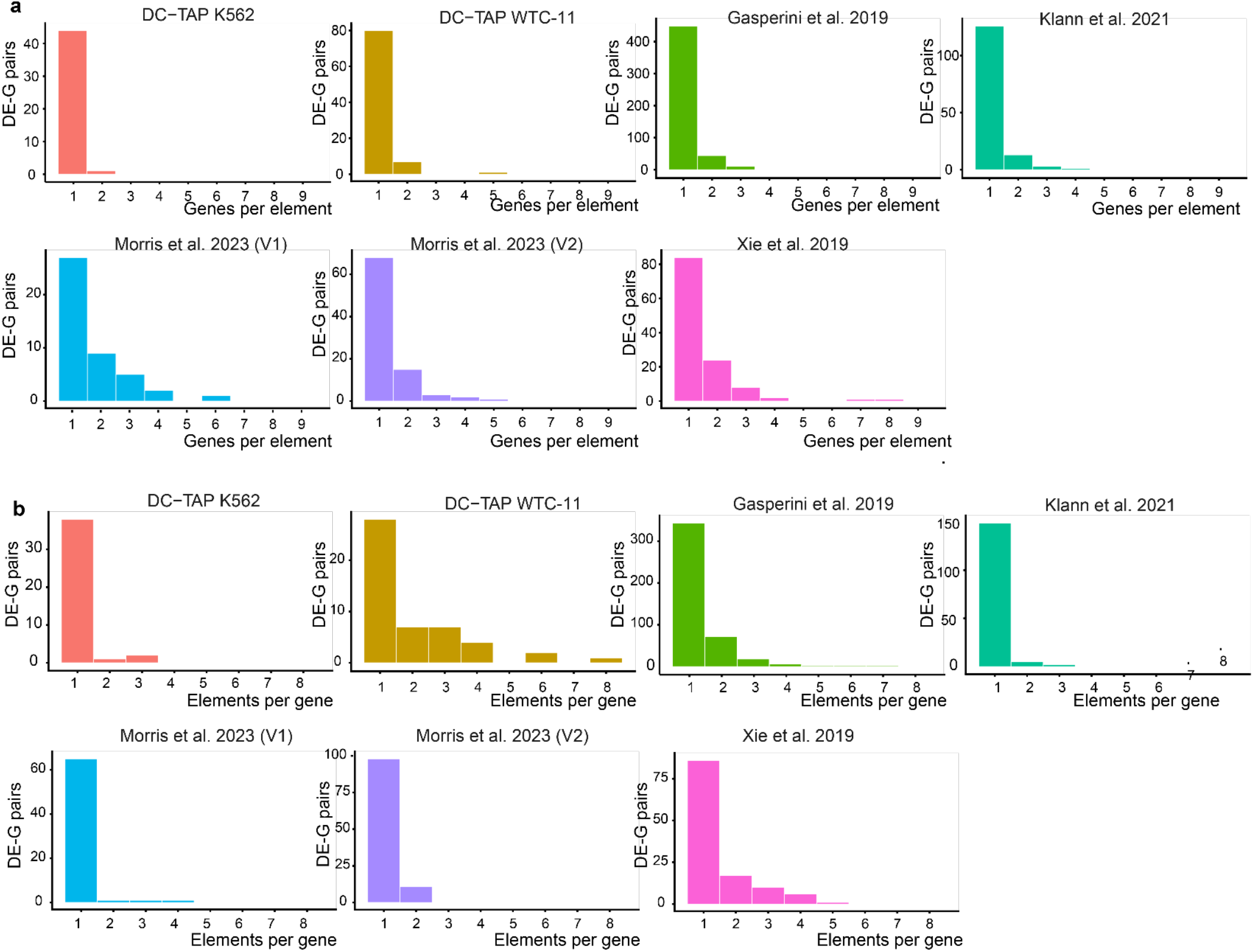
Summary statistics for element-gene connections. **a.** Summary statistics of genes per element for statistically significant (FDP 20%) DE-G pairs across these datasets and published studies. **b.** Summary statistics of elements per gene for statistically significant (FDP 20%) DE-G pairs across these datasets and published studies.

**Extended Data Fig. 4.**
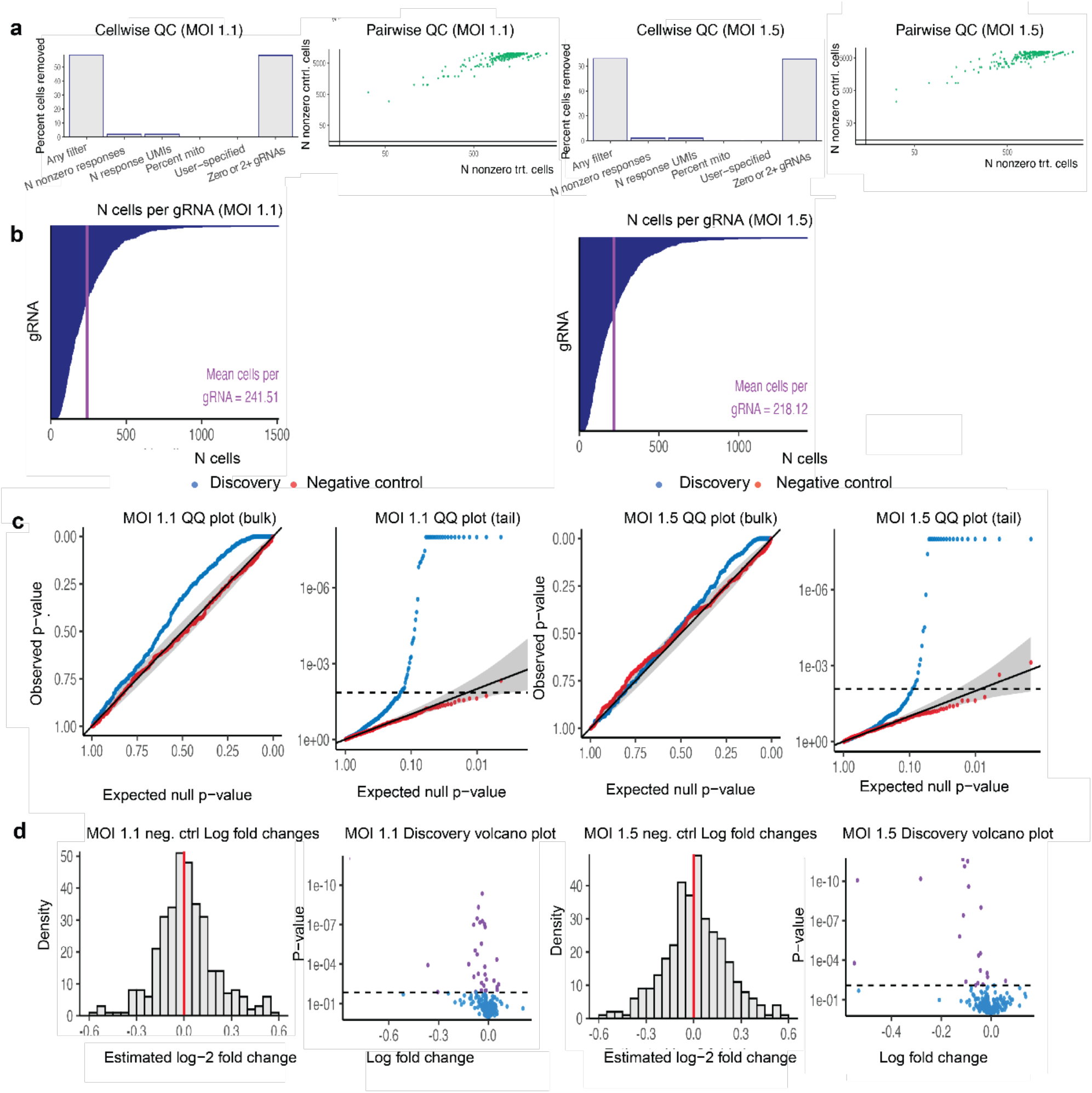
SCEPTRE analysis from WTC11 validation screens at low MOI. **a.** SCEPTRE results of cellwise and pairwise QC plots for WTC11 MOI 1.1 (left) and 1.5 (right). Cellwise QC plot, horizontal axis denotes QC filters. Pairwise QC plot, each point corresponds to a target- response pair. **b.** Barplot of number of cells per gRNA or WTC11 MOI 1.1 (left) and 1.5 (right). **c.** QQ plot of the p-values plotted for negative controls (red) and discovery pairs (blue) on an untransformed scale (bulk) and on a negative log10 transformed scale (tail) for or WTC 11 MOI 1.1 (left) and 1.5 (right). **d.** Histogram of estimated log_2_ fold changes for negative controls and volcano plot of the p-values and log_2_ fold changes of discovery pairs for or WTC11 MOI 1.1 (left) and 1.5 (right). For MOI 1.1, the number of false discoveries (at alpha 0.1) = 0, with a mean log_2_ fold change of 0.024; number of significant discovery pairs called (at alpha 0.1) = 51 out of 345. For MOI 1.5, the number of false discoveries (at alpha 0.1) = 0, with a mean log_2_ fold change of 0.014; number of significant discovery pairs called (at alpha 0.1) = 32 out of 345. Alpha 0.1 refers to the parameter utilized by SCEPTRE to fit the null

**Extended Data Fig. 5.**
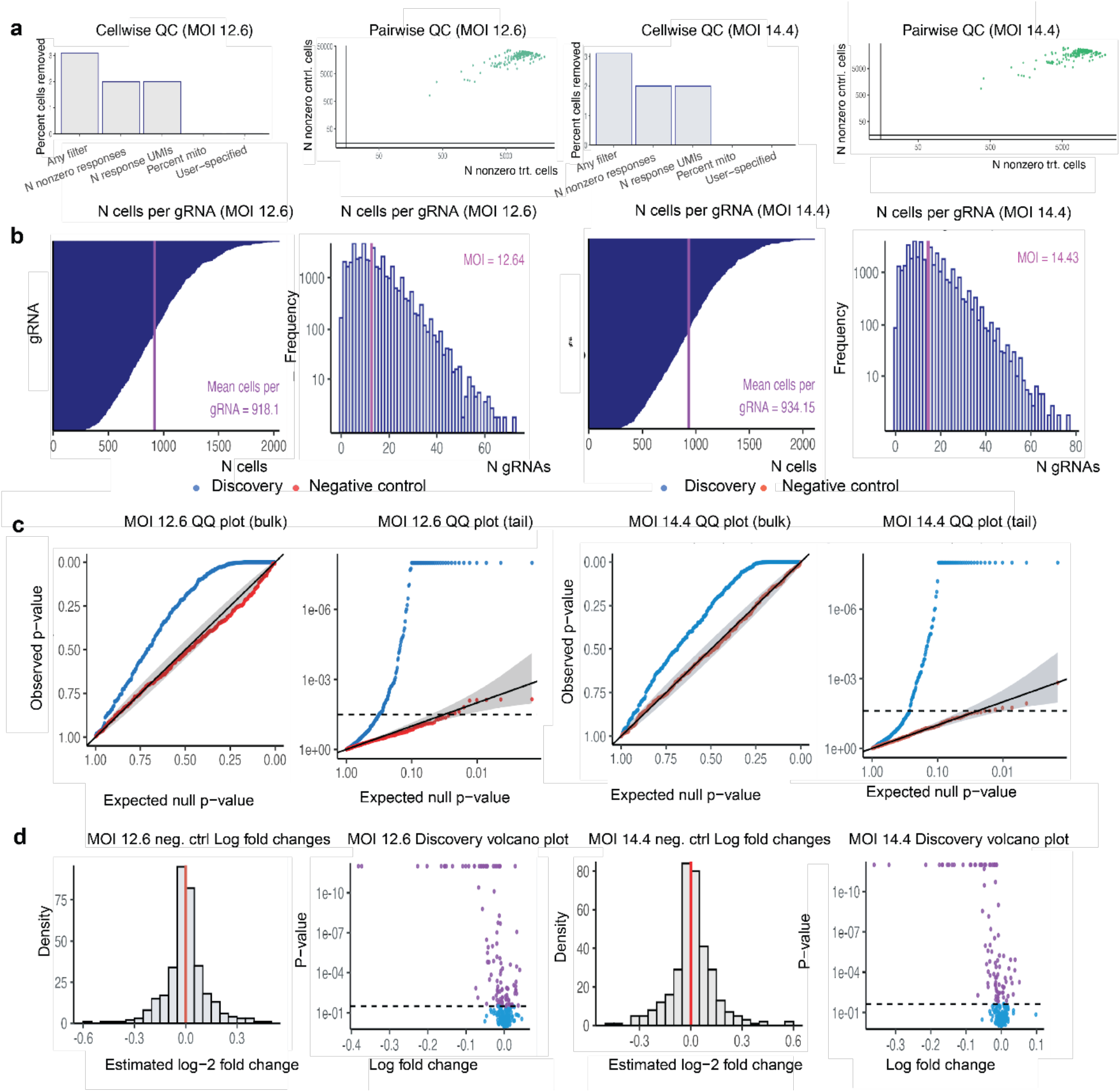
SCEPTRE analysis from WTC11 validation screens at high MOI. **a.** SCEPTRE results of cellwise and pairwise QC plots for WTC 11 MOI 12.6 (left) and 14.4 (right). Cellwise QC plot, horizontal axis denotes QC filters. Pairwise QC plot, each point corresponds to a target-response pair. **b.** Barplot of number of cells per gRNA or WTC 11 MOI 12.6 (left) and 14.4 (right). **c.** QQ plot of the p-values plotted for negative controls (red) and discovery pairs (blue) on an untransformed scale (bulk) and on a negative log10 transformed scale (tail) for or WTC 11 MOI 12.6 (left) and 14.4 (right). **d.** Histogram of estimated log_2_ fold changes for negative controls and volcano plot of the p-values and log_2_ fold changes of discovery pairs for or WTC 11 12.6 (left) and 14.4 (right). For MOI 12.6, the number of false discoveries (at alpha 0.1) = 0, with a mean log_2_ fold change of 0.0029; number of significant discovery pairs called (at alpha 0.1) = 111 out of 345. For MOI 14.4, the number of false discoveries (at alpha 0.1) = 0, with a mean log_2_ fold change of 0.0095; number of significant discovery pairs called (at alpha 0.1) = 93 out of 345. Alpha 0.1 refers to the parameter utilized by SCEPTRE to fit the null

**Extended Data Fig. 6.**
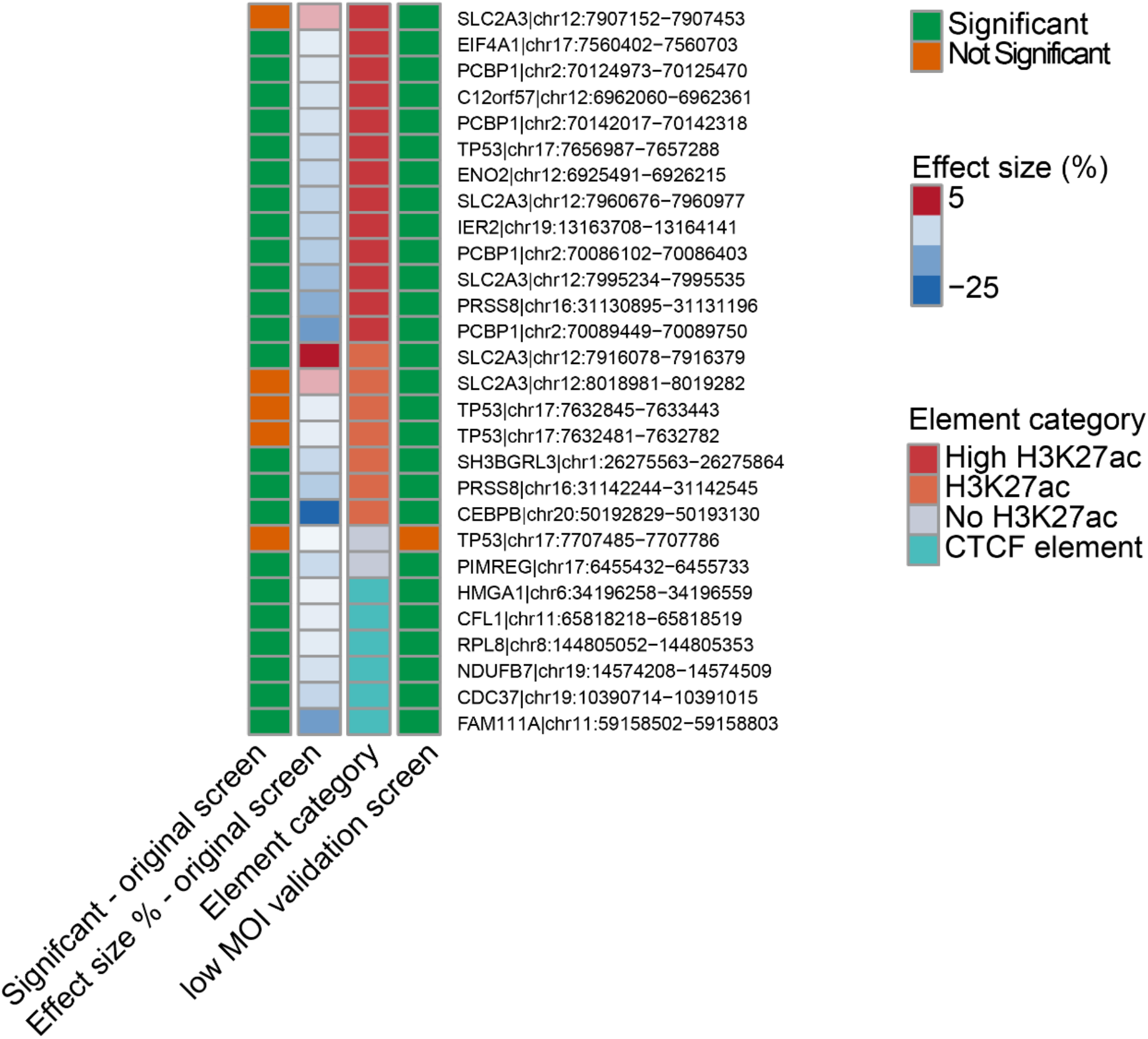
Summary of validated DE-G pairs in the original screen and the low MOI validation screen. Heatmap shows comparison of DE-G pairs tested in both the original large scale screen and low MOI validation screen in WTC11. The “23 of 23” results were calculated from a single SCEPTRE analysis of the two low-MOI transductions pooled. Transduction and 10x channel were included as covariates in the SCEPTRE model, together with per-cell technical covariates.

**Extended Data Fig. 7.**
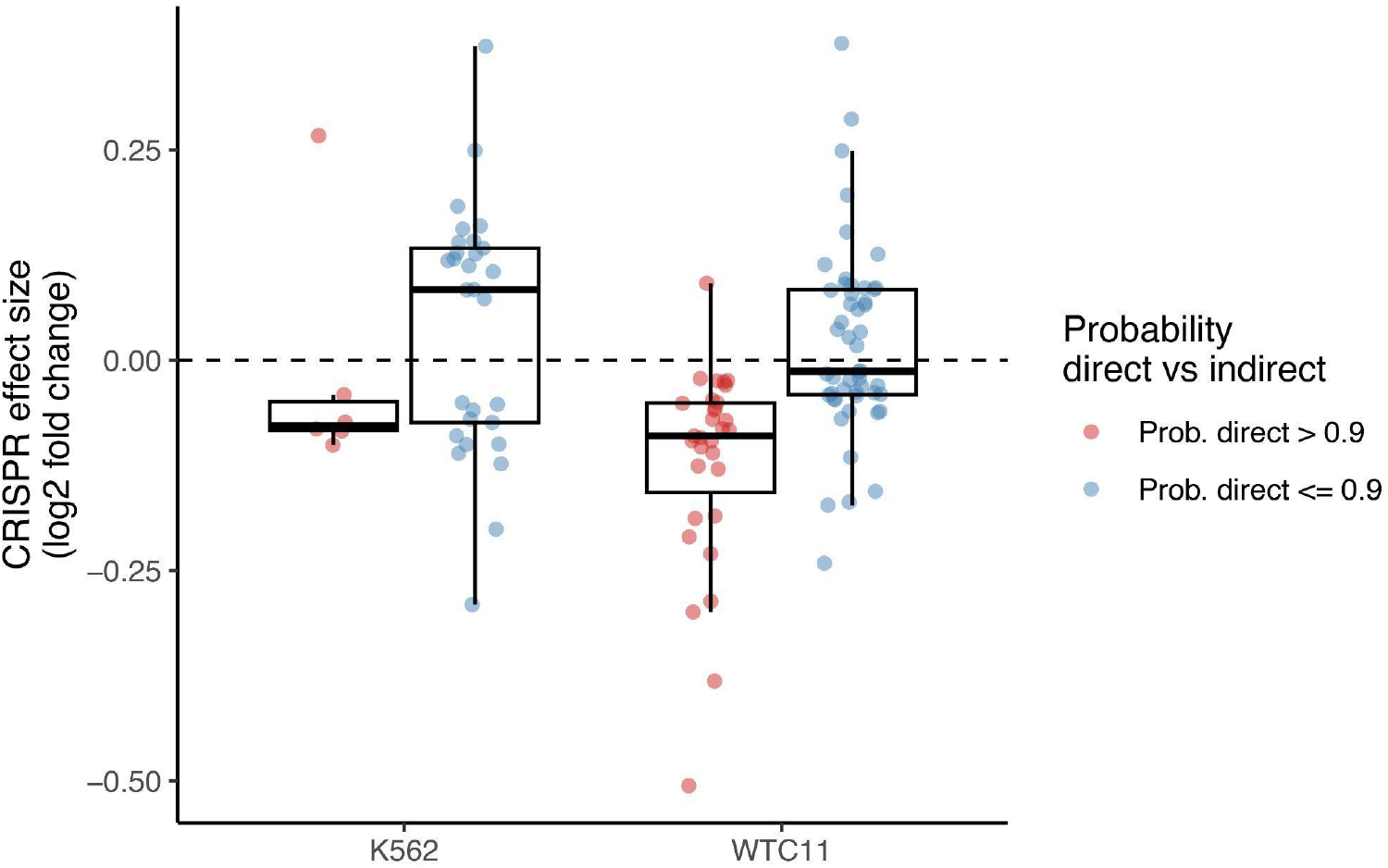
CRISPR effect size of likely direct vs indirect enhancer effects. CRISPR effect sizes (log_2_ fold change) for significant local DE-G pairs versus probability of being direct enhancer (negative) effects. DE-G pairs with likely direct effects were defined by an estimated probability of being a direct negative effect based on distance to TSS (see **Methods**) greater than 0.9.

**Extended Data Fig. 8.**
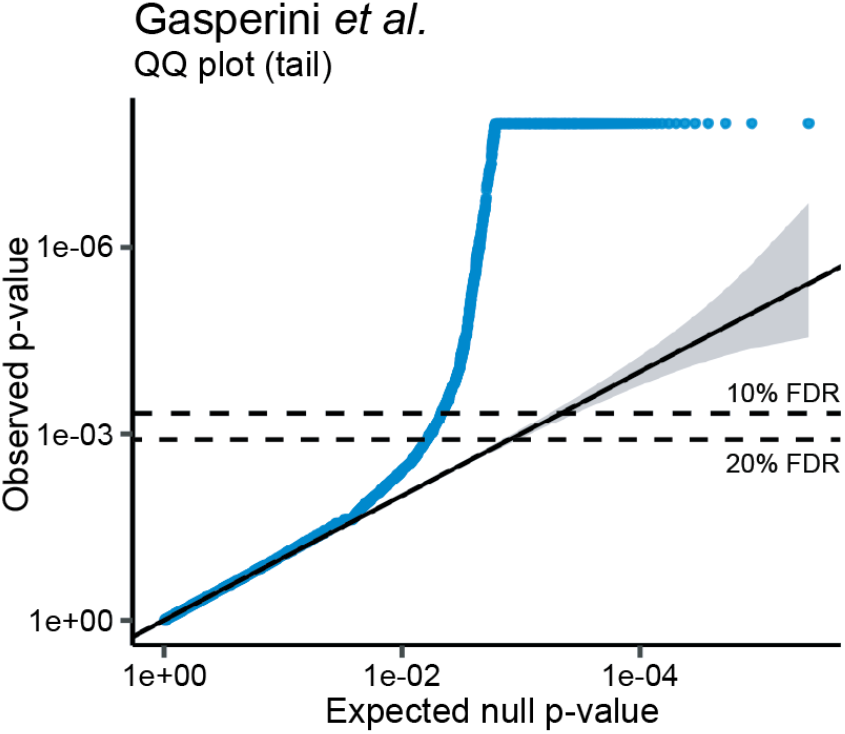
Observed versus expected p-values from Gasperini *et al.*^14^. QQ plot of the p-values plotted for discovery pairs (blue) on a negative log_10_ transformed scale (tail) for the Gasperini *et al.* dataset. Dashed lines indicate observed p-values corresponding to the 10% FDR and 20% FDR cutoffs (upon removal of positive controls).

**Extended Data Fig. 9.**
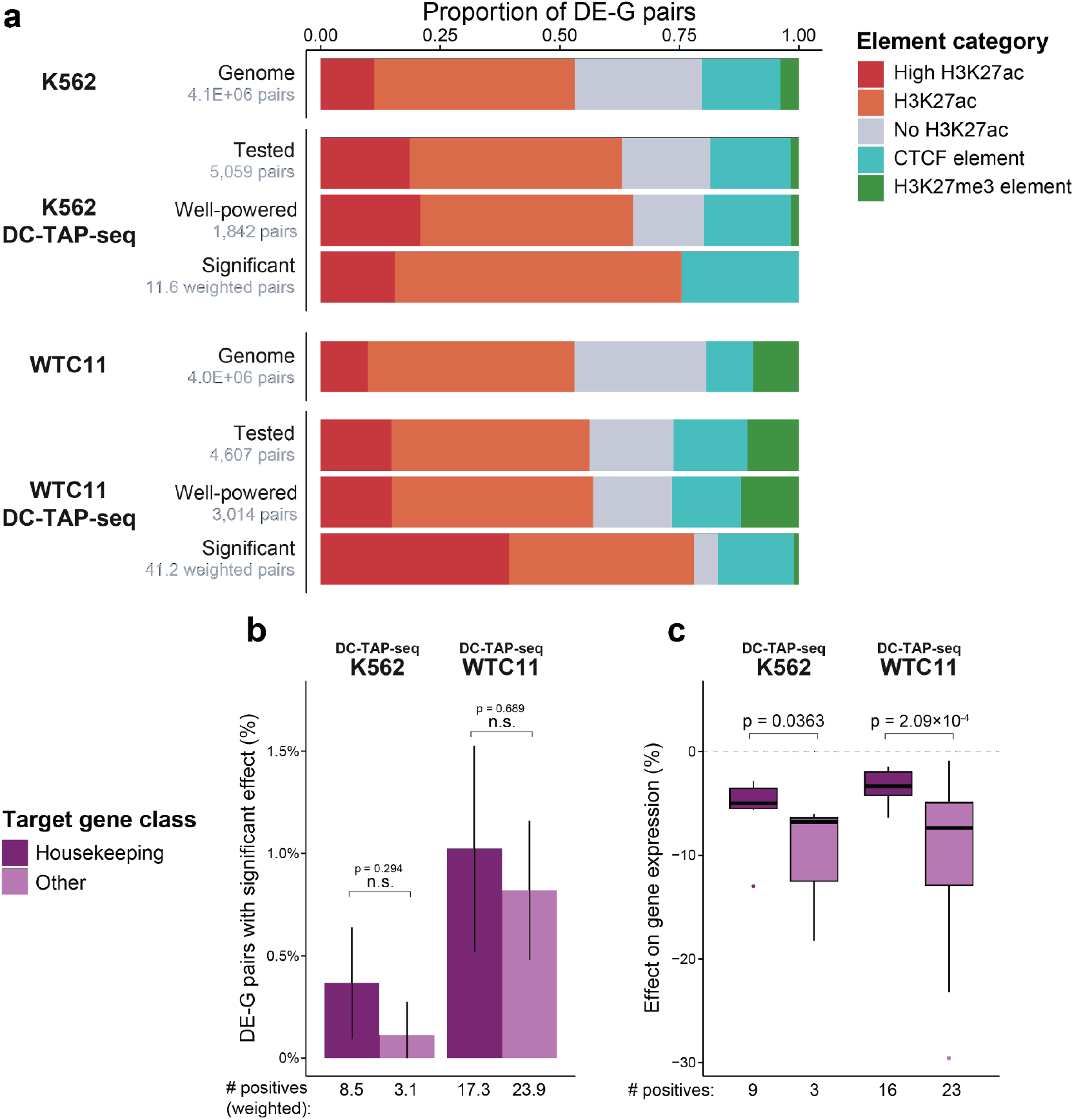
Element chromatin category and gene class of DC-TAP-seq DE-G pairs by cell type. **a.** Proportion of DE-G pairs for which the element is classified into different chromatin categories for all genome-wide DE-G pairs, tested DE-G pairs, well-powered DE-G pairs (tested with power to detect a 15% effect size in ≥80% of pairs), and DE-G pairs with negative effect sizes found as significant (weighted by probability of direct effect) for the K562 and WTC11 DC-TAP-seq datasets. **b.** Fraction of tested DE-G pairs from DC-TAP-seq for each cell type that were found to have a significant negative effect (weighted by probability of direct effect), stratified by target gene class. Error bars indicate 95% confidence intervals around the observed proportion, with the standard error estimated from the Agresti–Coull adjusted proportion and lower bounds are truncated at 0. Numbers below indicate the number of significant pairs with the target gene in the indicated class, weighted by probability of direct effect. P-values report significance from two-sample chi-squared tests with Yates’ continuity correction, adjusted with Benjamini-Hochberg correction. **c.** Effect sizes for significant DE-G pairs from DC-TAP-seq for each cell type with negative effect sizes (probability of direct effect > 50%) stratified by target gene class. P-values report significance from two- sided Wilcoxon rank-sum tests comparing housekeeping and other target genes within each dataset.

**Extended Data Fig. 10.**
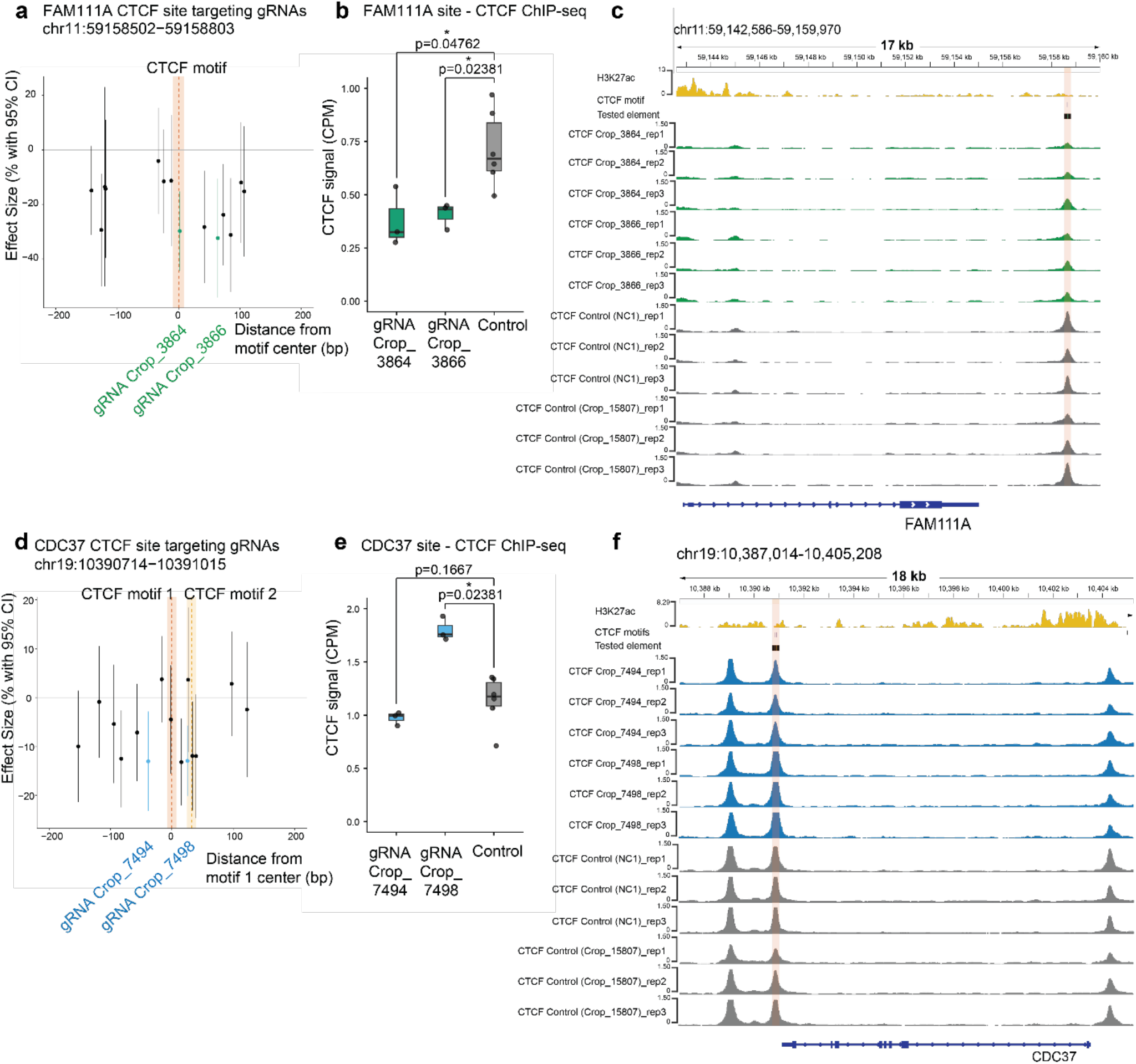
CRISPRi targeting of CTCF bound elements can reduce CTCF binding. Evaluation of CTCF binding disruption at two genomic sites in WTC11: chr11:59158502−59158803, target gene FAM111A **(a-c)** and chr19:10390714−10391015, target gene CDC37 (**d-f**). **a.** Positional mapping of gRNA effect size in the large-scale high MOI WTC11 screen relative to the target CTCF motif(s) in the FAM111A locus (indicated by the vertical orange shaded region). Black and colored dots represent individual gRNA effect sizes, with error bars denoting 95% C.I. Target guides chosen for validation are highlighted in green. **b.** Quantification of CTCF ChIP-seq following arrayed infection with either targeting gRNAs or controls at the FAM111A locus. Individual data points are shown as black dots (3 biological replicates per condition). p-values: Wilcoxon two-sided test. **c.** Tracks represent the FAM111A locus region: from top to bottom, H3K27ac ChIP-seq signal (ENCFF306ZMK), fold-change; CTCF motif location (http://hgdownload.soe.ucsc.edu/gbdb/hg38/jaspar/JASPAR2024.bb motif accession: MA0139); target element; CTCF ChIP seq (this study); gene track. **d.** As in **(a)**, now showing the CDC37 locus, target guides chosen for validation are highlighted in blue. **e.** As in **(b)**, now for the CDC37 locus. **f.** As in **(c)**, now for the CDC37 locus.

**Extended Data Fig. 11.**
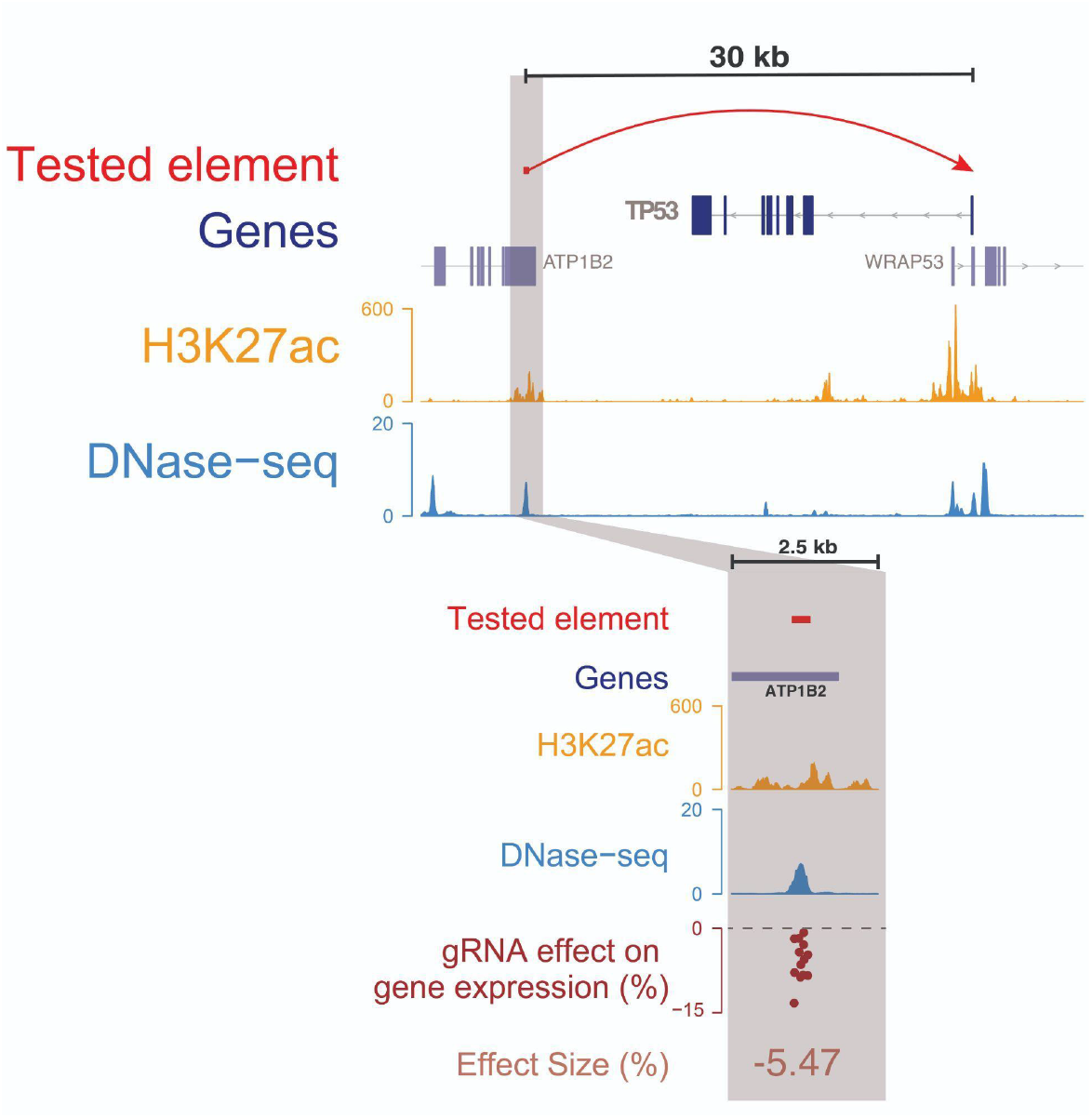
Regulation of a housekeeping gene by distal elements. Example locus showing a significant element-gene pair identified in WTC11 for the housekeeping gene *TP53.* Tracks show link between *TP53* and the tested element chr17:7656987-7657288, DNase-seq signal, H3K27ac ChIP-seq signal. The zoomed-in panel in the bottom shows the tested element, H3K27ac, DNase- seq, and the effect sizes of gRNAs on *TP53* expression.

**Extended Data Fig. 12.**
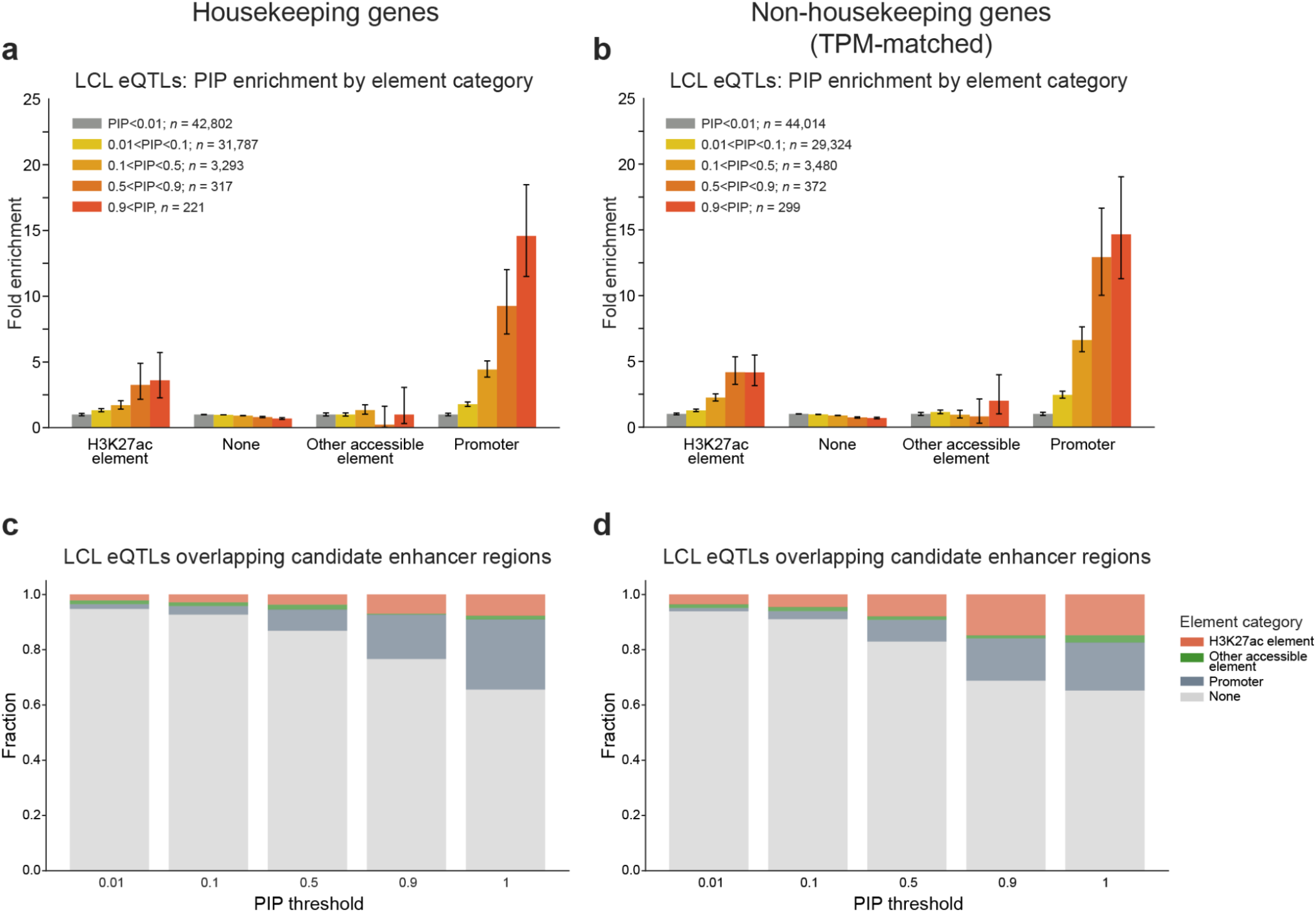
Enrichment and recall of fine-mapped eQTLs in elements with different chromatin states. **a.** Enrichment of fine-mapped eQTLs linked to housekeeping genes in LCLs that overlap element sets defined in Fig 5. Here, “H3K27ac element” refers to the chromatin categories “High H3K27ac” and “H3K27ac.” “Other accessible element” refers to the categories “H3K27me3 element,” “CTCF element,” and “No H3K27ac. “None” indicates non-accessible regions. **b.** Same as **a**, but for fine-mapped eQTLs linked to a set of non-housekeeping genes with TPMs matched to the housekeeping genes by deciles. **c.** Total fraction of LCL eQTLs linked to housekeeping genes that overlap elements in each category. **d.** Same as **c**, but for LCL eQTLs linked to TPM-matched non-housekeeping genes.

**Extended Data Fig. 13.**
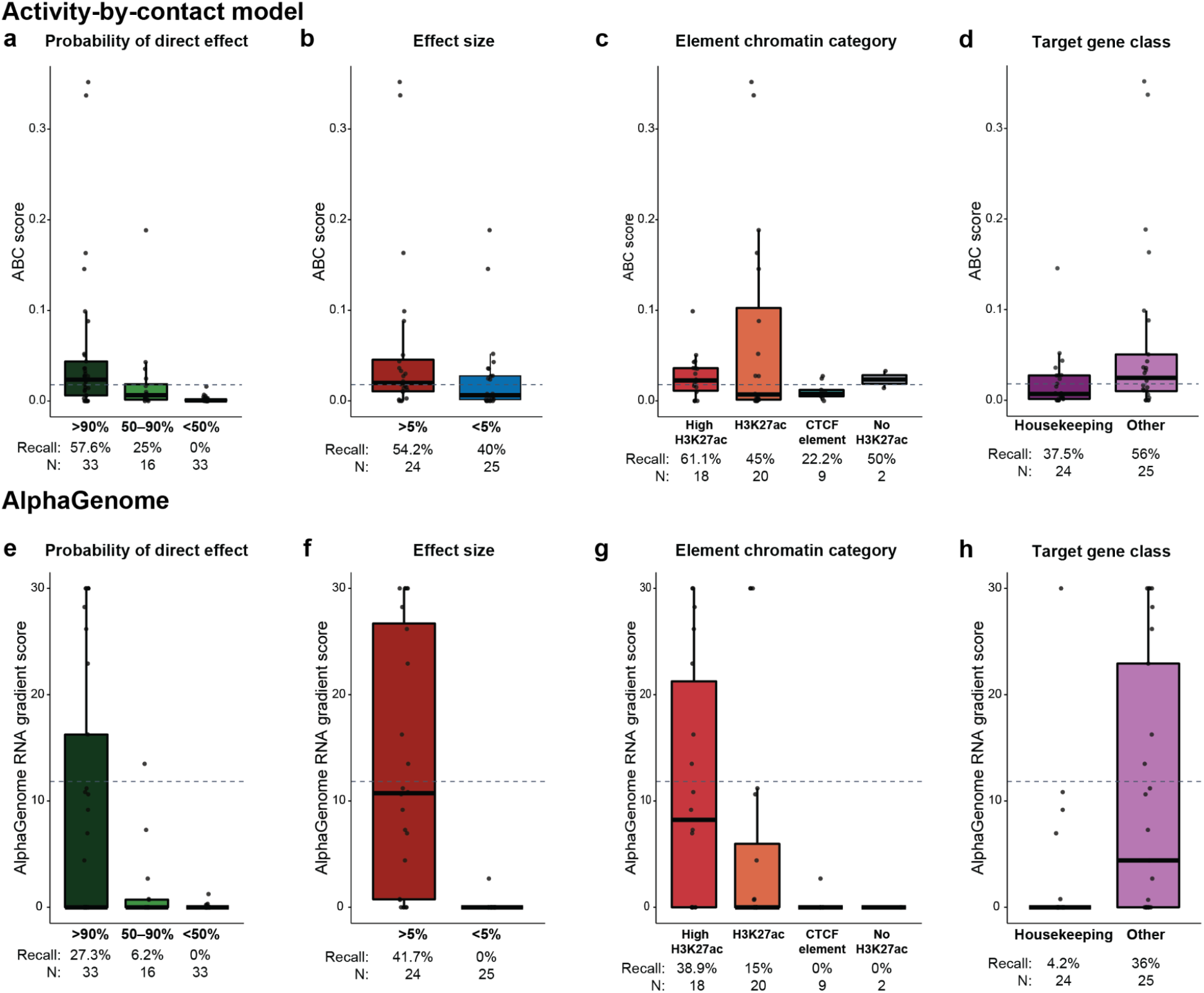
Sensitivity of additional predictive models for different categories of DC-TAP-seq DE-G pairs. **a.** ABC^16^ (A = DNase, C = Average Hi-C) scores for significant downregulated DE-G pairs from DC-TAP- seq, stratified by probability of direct effect. Recall is defined as the percent of pairs with scores above the model threshold. To annotate pairs with ABC scores, elements were expanded to 500 bp and merged if overlapping. Dashed grey line indicates the binary score threshold. **b.** ABC scores for significant downregulated DE-G pairs from DC-TAP-seq with a probability of direct effect greater than 50%, stratified by effect size. **c.** ABC scores for significant downregulated DE-G pairs from DC-TAP-seq with a probability of direct effect greater than 50%, stratified by chromatin category of the element. **d.** ABC scores for significant downregulated DE-G pairs from DC-TAP-seq with a probability of direct effect greater than 50%, stratified by class of the target gene. **e.** As **a**, but for AlphaGenome enhancer-gene linking scores computed using the RNA gradient method^46^. **f.** As **b**, but for AlphaGenome enhancer-gene linking scores. **g.** As **c**, but for AlphaGenome enhancer-gene linking scores. **h.** As **d**, but for AlphaGenome enhancer-gene linking scores.

**Extended Data Fig. 14.**
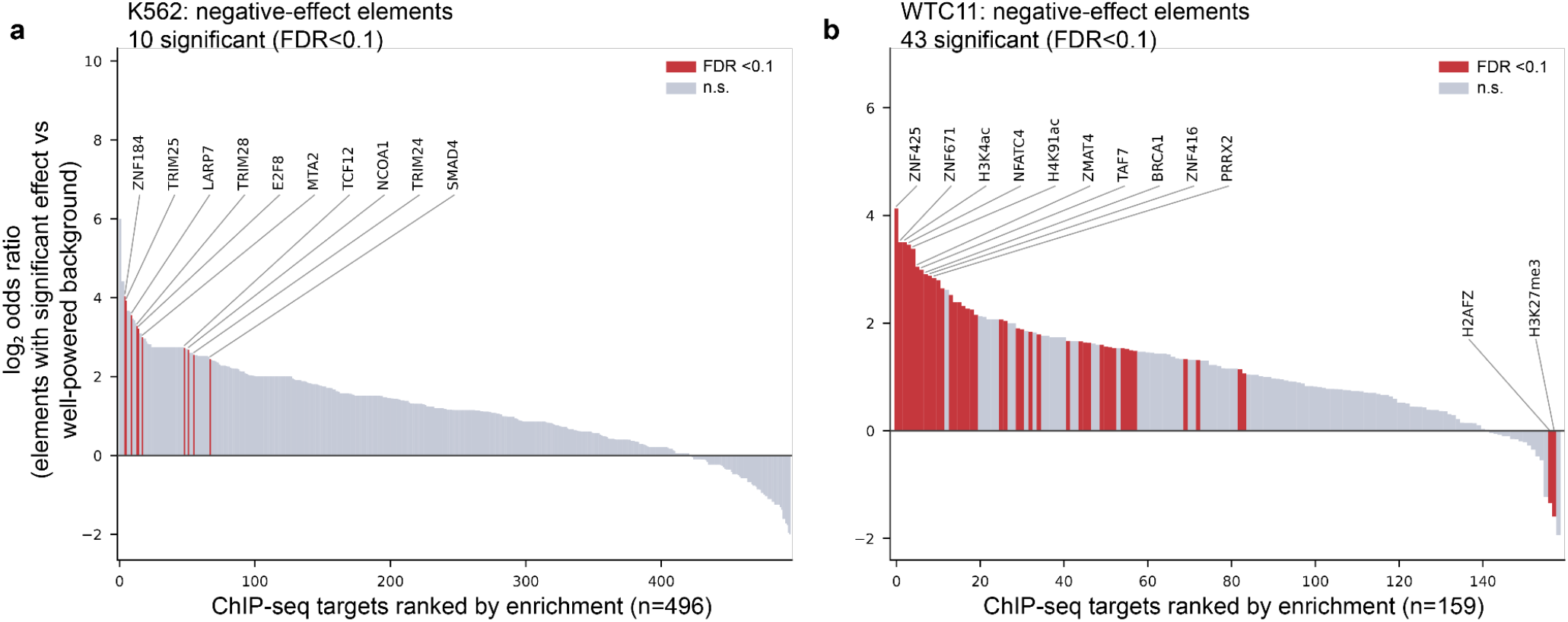
Comprehensive ChIP-seq overlap of DC-TAP-seq Des. Per-target enrichment of ENCODE ChIP-seq peak overlap among DC-TAP-seq elements with a significant negative effect on at least one gene, in (**a**) K562 and (**b**) WTC11. Each bar is one ChIP-seq target (transcription factor, other nuclear protein, or histone modification); bars are ordered left-to-right by decreasing enrichment. Enrichment was tested per target with a two-sided Fisher’s exact test on the 2×2 table of peak overlap versus non-overlap in positives (elements with a significant negative effect) versus well-powered negatives (no significant effect on any gene; >80% power to detect a 15% effect), using one representative experiment per target (**Methods**). Histone-mark peaks were extended ±175 bp; other peaks were used as called. The y-axis is the log₂ odds ratio (Haldane–Anscombe +0.5 continuity correction applied to the point estimate), with positive values indicating enrichment in positives relative to well-powered negatives. Bar color indicates significance after Benjamini–Hochberg FDR correction across all targets within each cell type (red, FDR < 0.1; grey, not significant); the horizontal line marks a log₂ odds ratio of 0. The ten most enriched significant targets are labeled above each panel; in (**b**), the two significantly depleted targets are also labeled. K562: 10 significant targets (all enriched); WTC11: 43 significant targets (41 enriched, 2 depleted).

**Extended Data Fig. 15.**
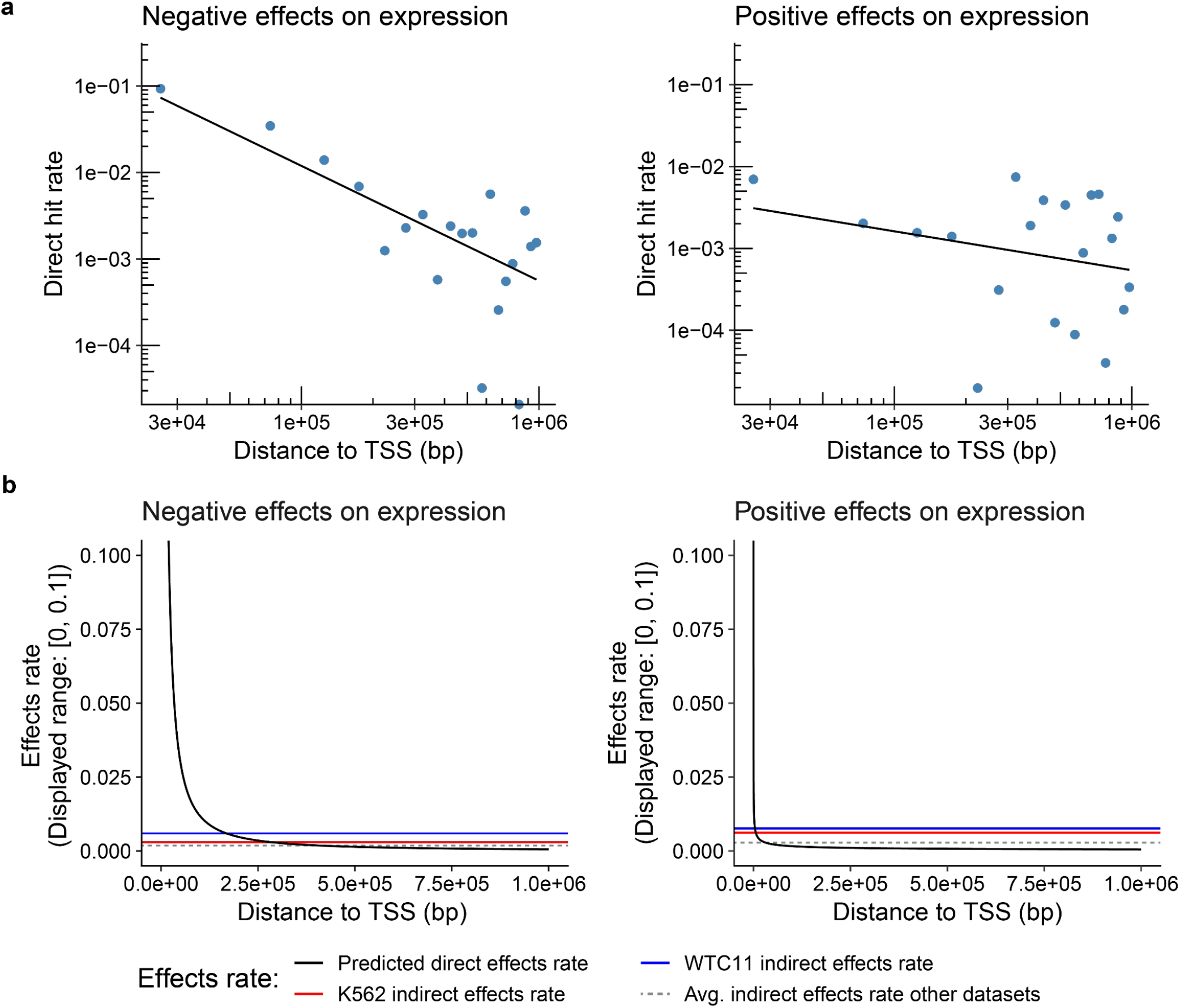
Modeling direct and indirect effect rates. **a.** Fitted power law function for direct hit rate of negative and positive effects on expression as a function of distance to target gene TSS. Plots show fit on direct hit rates calculated in 50kb windows and averaged across datasets used to fit power law function (Methods). **b.** Power law predicted direct effects rate as function of distance to TSS for negative and positive effects on expression (black lines) and indirect effect rates of negative and positive effects on expression for the K562, WTC11 and all other dataset used to model direct and indirect effect rates (Methods).

**Extended Data Fig. 16.**
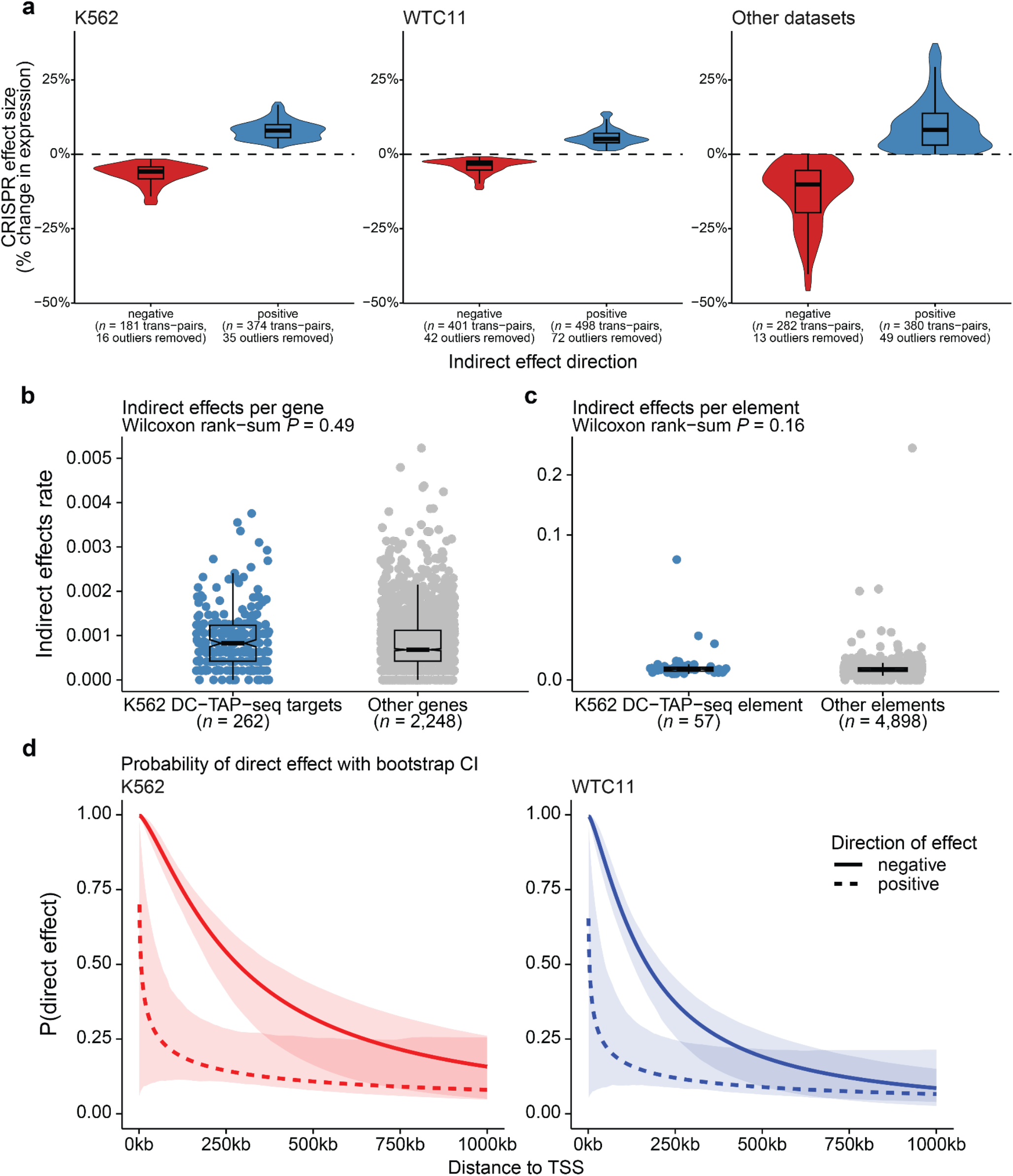
Modeling probability of direct effects. **a.** Percent change in expression effect size distributions of significant indirect (trans-acting) effects with negative and positive effects on expression used to calculate indirect effect rates in K562 and WTC11. Outliers were identified using Tukey’s method, defined as data points falling outside 1.5 times the interquartile range (IQR) below the first quartile (Q1) or above the third quartile (Q3). This calculation was done separately for each shown distribution and outliers were removed for visualization. **b.** Indirect effects rates per gene in the Gasperini *et al.*, 2019^14^ K562 enhancer perturbation dataset. All distal element perturbations were tested against all genes on other chromosomes than the perturbations to calculate indirect effect rates. Indirect rates are shown for genes part of the K562 DC-TAP-seq screen versus a random subset of other genes in the Gasperini dataset with TPM distribution matching the DC- TAP-seq genes. **c.** Indirect effects rates per element in the Gasperini *et al.*, 2019^14^ K562 enhancer perturbation dataset. All distal element perturbations were tested against all genes on other chromosomes than the perturbations to calculate indirect effect rates. Indirect rates are shown for distal elements also perturbed in the K562 DC-TAP-seq screen versus all distal elements in the Gasperini dataset. **d.** Modeled probability of direct effects as function of TSS for negative and positive effects on expression in the DC-TAP-seq datasets. 95% confidence intervals were computed using a cluster bootstrap approach, where for each iteration DE-G pairs from a set of randomly sampled perturbations were used in to calculate direct and indirect rates, fit a power law function to compute direct effect rates as function of distance to TSS, and calculate probability of direct effects (see Methods). Box plot elements in **a,b,c**: center line: median; box bounds: 25th and 75th percentile (interquartile range); whiskers extend to most extreme value within 1.5 x interquartile range from the box; outlying points not shown.

### Supplementary Tables

**Supplementary Table 1. List of guide pools used for pilot study**

Column names: name = name of the guide sequence; GuideSequenceMinusG = guide sequence without the 5’G; target_name = name of the target element; target_type = type of the element targeted by the guide (Enh = enhancer element, TSS = Transcription Start Site, negative_control = no site present in the genome, safe_targeting = non-functional genomic site); vector_type = vector expressing the guide RNA (sgOpti-CS = sgOpti plasmid with the capture sequence scaffold downstream of hU6 promoter, pBA904 = Addgene #122238 vector with capture sequence scaffold downstream of mU6 promoter, CROP-HyPR = CROP-seq vector with HyPR barcode); cell_type = cell type where the guides were used in.

**Supplementary Table 2. Genes and primers used in K562 pilot study**

Column names: gene = official gene symbol; outer_primer = outer tap seq primer; inner_primer = inner tap seq primer (no adapter); cell_type.

**Supplementary Table 3. Summary of all element-gene pairs tested in DC-TAP-seq**

Column name: intended_target_name_hg38 = The region of the targeted element lifted over from hg19 to hg38 in chr:start-end format;

targeting_chr_hg38 = The chromosome (chr) of intended_target_name_hg38;

targeting_start_hg38 = The start of intended_target_name_hg38;

targeting_end_hg38 = The end of intended_target_name_hg38;

intended_target_name_hg19 = The region of the targeted element in hg19 in chr:start-end format;

targeting_chr_hg19 = The chromosome (chr) of intended_target_name_hg19;

targeting_start_hg19 = The start of intended_target_name_hg19;

targeting_end_hg19 = The end of intended_target_name_hg19;

element_gene_pair_identifier_hg38 = Unique identifier for each element gene pair in gene_id|intended_target_name_hg38 format;

element_gene_pair_identifier_hg19 = Unique identifier for each element gene pair in gene_id|intended_target_name_hg19 format;

gene_symbol = Gene tested for differential expression upon CRISPRi perturbation of the targeted element in gene symbol format;

gene_id = Gene tested for differential expression upon CRISPRi perturbation of the targeted element in ensembl id format;

cell_type = Cell type tested;

fold_change_effect_size = The effect size of the tested element-gene pair in fold change;

log_2_FC_effect_size = The effect size of the tested element-gene pair in log 2 fold change;

pct_change_effect_size = The effect size of the tested element-gene pair in percent change; standard_error_fold_change = Standard error of the fold change;

standard_error_log_2_FC = Standard error of the log 2 fold change;

standard_error_pct_change = Standard error of the percent change;

lower_CI_95_fold_change = Lower bound of the 95% confidence interval for fold change effect size;

upper_CI_95_fold_change = Upper bound of the 95% confidence interval for fold change effect size;

lower_CI_95_log_2_FC = Lower bound of the 95% confidence interval for log 2 fold change effect size;

upper_CI_95_log_2_FC = Upper bound of the 95% confidence interval for log 2 fold change effect size;

lower_CI_95_pct_change = Lower bound of the 95% confidence interval for percent change effect size;

upper_CI_95_pct_change = Upper bound of the 95% confidence interval for percent change effect size;

sceptre_p_value = Raw p-value from SCEPTRE test for association between the element and gene;

sceptre_adj_p_value = Adjusted p-value after multiple hypothesis testing correction;

significant = Boolean indicator of whether the element-gene pair shows significant association at adjusted p-value of <10%;

distance_to_gencode_gene_TSS = Distance in base pairs from element to gene’s transcription start site (TSS) in GENCODE;

distance_to_abc_canonical_TSS = Distance in base pairs from element to gene’s transcription start site in ABC model;

chrTSS_hg38 = Chromosome of the gene’s transcription start site in hg38;

startTSS_hg38 = Start position of the gene’s transcription start site in hg38;

endTSS_hg38 = End position of the gene’s transcription start site in hg38;

element_location = Whether the element tested lies in a promoter or not (distal);

gencode_promoter_overlap = element overlap with protein-coding promoters in gencode;

abc_tss_overlap = element overlap with protein-coding promoters from ABC paper;

gencode_protein_coding_gene_body_overlap = element overlap with protein coding transcript gene bodies defined by GENCODE;

DistalElement_Gene = all high-confidence distal element-gene pairs created by removing any pair where the element overlaps a promoter or where the element is in the gene body of the gene it’s tested against or where the element is <1kb from the transcription start site of the gene it’s tested against; DistalPromoter_Gene = Pairs where the element overlaps the promoter of a gene different from the gene it’s tested against;

selfPromoter = Pairs where the element overlaps the promoter of their tested gene (promoter elements tested against their own gene);

Positive_Control_DistalElement_Gene = High-confidence distal element-gene pairs that are denoted as positive control distal elements in the screen design;

Positive_Control_selfPromoter = Pairs that are selfPromoter and which were originally designed to be TSS positive controls (tss_pos in design_file_type column);

Random_DistalElement_Gene = High-confidence distal element-gene pairs that are “unbiased” elements targeted in each of the 2.5 Mb loci;

power_at_effect_size_2 = Statistical power to detect an effect size of 2% change at adjusted p-value of <10%;

power_at_effect_size_3 = Statistical power to detect an effect size of 3% change at adjusted p-value of <10%;

power_at_effect_size_5 = Statistical power to detect an effect size of 5% change at adjusted p-value of <10%;

power_at_effect_size_10 = Statistical power to detect an effect size of 10% change at adjusted p-value of <10%;

power_at_effect_size_15 = Statistical power to detect an effect size of 15% change at adjusted p-value of <10%;

power_at_effect_size_20 = Statistical power to detect an effect size of 20% change at adjusted p-value of <10%;

power_at_effect_size_25 = Statistical power to detect an effect size of 25% change at adjusted p-value of <10%;

power_at_effect_size_50 = Statistical power to detect an effect size of 50% change at adjusted p-value of <10%;

intended_positive_control_distal_element_target_gene = the intended response gene for the positive control distal elements;

intended_positive_control_target_gene = the intended response gene for positive control distal elements (tss_pos and tss_random);

design_file_target_name = the original target name in the screen design;

design_file_type = the original target type in the screen design (tss_pos = promoter element of a positive control gene;

tss_random = promoter element of random gene in random loci;

enh = candidate cis-regulatory element; DE = positive control distal element);

guide_ids = list of the guide ids targeting the element;

include_in_fdr = indicator if the element-gene pair was included for FDR calculation without the positive controls, i.e. with the conditional (selfPromoter == FALSE & Positive_Control_DistalElement_Gene == FALSE & Positive_Control_selfPromoter == FALSE);

sceptre_adj_p_value_wo_pos_controls = Adjusted p-value after multiple hypothesis testing correction upon removing the positive controls;

significant_wo_pos_controls_20fdr = indicator if the element-gene pair shows significant association after removal of positive controls at adjusted p-value of <20%;

power_at_effect_size_2_wo_pos_controls_20fdr = Statistical power to detect an effect size of 2% change upon removal of positive controls at adjusted p-value of <20%;

power_at_effect_size_3_wo_pos_controls_20fdr = Statistical power to detect an effect size of 3% change upon removal of positive controls at adjusted p-value of <20%;

power_at_effect_size_5_wo_pos_controls_20fdr = Statistical power to detect an effect size of 5% change upon removal of positive controls at adjusted p-value of <20%;

power_at_effect_size_10_wo_pos_controls_20fdr = Statistical power to detect an effect size of 10% change upon removal of positive controls at adjusted p-value of <20%;

power_at_effect_size_15_wo_pos_controls_20fdr = Statistical power to detect an effect size of 15% change upon removal of positive controls at adjusted p-value of <20%;

power_at_effect_size_20_wo_pos_controls_20fdr = Statistical power to detect an effect size of 20% change upon removal of positive controls at adjusted p-value of <20%;

power_at_effect_size_25_wo_pos_controls_20fdr = Statistical power to detect an effect size of 25% change upon removal of positive controls at adjusted p-value of <20%;

power_at_effect_size_50_wo_pos_controls_20fdr = Statistical power to detect an effect size of 50% change upon removal of positive controls at adjusted p-value of <20%;

mean_sim_pert_cell = Mean number of cells perturbed between 100 replicates of power simulation; resized_merged_targeting_chr_hg38 = The chromosome of the resized_merged_element_gene_pair_identifier_hg38;

resized_merged_targeting_start_hg38 = The start of the resized_merged_element_gene_pair_identifier_hg38;

resized_merged_targeting_end_hg38 = The end of the resized_merged_element_gene_pair_identifier_hg38;

resized_merged_element_gene_pair_identifier_hg38 = Unique identifier for the resized and merged regions that were inputted into the chromatin categories pipeline in gene_symbol|chr:start-end format; ubiq_category = classification indicating whether the target gene is classified as ubiquitously and uniformly expressed (e.g., a housekeeping gene);

direct_rate_negative = Estimated rate of direct negative effects based on distance to TSS for tested element-gene pairs;

indirect_rate_negative = Estimated rate of indirect negative effects for the dataset to which this pair belongs;

direct_vs_indirect_negative = Probability that an observed negative effect size represents is the result of a direct cis-regulatory interaction;

direct_rate_positive = Estimated rate of direct positive effects based on distance to TSS for tested element-gene pairs;

indirect_rate_positive = Estimated rate of indirect positive effects for the dataset to which this pair belongs;

direct_vs_indirect_positive = Probability that an observed positive effect size represents is the result of a direct cis-regulatory interaction;

element_category = Classification of perturbed elements based on chromatin marks.

**Supplementary Table 4. Genes and primers used in the random screens**

Column names: gene = official gene symbol; ensemble_id = ensemble_id; tpm = tpm; type - whether it is an index gene, locus gene, positive control, negative control gene; outer_primer = outer tap seq primer; inner_primer = inner tap seq primer (no adapter); cell_type.

**Supplementary Table 5. Cell Ranger metrics for K562 and WTC11 screens**

Column names: sample = 10x lane number with cell type identifier; cells_recovered = number of cell recovered from the lane; mRNA reads = number of reads in the mRNA library; mean_mRNA_reads_per cell; reads_mapped_to_targeted_transcriptome; median_targeted_mRNA_UMI_per_cell; mRNA_sequencing_saturation; guide_reads = number of reads in the gRNA library; percent_guide_reads_with_protospacer; median_guide_UMIs_per_cell; guide sequencing saturation; cell_type

**Supplementary Table 6. Summary counts of tested pairs by category in the random screen**

Column names: Pair_Category = Type of element-gene pair (selfPromoter = Any promoter paired with the same gene, Positive_Control_selfPromoter = Positive control promoter paired with the same gene, DistalPromoter-gene = Any promoter paired with any gene but the same gene, DistalElement-gene = Any element which is not a promoter paired with a gene, Positive_Control_DistalElement_Gene = Positive control distal element paired with a gene, Random_DistalElement_Gene = Any distal element- gene pair that is not a positive control, Other_DistalElement-gene = Any DistalElement-gene that is not a Positive_Control_DistalElement_Gene or a Random_DistalElement_gene); Significant_Upregulated = pairs that were called significant and upregulated at adjusted p-value of <10%; Significant_Downregulated = pairs that were called significant and downregulated at adjusted p-value of <10%; Non_Significant_Well_Powered = pairs that were called non-significant at adjusted p-value of <10% but had ≥80% power to detect 15% effect size; Non_Significant_Under_Powered = pairs that were called non-significant at adjusted p-value of <10% and did not have ≥80% power to detect 15% effect size, Significant_Upregulated_wo_Positive_Controls_20FDR = pairs that were called significant and upregulated without positive controls at adjusted p-value of <20%; Significant_Downregulated_wo_Positive_Controls_20FDR = pairs that were called significant and downregulated without positive controls at adjusted p-value of <20%; Non_Significant_Well_Powered_wo_Positive_Controls_20FDR = pairs that were called non-significant without positive controls at adjusted p-value of <20%, but had ≥80% power to detect 15% effect size; Non_Significant_Under_Powered_wo_Positive_Controls_20FDR = pairs that were called non-significant without positive controls at adjusted p-value of <20%, and did not have ≥80% power to detect 15% effect size; Cell_Type = cell type (K562 or WTC11); Sum = sum of pairs in the columns Significant_Upregulated, Significant_Downregulated, Non_Significant_Well_Powered, Non_Significant_Under_Powered; Sum_wo_pos_ctrls = sum of pairs in the columns Significant_Upregulated_wo_Positive_Controls_20FDR, Significant_Downregulated_wo_Positive_Controls_20FDR, Non_Significant_Well_Powered_wo_Positive_Controls_20FDR, Non_Significant_Under_Powered_wo_Positive_Controls_20FDR.

**Supplementary Table 7. Metrics summary for datasets used in this study**

Column names: dataset_id = Identifier for the study/dataset; mean_cellsPerGrna = Average number of cells in which each guide RNA (gRNA) is detected; n_targeting_Grna = Total number of guide RNAs designed to target regulatory elements; n_Targets = Number of elements targeted in the dataset; mean_cellsPerTarget = Average number of cells captured per element; mean_umisPerCellPerGene = Average number of unique molecular identifiers (UMIs) per gene per cell; mean_Guide_UMIs_per_Cell_per_Guide = Average number of guide-specific UMIs detected per cell.

**Supplementary Table 8. Epigenetic ENCODE datasets used in this study**

Column names: Dataset_Name = The name or type of the genomics dataset; Experiment_Dataset_ID = The unique ENCODE experiment accession identifier; Lab = Depositor lab, Date_Released = The date when the dataset was made publicly available; Biosample = The biological sample used for the experiment including species and cell line or tissue type; Assay = The specific experimental technique used to generate the data; Target = The specific protein or histone modification target being studied; Nucleic_Acid_Type = The type of nucleic acid being sequenced either DNA or RNA; Accession_Data_File = The unique ENCODE file accession IDs for the processed data files, Filetype = The format of the data files.

**Supplementary Table 9. TPMs of WTC11 genes**

Column names: Gene_Id = Ensembl gene identifier; Chr = Chromosome name; Start = Start position of the gene on the chromosome (0-based); End = End position of the gene on the chromosome; Length = Total gene span (End - Start); Reads = Total number of reads mapped; TPM = Transcripts Per Million for the full gene; ExonLength = Total length of all exonic regions; ExonReads = Reads mapped to exonic regions; ExonTPM = TPM computed only from exonic reads; IntronLength = Total length of intronic regions; IntronReads = Reads mapped to intronic regions; IntronTPM = TPM computed only from intronic reads; UniqueLength = Length of gene regions that uniquely map to this gene; UniqueReads = Reads uniquely mapped to this gene; UniqueTPM = TPM based on uniquely mapped reads; UniqueExonLength = Length of exonic regions uniquely assigned to this gene; UniqueExonReads = Reads mapped to uniquely assigned exonic regions; UniqueExonTPM = TPM based on uniquely mapped exonic reads; UniqueIntronLength = Length of intronic regions uniquely assigned to this gene; UniqueIntronReads = Reads mapped to uniquely assigned intronic regions; UniqueIntronTPM = TPM based on uniquely mapped intronic reads.

**Supplementary Table 10. List of single gRNAs and primers used in RT-qPCR and CTCF guides used for validation**

Sheet qPCR_guide_primer, Column names: grna_id = Unique identifier for each guide RNA tested; grna_sequence = Sequence of the gRNA; target_name_hg38 = Name of the target element (based on hg38 genome build) that the gRNA is designed to target; type = Category of the target (e.g., TSS or Distal-element); Positive_control = Indicates whether the target is a positive control; readout_gene = gene that was readout in qPCR; Forward-primer = Forward primer sequence for PCR; Reverse-primer = Reverse primer sequence for PCR; cell_type.

Sheet CTCF_guides, Column names: grna_id = Unique identifier for each guide RNA tested; grna_sequence = Sequence of the gRNA; target_name_hg38 = Name of the target element (based on hg38 genome build) that the gRNA is designed to target.

